# An Atlas of Cells in the Human Tonsil

**DOI:** 10.1101/2022.06.24.497299

**Authors:** Ramon Massoni-Badosa, Paula Soler-Vila, Sergio Aguilar-Fernández, Juan C. Nieto, Marc Elosua-Bayes, Domenica Marchese, Marta Kulis, Amaia Vilas-Zornoza, Marco Matteo Bühler, Sonal Rashmi, Clara Alsinet, Ginevra Caratù, Catia Moutinho, Sara Ruiz, Patricia Lorden, Giulia Lunazzi, Dolors Colomer, Gerard Frigola, Will Blevins, Sara Palomino, David Gomez-Cabrero, Xabier Agirre, Marc A. Weniger, Federico Marini, Francisco Javier Cervera-Paz, Peter M. Baptista, Isabel Vilaseca, Felipe Prosper, Ralf Küppers, Ivo Glynne Gut, Elias Campo, José Ignacio Martin-Subero, Holger Heyn

## Abstract

Palatine tonsils are secondary lymphoid organs representing the first line of immunological defense against inhaled or ingested pathogens. Here, we present a comprehensive census of cell types forming the human tonsil by applying single-cell transcriptome, epigenome, proteome and adaptive immune repertoire sequencing as well as spatial transcriptomics, resulting in an atlas of >357,000 cells. We provide a glossary of 121 annotated cell types and states, and disentangle gene regulatory mechanisms that drive cells through specialized lineage trajectories. Exemplarily, we stratify multiple tonsil-resident myeloid slancyte subtypes, establish a distant *BCL6* superenhancer as locally active in both follicle-associated T and B cells, and describe SIX5 as a potentially novel transcriptional regulator of plasma cell maturation. Further, our atlas is a reference map to understand alterations observed in disease. Here, we discover immune-phenotype plasticity in tumoral cells and microenvironment shifts of mantle cell lymphomas (MCL). To facilitate such reference-based analysis, we develop HCATonsilData and SLOcatoR, a computational framework that provides programmatic and modular access to our dataset; and allows the straightforward annotation of future single-cell profiles from secondary lymphoid organs.

## Introduction

Secondary lymphoid organs (SLO) are essential to develop tolerance and adaptive immunity against self or foreign antigens, respectively. They include lymph nodes, spleen, Peyer’s patches, mucosa-associated lymphoid tissue (MALT) and the Waldeyer’s tonsillar ring (including pharyngeal, adenoids, tubal, lingual and palatine tonsils) (Ruddle and Akirav, 2009). Tonsils are under constant exposure to antigens via the upper respiratory tract which makes them a compelling organ to study the interplay between innate and adaptive immune cells during follicular and germinal center (GC) development to build adaptive immunity (Crotty, 2011a).

Tonsils have a non-keratinizing stratified squamous epithelium, organized into tubular, branched crypts that enlarge the tonsillar surface. Within the crypts, microfold cells (or M cells) sample antigens at their apical membrane. Subsequently, antigen presenting cells (APC), such as dendritic cells (DC), process and present antigens to T cells in the interfollicular or T cell zone. Alternatively, antigens are kept intact by follicular dendritic cells (FDC) in lymphoid follicles, where they are recognized by specialized B cells (Nave et al., 2001). Such recognition triggers the GC reaction, whereby activated naive B cells (NBC) give rise to GC B cells (GCBC) which undergo clonal selection, proliferation, somatic hypermutation, class switch recombination (CSR) and further mature into long-lived plasma cells (PC) or memory B cells (MBC) (De Silva and Klein, 2015). Hallmarks of tonsillar immune cells are their vast specialization and plasticity to differentiate into different functional states, tailoring immune responses to distinct insults. As an example, PC can produce different antibody isotypes, such as IgA and IgE, with specific effector functions to neutralize pathogens at mucosal surfaces or to respond against parasitic worms, respectively (Schroeder and Cavacini, 2010). Similarly, T helper subtypes (Th1, Th2 or Th17) can secrete distinct sets of cytokines (*i*.*e*., IFN-g, IL4, IL17) with markedly specific functions in the adaptive immune response and memory (Zhu et al., 2010). Thus, to comprehensively grasp the heterogeneity of tonsillar cells, a granular taxonomy of cell types and states is needed.

In this setting, the inherent discriminative power of single-cell RNA-sequencing (scRNA-seq) has catalyzed the creation of cellular taxonomies of lymphoid organs, such as the thymus (Park et al., 2020) and the bone marrow (Baccin et al., 2020). In the context of the Human Cell Atlas (HCA) (Regev et al., 2017), these taxonomies identified previously uncharacterized cell types and provided a reference to annotate cell types and states by training classifiers (Kang et al., 2021; Lotfollahi et al., 2022) and through curated cell ontologies (Osumi-Sutherland et al., 2021). While the transcriptome allows precise cellular phenotyping, recent atlases also incorporate additional layers, such as the epigenome or spatial profiles for mechanistic and structural insights, respectively. Together, such complementary modalities contribute multiple layers to define cell identities (Wagner et al., 2016). Previous single-cell profiling efforts of the human tonsil already provided insights into specific cell populations (*e*.*g*., B cells (King et al., 2021b, 2021a) or innate lymphoid cells, ILCs (Björklund et al., 2016)), but lacked cell numbers and multimodal information to fully capture the cellular complexity of the organ. Here, we generated a human tonsil atlas composed of >357,000 cells profiled across five different data modalities, such as transcriptome, epigenome, proteome, adaptive immune repertoire and spatial location. We identified 121 cell types and states, connected through a continuum of gene regulatory events, cell-cell communication and spatial colocalization to form the functional units of the human tonsil. To showcase the translational applicability of our reference for the analysis of pathological conditions, we projected mantle cell lymphoma (MCL) cells, an aggressive lymphoid neoplasm (Campo and Rule, 2015) that frequently presents in the tonsil (Tashakori et al., 2021), onto the atlas to elucidate transcriptional plasticity and compositional shifts in the tumor microenvironment (TME) of the organ.

## Results

### A single-cell multiomic atlas of human tonsillar cells

To create a comprehensive census of tonsillar cells, we sequenced the transcriptome of over 209,000 unselected cells from ten human tonsils by scRNA-seq. These tonsils covered three age groups: children (n=6, 3-5 years), young adults (n=3, 26-35 years) and old adults (n=1, 65 years) (Figure 1A). We complemented the transcriptional profiles with single-cell-resolved open chromatin landscapes (scATAC-seq and scRNA/ATAC-seq; *i*.*e*., Multiome) as well as protein (CITE-seq), adaptive repertoire (single-cell B and T cell receptor sequencing; *i*.*e*., scBCR/TCR-seq) and spatial transcriptomics (ST) profiles (Figure 1A; Tables S1 and S2). Initially, we created a high-level visualization and annotation of all cells across technologies by integrating high-quality transcriptome profiles from scRNA-seq and Multiome (Figure 1B). Our integration strategy (STAR Methods) removed most technical variability, confirmed by an increased local inverse simpson index (LISI) (Korsunsky et al., 2019) across confounders (Figures S1A and S1B), and preserved biological heterogeneity, highlighted by integrating an external, well-annotated dataset of ∼35,000 tonsillar cells (Figures S1A and S1C) (King et al., 2021a).

**Figure 1.**
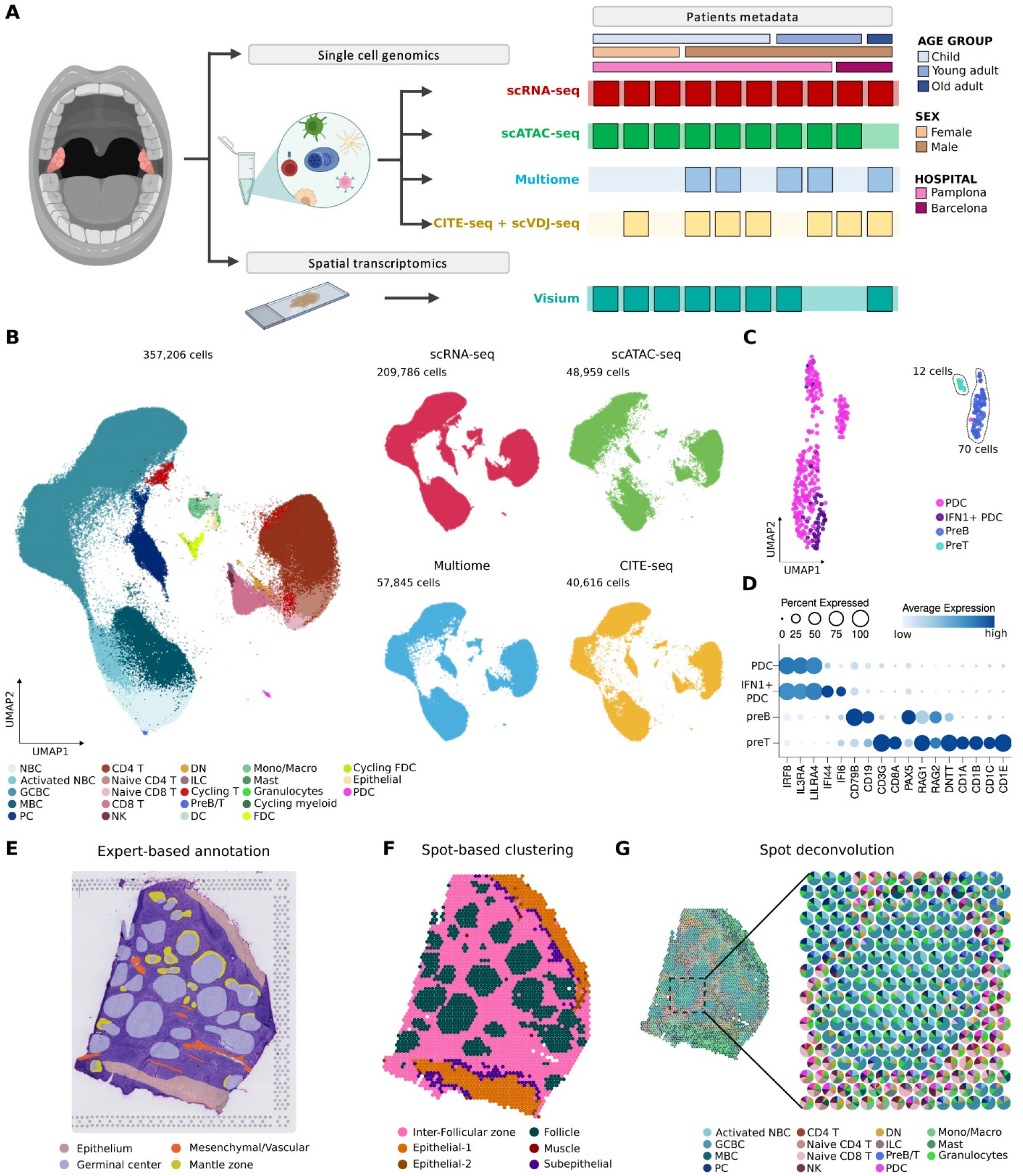
A single-cell multiomic atlas of human tonsillar cells. (**A**) Schematic diagram of the multiomic approach followed and boolean table annotated with patient metadata (age group, sex and hospital) showing data modalities performed for each sample. (**B**) UMAP projection of the 357,206 tonsillar cells analyzed. *Left*, colored by the main 23 populations. *Right*, split per data modality. (**C)** UMAP projection of tonsillar plasmacytoid dendritic cells (PDC) and precursor B and T cells (preB/T clusters). (**D)** Dotplot showing average gene expression for the top 17 genes *(x-axis)* per cluster *(y-axis)*. Dot size reflects the percentage of expressing cells and the color shows the average expression level. (**E**) Representative histologically annotated slide of a human tonsil. (**F**) Gene expression-based clusters of ST spots on a representative slide. (**G**) Spatial scatter pie plot showing each ST spot as a pie chart representing the predicted proportion of cell types for a representative slide.

Following Louvain clustering, we first assigned cells into nine broad groups: (1) NBC and MBC, (2) GC B cells (GCBC), (3) PC, (4) CD4 T cells, (5) cytotoxic cells [CD8 T cells, NK, ILC, double negative T cells (DN)], (6) myeloid cells (DC, macrophages, monocytes, granulocytes, mast cells), FDC, (8) epithelial cells and (9) plasmacytoid dendritic cells (PDC) (Figure 1B; Figure S1D). As observed in previous studies, naive CD8 T cells shared similar transcriptomes with naive CD4 T cells (Hao et al., 2021) and NBCs with MBCs (Figure 1B) (Vilarrasa-Blasi et al., 2021). In addition, we observed subsets of proliferative cells in several clusters (Figure 1B; Figure S1E). Inside the PDC cluster, we identified two intriguing additional clusters of precursor T cells (preT; *CD3G, CD8A*) and precursor B cells (preB; *CD19, CD79B, PAX5*), likely because PDC develop from common lymphoid progenitors (Figures 1C and 1D; Table S3) (Dress et al., 2019). Both preT and preB cells expressed members of the VDJ recombinase, including *RAG1, RAG2*, and *DNTT* (TdT), supporting T and B cell development within human tonsils (Figure 1D) (McClory et al., 2012; Strauchen and Miller, 2003). PreT cells further expressed several components of the CD1 family of MHC class I-like genes (Figure 1D; Table S3). PreB and preT clusters were composed of only 70 (0.033%) and 12 cells (0.0057%), respectively, highlighting the high discriminatory power of our atlas.

Subsequently, we followed a recursive, top-down, clustering approach (STAR Methods), resulting in a total of 121 clusters, which we thoroughly annotated using the complementary information across all profiling layers. To generate integrated multimodal profiles, we transferred the cluster labels and UMAP coordinates learnt from scRNA-seq and Multiome data to scATAC-seq and CITE-seq data, respectively (STAR Methods; Figure 1B). Notably, such integration corrected batch effects and yielded high cell type prediction probabilities (Figure S1F-S1H). As both cellular composition and position are defining the functionality of an organ, we further integrated single-cell with ST profiles. Broadly, spot-based clustering, expert annotation and spot deconvolution identified the main histological areas of human tonsils: epithelium, B cell follicles, and T cell/interfollicular zones (Figures 1E and 1F; Figure S2A), and formed the basis for subsequent spatial mapping of tonsillar cell types (Figure 1G; Figure S2B). Together, our tonsil atlas resource includes 357,206 cells (209,786 scRNA-seq, 48,959 scATAC-seq, 57,845 Multiome, 40,616 CITE-seq; Figure 1B) and 16,224 spatial transcriptomics spots.

### CD4 T follicular and non-follicular cell fate decision in the human tonsil

CD4 T follicular helper cells (Tfh) are the specialized providers of B cell help for the formation of GC in tonsils. Tfh cell differentiation depends on the expression of the master regulator transcription factor (TF) BCL6 and the absence of PRDM1. Tfh specification is a multifactorial and multistage process that begins with the activation of naive CD4 T cells receiving signals provided by antigen-presenting DCs (Choi and Crotty, 2021). We identified a naive CD4 subpopulation with canonical naive CD4 T cell markers (*LEF1, CCR7* and CD62L+, CD45RA+; Figures 2A-2C; Table S4 and S5). Following activation, CD4 T cells differentiate into central memory (CM) CD4 T cells (decreased *CCR7* expression and CD45RO+, CD127+), here identified as two clearly distinct subpopulations differing in the upregulation of *IL7R* (CD127+; Figures 2A-2C; Table S5). Intriguingly, one of these populations showed higher expression of follicular marker genes (*e*.*g. IL6ST*), providing signals for early Tfh differentiation (CM pre-Tfh cells; Figure 2B; Figure S3A; Table S5) (Cano-Gamez et al., 2020). Conversely, CM pre-non-Tfh cells showed lower expression of follicular markers and increased *ANXA1, S100A4* and *ITGB1* levels, classical markers of CM T cells (Figure 2B; Figure S3A; Table S5) (Cano-Gamez et al., 2020).

**Figure 2.**
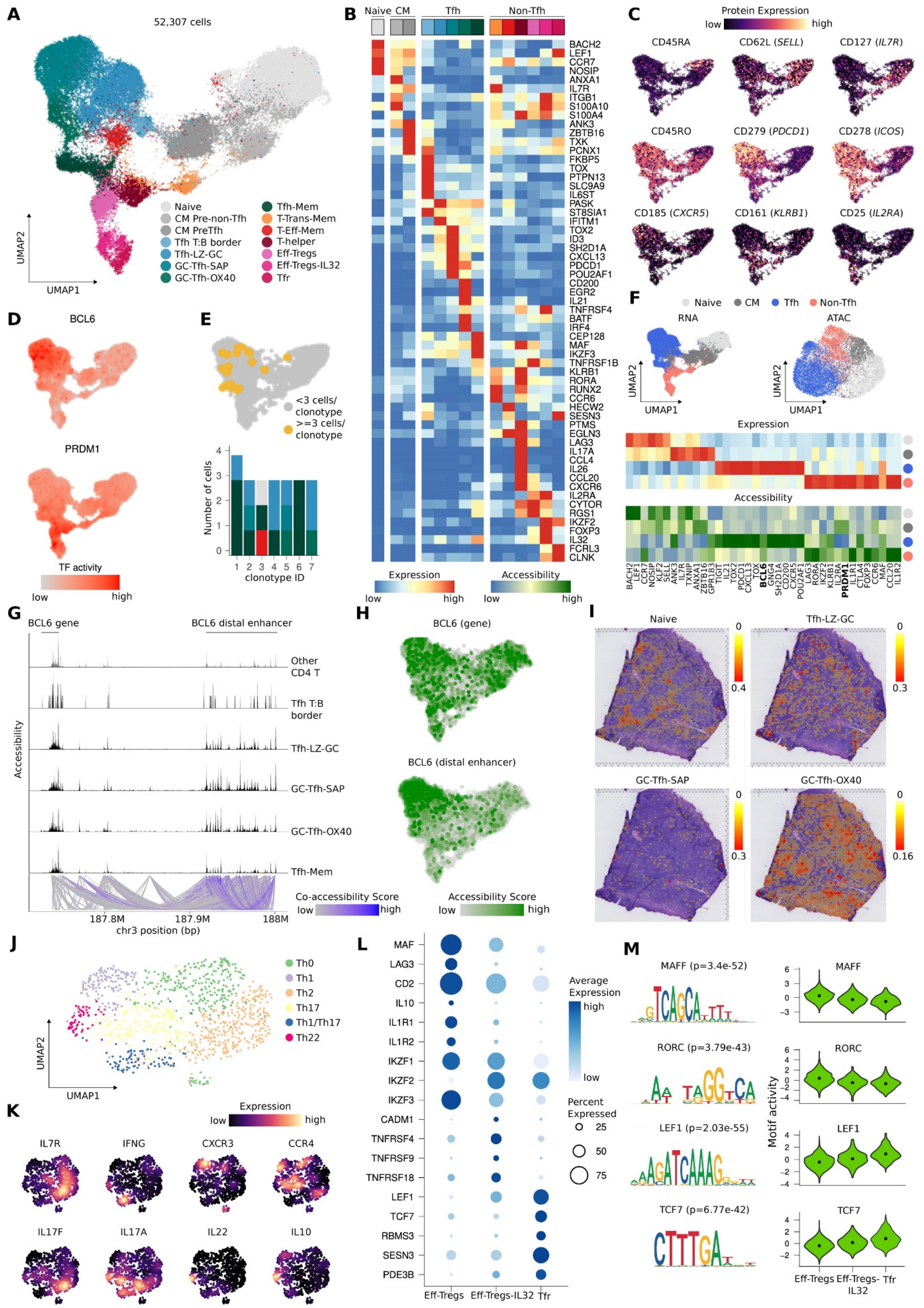
CD4 T follicular and non-follicular cell fate decision in the human tonsil. (**A**) UMAP projection of tonsillar CD4 T cells colored by scRNA-seq clusters. (**B**) Heatmap showing scaled mean marker expression by subpopulation. (**C**) UMAPs projection colored by protein expression of canonical phenotype markers of CD4 T cells. (**D**) UMAP projection highlighting the activity (AUCell score) of BCL6 and PRDM1 transcription factors (TFs). (**E**) Clonal expansion and diversity analysis in CD4 T cells. *Top*: UMAP projection showing clonal expansion denoted by >= 3 cells having identical CDR3 sequence (yellow). *Bottom*: Barplot of CD4 T cell subpopulations distribution across the top 7 most expanded clonotypes (*x-axis*) and the number of expanded cells (*y-axis*). The color code refers to the UMAP in panel A. (**F**) CD4 T main groups analysis: Naive; Central Memory (CM); T follicular helper (Tfh); Non T follicular helper (Non-Tfh). *Top*: UMAP projection of scRNA-seq (*left*) and scATAC-seq (*right*) cells colored by the CD4 T main groups. *Bottom*: Heatmap showing scaled (top) mean RNA gene expression and (bottom) mean accessibility values (all peaks in gene body + 2kb upstream) of key markers for the CD4 T main subpopulations. (**G**) Genome browser visualization of combined DNA accessibility and co-accessible links predicted by Cicero at the *BCL6* locus and nearby linked region. (**H**) UMAP projection showing the accessibility score of *BCL6* gene (gene body + 2kb upstream) (*top*) and the identified distal enhancer (*bottom*). (**I**) Predicted proportions of CD4 T subpopulations of interest analyzed with SPOTlight. We deconvoluted spots using CD4 T cell subpopulation annotations along with the general annotation of other cell types ensuring we captured the biological signal of all our cell types on a representative slide. (**J**) UMAP projection of tonsillar T-helper (Th) cells colored by the six Th scRNA-seq clusters. (**K**) UMAPs projection highlighting the estimated Nebulosa density expression (STAR Methods) for key interleukin and chemokine receptors. (**L**) Dotplot showing average expression for the top 18 genes *(y-axis)* for Treg subpopulations *(x-axis*). Dot size reflects the percentage of cells in a cluster expressing each gene and the color the average expression level. (**M**) *Left*: DNA sequence motifs for overrepresented motifs in Treg cells. *Right*: Violin plots of the motif activity score computed with ChromVar for the top four TF motifs in Eff-Tregs, Eff-Tregs-IL32 and Tfr.

We then aimed to classify the remaining CD4 T cell clusters into Tfh or non-Tfh cells based on the expression of *BCL6* and *PRDM1*. Because these TFs are expressed at low levels and affected by technical dropout events (Iacono et al., 2019), we inferred their activity using gene regulatory network (GRN) modeling (Van de Sande et al., 2020) (Tables S6 and S7) and identified five Tfh and six non-Tfh cell clusters (Figures 2D-2F). In line with this assignment, scTCR-seq analysis revealed an increased clonal expansion of Tfh cells (Figure 2E). At the epigenetic level, we observed general concordance between increased gene expression and open chromatin of key cell fate regulators, such as *PRDM1* (Figure 2F). However, the *BCL6* locus was invariably accessible in both Tfh and non-Tfh clusters pointing to alternative mechanisms driving its expression (Figure 2F). Then, we focused on the cis-regulatory landscape of *BCL6* and putative enhancer interactions for each cluster. We identified a strong connection between the *BCL6* locus and a distant superenhancer region exclusively in the Tfh-specific chromatin profile (Figure 2G). This finding was further confirmed by plotting the accessibility signature derived from open chromatin peaks specific to the *BCL6* gene and the cis-regulatory region, being only the latter enriched in terminally differentiated Tfh cells (Figure 2H; Figure S3B). Most interestingly, this Tfh-specific cis-regulatory region has been previous described to control *BCL6* expression in GCBC (Bunting et al., 2016), suggesting that this superenhancer is a master regulator in GC function across T and B cell lineages.

Following the initial priming of naive CD4 T cells by DC (Goenka et al., 2011) and upregulation of BCL6 (Choi et al., 2011), the expression of the chemokine receptor *CXCR5* is induced, while *CCR7* is repressed. *CXCR5* expression triggers the migration of pre-Tfh cells to the border of B cell follicles where they further differentiate (Weinstein et al., 2016). In line, we identified a CD4+ cluster with downregulation of *CCR7* and increased expression of *CXCR5* and *TOX*, the latter pointing to an early Tfh differentiation and migratory program to the border and primed interaction with B cells via ICOS-ICOSL (Tfh T:B border) (Figures 2A and 2B; Figure S3C). Following the Tfh migration trajectory, we identified a Tfh Light Zone GC (Tfh-LZ-GC) cluster with early signs of GC Tfh differentiation. Tfh-LZ-GC cells upregulated *IL21*, a well-described inducer for early GC Tfh differentiation (Figures 2A and 2B; Figure S3C) (Vinuesa et al., 2016). We further identified two clusters of terminal state differentiation and polarization of CD4 Tfh cells (e.g. *PDCD1, IL21*; Figure 2A-2C, Figure S3C), which differed in specific terminal differentiation programs. Specifically, the high expression of *SH2D1A* (SAP) identified one subpopulation as potent GC B cell state inducers by providing signals of adhesion and survival (GC Tfh-SAP; Figure 2B). *SH2D1A* is essential for CD4 T cell GC responses and the generation of memory B cells and long-lived plasma cells (Vinuesa et al., 2016). In the absence of *SH2D1A*, Tfh cells have impaired adhesion to GCBCs and fail to be retained in GCs as observed in X-linked lymphoproliferative disease (XLP), where genetic mutations results in a complete loss of GC Tfh and GC B cells (Booth et al., 2011). In contrast, the second terminally differentiated CD4 Tfh GC cluster was characterized by the expression of *TNFRSF4* (OX40; GC Tfh-OX40; Figures 2A and 2B). This population interacts via OX40-OX40L with B cells in the GC, where engagement of OX40L on activated B cells was reported to enhance immunoglobulin (Ig) secretion by PC for an improved maturation process (Fu et al., 2021). Finally, we observed a cluster of Tfh memory cells, a controversial subtype of follicular T cells (Figures 2A-2C) (Crotty, 2014). Tfh memory cells retained stable expression of *PDCD1, MAF, CXCR5* and upregulated *KLRB1*, preserving highly functional follicular characteristics (Figures 2B and 2C; Figure S3C). The memory phenotype was confirmed at protein level with the higher expression of CD45RO and CD161 (*KLRB1*) as well as by retaining the protein expression of PD-1 and ICOS (Figure 2C). This molecular setup provides further support for the capacity of Tfh memory cells to reenter the Tfh differentiation process upon activation and to become CD4 Tfh GC cells.

For the spatial mapping of the Tfh multistage differentiation process, we integrated scRNA-seq and ST data to deconvolute ST capture spots and to predict cell type locations (Figure 2I; Figure S4) (Elosua-Bayes et al., 2021). Naive T cells surround the follicles forming the T cell zone (Figure 2I), whereas Tfh T:B-border cells localize at the edge of follicles, where they interact with B cells (Figure S4A). Entering the follicles, Tfh-LZ-GC cells spatially mapped to the light zone (Figure 2I) and we observed enrichment of GC Tfh-SAP and GC Tfh-OX40 in the GCs (Figure 2I). Tfh-Mem cells were preferentially localized outside of the follicles, in line with the concept that Tfh cells acquire a less polarized Tfh phenotype and develop into memory Tfh cells upon leaving the GC (Figure S4A).

CD4 T cell fate decision is determined within the first few rounds of cell division to commit either towards Tfh or non-Tfh differentiation programs (Choi et al., 2011). An upregulation of chemokine receptors drives non-Tfh cells to exit the lymphoid tissue and to traffic to primary infection or inflammation sites (Mahnke et al., 2013). We identified six clusters of non-Tfh cells, derived from primed CM pre-non-Tfh cells, that expressed *PRDM1* and respective regulatory activity (Figures S3A and S3D). Compared to naive T cells, CM pre-non-Tfh cells expressed higher levels of CD28 and CD29 (Figure S3E). Such subpopulation has recently been described as a precursor of cytotoxic CD4 T cells in blood, tumor, and after viral infection. Accordingly, *ITGB1* (CD29) high CD4 T cells produce proinflammatory cytokines and granzymes and have superior cytolytic capacity (Nicolet et al., 2021). In the subsequent differentiation towards effector-memory T cells (T-Eff-Mem), a linear progression along a continuum of states is expected, initiated by transitional memory cells (T-Trans-Mem). T-Trans-Mem cluster markers (upregulation of *IL7R* and downregulation of *CCR7* and CD45RA; Figures 2A-2C; Figure S3C) supported an intermediate state between CM pre-non-Tfh cells and fully differentiated T-Eff-Mem cells. The further differentiated clusters split into T-Eff-Mem (*KLRB1*, CD45RO+ and CD161+) and different CD4 T-helper cell types (Figures 2A-2C). To identify the entirety of T-helper types in tonsils, we reclustered these cells and inferred their interleukin (IL) and chemokine receptor activity to guide their assignment to Th_0_ (*IL7R, IFNG, IL17A, GATA3*), Th_1_ (*CXCR3, TBX21, GZMK, CCR5*), Th_2_ (*CCR4, GATA3, IL17RB, IL4IL*), Th_17_ (*CCR6, RORC, IL17A, IL17F*), Th_17_/Th_1_ (*IFNG, IL26, IL17A*) and Th_22_ (*IL22, IL10, TNF*) cells (Figures 2J and 2K; Figure S3F; Table S8).

CD4 T regulatory (Treg) cells are vital for the maintenance of immune tolerance and homeostasis in GCs. In SLO, Tregs are a subset of CD4 T cells with a critical role in controlling antibody responses (Sage and Sharpe, 2016). We identified three subtypes of Tregs in tonsils (Figures 2A). We annotated effector Tregs (Eff-Tregs) based on gene expression (*IL2RA, CTLA4, IL1R1, IL1R2* and *IL10*), TF (*FOXP3, MAF, IKZF1* and *IKZF3*) and protein levels (CD25+ and CD28+; Figures 2B and 2C; Figure 2L; Figure S3C and S3E; Table S9). GRN analysis validated the increased activity of *PRDM1* and *FOXP3* in Eff-Tregs (Figure S3G). In line, scATAC-seq revealed an increased TF motif activity of RORC and MAF families in Eff-Tregs (Figure 2M; Table S9) (Wheaton et al., 2017). Further, we observed a second effector Treg population, positive for *FOXP3* and *IKZF1*, a positive regulator for *FOXP3* during Treg-polarization (Figure 2B and 2L) (Powell et al., 2019). In addition, this subpopulation expressed higher levels of *IL32* (Eff-Tregs-IL32), a proinflammatory molecule previously linked to Treg function in tumor-infiltrating Tregs to suppress antitumor responses (Figure 2B) (Galván-Peña et al., 2021). The third Treg subpopulation, identified as T follicular regulatory cells (Tfr), downregulated *IKZF1, FOXP3, IL2RA* (CD25) and *PRDM1* (Figure 2B and 2L; Figure S3C; Table S9). Tfr cells further presented increased naive markers, in concordance with an increased TF motif activity of LEF1 and TCF7 (Figure 2L and 2M). *TCF7* and *LEF1* are essential for Tfr development in mice (Yang et al., 2019). Intriguingly, the top marker *FCRL3* can bind secretory IgA to suppress the Tfr inhibitory function. This mechanism could be therapeutically targeted to modulate the regulatory activity in both malignancies and autoimmune diseases (Agarwal et al., 2020). To further validate effector and follicular signatures in all subtypes of Tregs, we scored previous Treg gene sets and observed an increased effector and follicular signature in the respective Treg subpopulations (Figure S3H) (Wing et al., 2017).

### Landscape of CD8 and Innate lymphoid cells in the human tonsil

CD8 T cells are cytotoxic, effector and memory cells that respond to different pathogens and control infections (Martin and Badovinac, 2018). In tonsils, we identified a large CD8 T naive subpopulation with canonical transcript (*LEF1* and *NOSIP*) and protein (CD45RA+) markers (Figure 3A-3C; Figure S5A; Table S10). After antigen encounter, naive cells initiate a program of effector differentiation and a subsequent formation of memory states. These recently formed memory populations are organized in a differentiation hierarchy, originating from stem-cell memory T cells (SCM CD8 T; *TCF7* and CD95+; Figure 3A; Figure S5; Table S10). SCM CD8 T cells self-renew and generate long-lived central memory T cells (CM CD8 T) (Gattinoni et al., 2011). CM CD8 T are considered to be multipotent with a subset displaying increased expansion potential upon antigen re-encounter (*IL7R, ANXA1* and CD45RO+; Figure 3A-3C; Table S10). Subsequently, we identified two distinct clusters of resident memory (RM) CD8 T cells (*CD69, EOMES*; Figure 3A and 3B; Table S10). One subpopulation differentially expressed higher levels of the activation markers *HLA-DRB1* and CD69 (RM CD8 activated T cells; Figure 3A; Figure S5; Table S10). The second RM population expressed CD103, the main markers of retention of resident cells in tissues (Figure 3A and 3C; Table S10) (Kok et al., 2022).

**Figure 3.**
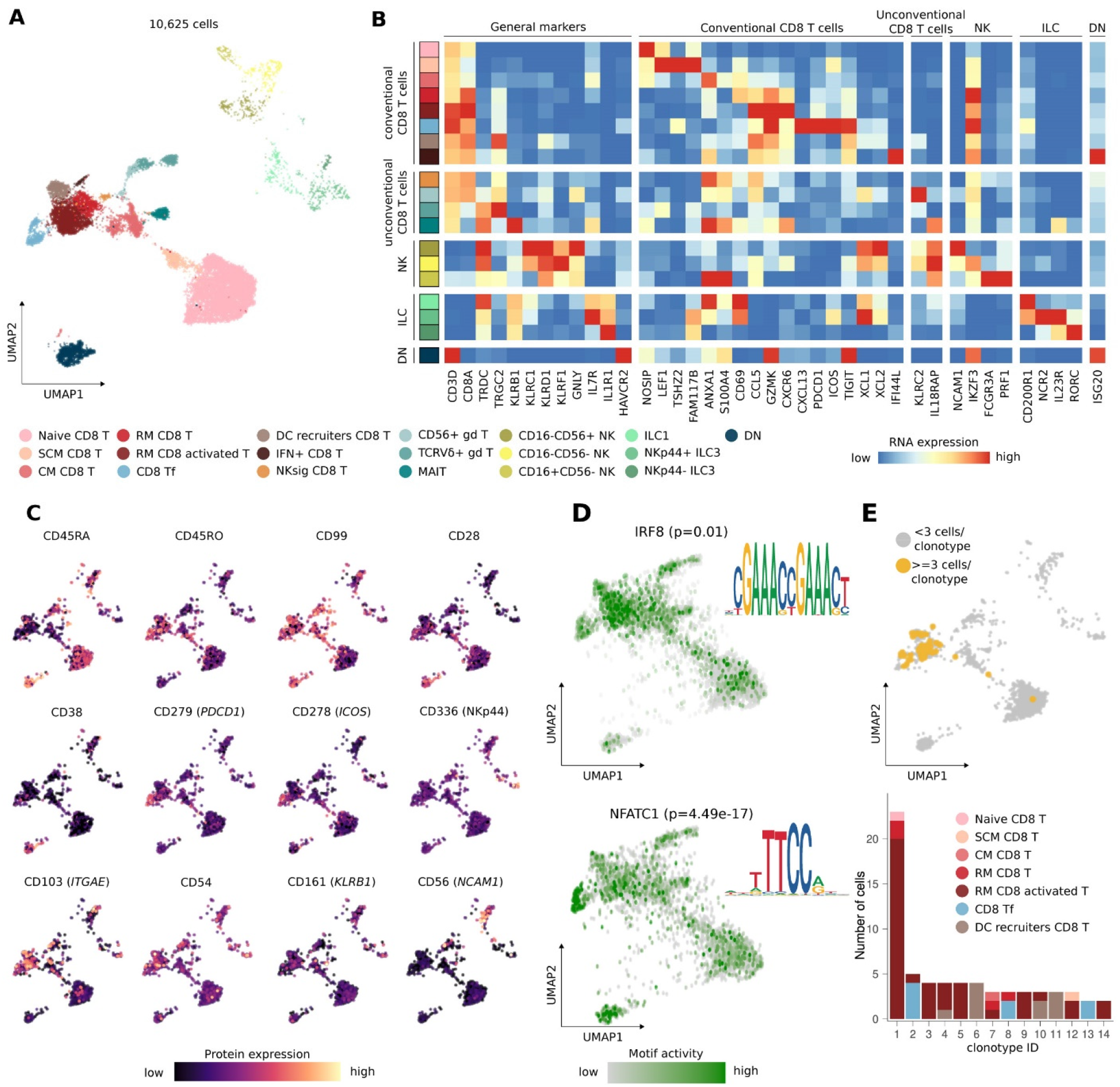
Landscape of CD8 and innate lymphoid cells (ILC) in the human tonsil. (**A**) UMAP projection of tonsillar CD8 T cells and ILC colored by scRNA-seq clusters. (**B**) Heatmap showing scaled mean marker expression per subpopulation. (**C**) UMAP projections highlighting the protein expression of the canonical phenotype markers of CD8 T and ILC. (**D**) UMAP projection highlighting the motif activity of IRF8 and NFATC1. The p-value represents the significance of the pairwise differential motif analysis performed for each TF. DNA sequence motifs logos for each TF. (**E**) Clonal expansion and diversity analysis in CD8 T cells. *Top*: UMAP projection showing clonal expansion denoted by >=3 cells having identical CDR3 sequence (yellow). *Bottom*: Barplot of CD8 T cell subpopulations distribution across the top 14 most expanded clonotypes (*x-axis*) and the number of expanded cells (*y-axis*).

Next, we identified a cluster of CD8 T cells with a follicular lineage (CD8 Tf; *e*.*g. CXCL13, ICOS, PDCD1* and PD-1+, ICOS+; Figure 3A-3C; Table S10). CD8 Tf cells can control B cell responses in GC and the production of autoantibodies (Chen et al., 2019). Pairwise differential motif activity analysis between CD8 naive T and RM CD8 T cells or CD8 Tf cells revealed NFATC-family TF motifs to be enriched in open chromatin peaks specific to CD8 Tf (Figure 3D; Table S11). NFAT TFs can promote the generation of follicular helper T cells in response to acute viral infections (Martinez et al., 2016). Conversely, IRF8, IRF9 and IRF7 motifs were specifically enriched in RM CD8 T cells, which play a key role in cytotoxic adaptive immune responses (Figure 3D; Table S11). Particularly, IRF8 is an important integrator of TCR costimulation and signaling, contributing to the phenotypic maturation and functionality of CD8 effector T cells (Miyagawa et al., 2012). RM CD8 activated T cells were the most clonally expanded CD8 T cell subset, as revealed by scTCR-seq (Figure 3E). Finally, CD8 T cells can also instruct pDC recruitment via CCL3 and CCL4 and attract the lymph node-resident XCR1 chemokine receptor-expressing DCs via secretion of XCL1 (Brewitz et al., 2017). We identified a CD8 T cell cluster expressing *CCL4, XCL1* and CD99, pointing to an interaction between these CD8 and DC subsets (Figure 3A-3C; Figure S5A; Table S10). CD99 is upregulated during DC and T cell interaction and plays an important role in subsequent T cell activation (Takheaw et al., 2019).

One of the main roles of CD8 cells is to control viral infections. We found a yet unreported CD8 population in tonsils with a NK-signature on both transcript (*e*.*g. GNLY, FCGR3A*) and protein level (CD16, CD122; Figure 3A and 3B; Figure S5B; Table S10). In addition, we found two different subsets of g/d T cells (Figure 3A; Table S10). One subset was identified as common g/d T cells (*TRGC2, TRDC*; Figure 3B). The second subset represented a unique subtype [*NCAM1* (CD56), RNA and protein] previously only reported in peripheral blood from pregnant women but not yet in tonsils (Figure 3B and 3C) (Nörenberg et al., 2021). Of note, CD56+ g/d T cells expressed markers of tissue-residency (*ITGAE* and CD69+; Figure S5A and S5B). We further annotated mucosal-associated invariant T (MAIT) cells and a cluster of DN CD8/CD4 T cells with a profile of proinflammatory activation (Figure 3A-3C; Table S10). DN cells have a role in inflammation and virus infection and proinflammatory DN T cells can attack healthy tissue in certain autoimmune diseases (Wu et al., 2022).

NK and ILC differed in their expression of *KLRF1* and *IL7R* (CD127), respectively (Figure 3A and 3B; Figure S5B). NK cells followed a well-described differentiation path (Freud et al., 2006; Pfefferle et al., 2019), which was guided by the reciprocal expression of CD16 (*FCGR3A*) and CD56 (*NCAM1;* Figure 3A-3C). The trajectory initiated from a CD16-CD56-NK precursor (*SELL*), followed by an intermediate state of CD16-CD56+ a highly cytotoxic (*GZMH, PRF1*), and a CD16+CD56-state (e.g. *CX3CR1*, and *TBX21*; Figure 3B; Figure S5A). CD16-CD56+ precursor NK cells displayed an intermediate *IL7R* expression compared to other NK populations and ILCs (Figure 3B). ILC1 cells could be distinguished from CD16-CD56+ precursor NK cells through their higher expression of *CD200R1* (Figure 3B) (Colonna, 2018). Two remaining ILC clusters could be annotated as NKp44+ ILC3 and NKp44-ILC3, based on the expression of *RORC, IL4I1, IL23R, IL1R1, ICOS, NCR2* (NKp44) and *AHR* (Figure 3B; Figure S5A), in line with the most recent classification of the International Union of Immunological Societies (IUIS) (Vivier et al., 2018). CITE-seq further revealed the specific protein expression of Nkp44 in Nkp44+ ILC3 cells (Figure 3C).

### B cell activation and GC dynamics

The tonsil is an important niche where key B cell maturation steps take place. NBCs become activated upon first antigen recognition to generate GCBCs that progress from the highly proliferative dark zone (DZ) to the light zone (LZ), where cells are selected and differentiate either into MBCs or PCs (De Silva and Klein, 2015). At low resolution, we identified two predominant clusters consisting of NBCs/MBCs and GCBCs (Figure 1B). Although NBCs and MBCs reflect different maturation stages, they converge into similar transcriptomes (Figure 1B) (Kulis et al., 2015). However, a detailed evaluation of the NBC/MBC cluster allowed us to annotate the two main states according to the expression of *FCER2* (CD23) and *TCL1A* in NBCs and *CD27*/*CD267* in MBCs (Figure 4A and 4B). We identified six MBC-related subclusters which were mostly classified according to their Ig isotype (class switch IgA/G vs. non-class-switch IgM/D), and the expression of *FCRL4/5* (Figure 4A and 4B, Figure S6A) (Ehrhardt et al., 2008; Li et al., 2020). NBCs were divided into eight clusters depending on their activation level. To map the NBC-to-GCBC transition, we included some non-proliferative DZ-GCBCs in subsequent analysis.

**Figure 4.**
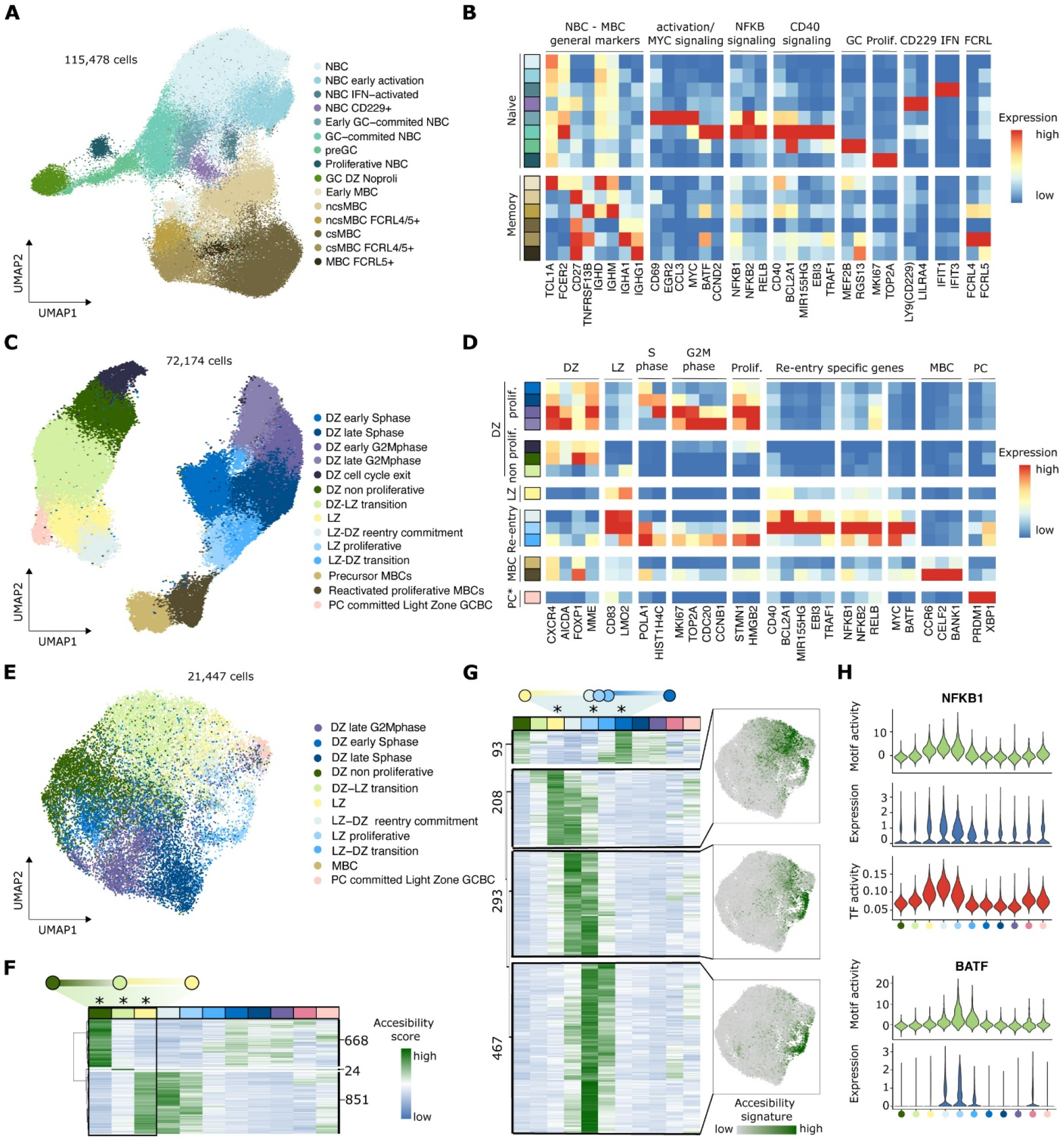
B cell activation and GC dynamics. (**A**) UMAP projection of tonsillar Naive (NBC) and Memory (MBC) B cells colored by scRNA-seq clusters. (**B**) Heatmap showing scaled mean marker expression per NBC and MBC subpopulations. GC DZ Noproli cells are included from the GCBC cluster, and will be described in panel 4D. (**C**) UMAP projection of tonsillar GCBCs colored by scRNA-seq clusters. (**D**) Heatmap showing scaled mean marker expression per GCBC subpopulations. Note that the PC* corresponds to the PC-committed Light Zone GCBC from the PC section in Figure 5. (**E**) UMAP projection of GCBC colored by scATAC-seq clusters. (**F**) Heatmap showing normalized accessibility scores of the differential accessible regions (DARs) in the DZ-to-LZ transition (DZ no proliferative → DZ-LZ transition → LZ). Numbers indicate the number of DARs included in each cluster. Asterisks highlight the columns for which the differential accessible analysis was performed. (**G**) Left: Heatmap showing normalized accessibility score of the DARs in the LZ-to-DZ re-entry (LZ → LZ-DZ reentry commitment → LZ-proliferative → LZ-DZ transition → DZ early S phase). Numbers indicate the number of DARs included in each cluster. Asterisks highlight the columns for which the differential accessible analysis was performed. Right: UMAP projection highlighting the accessibility signature scores for each of the main clusters. (**H**) Violin plot showing motif activity (green), gene expression (blue) and TF activity (AUCell score, red) for NFKB1 and BATF. BATF TF activity is not shown, as it was not identified as a regulon by pySCENIC.

Zooming into the naive compartment, we identified a cluster of initially activated NBCs, which, as compared to resting NBCs, showed a moderate upregulation of early activation markers (*e*.*g. CD69*) (Figure 4A and 4B, Figure S6A). We also identified two NBC subclusters respectively characterized by the expression of interferon-induced genes and *LILR4A/LY9* (CD229) (Figure 4A and 4B, Figure S6A). We then observed a subcluster with strong expression of *MYC, CD69, EGR2/3, and CCL3/4*, which seems to correspond to early GC committed cells due to the transient MYC upregulation (Figure 4A and 4B, Figure S6A) (Dominguez-Sola et al., 2012). We also found further GC-committed cells characterized by *CCND2* (a bona fide downstream target of MYC), *TRAF1/4*, and *MIR155HG* (Figure 4A and 4B, Figure S6A). In line, these two GC-committed populations showed upregulation of the NF-kB genes (Figure 4B, Figure S6A) (Dominguez-Sola et al., 2012). Next, we detected a subcluster of pre-GC cells, which downregulated most of the genes transiently expressed in GC-committed cells and showed early seeds of GC-specific genes (*MEF2B* and *RGS13*; Figure 4A and 4B, Figure S6A). Interestingly, we detected a subpopulation of proliferative cells with a NBC transcriptome lacking any GC marker (Figure 4A and 4B, Figure S6A). We hypothesize that this population may correspond to primary focal reaction upon very early antigen stimulation, which leads to the generation of MBCs in a GC-independent manner (Jacob and Kelsoe, 1992; Taylor et al., 2012).

Once having delineated the differentiation path of NBC into DZ B cells, we next focused on the GCBC compartment. Both gene expression and cell cycle differences between DZ and LZ drove transcriptomic variability in GCBCs (Figure 4C and 4D; Figure S6B and S6C). The latter represents the major histological feature of tonsillar follicles, where DZ-GCBCs proliferate and undergo somatic hypermutation (SHM) in their Ig genes, before rapidly differentiating into LZ-GCBCs. These are then stringently selected for improved BCR affinity and then either return to the DZ or give rise to MBCs and PCs (De Silva and Klein, 2015). In line, we observed subclusters of early MBCs (*CCR6*) (Suan et al., 2017) and early PCs (*XBP1, PRDM1* and *IRF4*) derived from LZ-GCBCs, and identified a cluster of proliferative MBC that may correspond to reactivated MBCs that rapidly proliferate and differentiate into PCs (Figure 4C and 4D; Figure S6B) (Moran et al., 2018). LZ-GCBCs experience a re-entry into the DZ for additional cycles of SHM and clonal expansion (De Silva and Klein, 2015). Therefore, DZ- and LZ-GCBCs cyclically transit into each other. Here, we identified DZ cells that gradually decrease the expression of cell cycle genes and transit through a subcluster with an intermediate DZ-LZ phenotype, before giving rise to LZ-GCBCs (Figure 4C and 4D: Figure S6B). DZ-LZ transition cells have low expression of DZ markers *(e*.*g. CXCR4*) and start expressing a fraction of LZ markers *(e*.*g. LMO2*). However, other classical LZ markers such as *BCL2A1* are still not expressed (Figure 4C and 4D; Figure S6B). For LZ-GCBCs, we observed several subclusters related to the re-entry to the DZ (Dominguez-Sola et al., 2012), from DZ-committed LZ cells, with a transitory expression of *MYC, BATF, MIR155HG*, and *TRAF1/4* to proliferative cells maintaining the LZ phenotype but upregulating S phase genes, and finally to proliferative cells with an intermediate LZ-DZ phenotype, with loss of CD83 and upregulation of cell cycle progression genes (Figure 4C and 4D; Figure S6B).

Next, we studied how the chromatin accessibility profile is modulated during the DZ-to-LZ and LZ-to-DZ transitions. A label transfer allowed the identification of transcriptional subclusters related to scATAC-seq profiles (Figure 4E). We noticed that cell cycle activity, which generated two major transcriptional clusters, does not profoundly impact chromatin accessibility, suggesting the cyclic dynamics between the DZ and LZ to homogenize the chromatin accessibility landscape in spite of extensive transcriptional changes. However, DZ-to-LZ and LZ-to-DZ transition chromatin profiles were remarkably different. The DZ-to-LZ transition was seamless, with most of the DZ and LZ-specific differentially accessible regions showing an intermediate level and only 24 specific regions (Figure 4F). In sharp contrast, for a LZ B cell to return to the DZ, widespread epigenetic programming seems to be necessary. We identified three main modules that transiently increased chromatin accessibility as LZ-GCBCs dedifferentiate through different sub-clusters to return to the DZ-GCBC state (Figure 4G). To investigate the mechanisms leading to such chromatin remodeling, we performed TF motif enrichment analysis. Remarkably, the initial commitment to the DZ was strongly enriched in binding sites of NF-kB family TFs, while NF-kB binding sites were gradually replaced by AP-1 family footprints during cell state shifts towards DZ-specific profiles (Figure 4H). In particular, BATF, which is known for controlling AID (AICDA) expression in DZ-GCBCs and for its involvement in the CSR process (Ise et al., 2011), was highly enriched in more advanced DZ committed states (Table S12). Accordingly, we observed an increased gene expression of NF-kB family members and *BATF* at the beginning and end of the LZ-DZ transition, respectively (Figure 4H, Table S12). Altogether, these results suggest that transient epigenetic programming, initially through NF-kB followed by BATF, play a key role in determining the fate of LZ-GCBCs for returning to the DZ for subsequent rounds of affinity maturation. Most remarkably, NF-kB activation followed by BATF upregulation was also observed in activated NBCs differentiating into DZ-GCBCs (Figure S6D and S6E). These results confirm and extend previous observations focused on *MYC* (Dominguez-Sola et al., 2012), indicating that similar molecular mechanisms are necessary for a B cell, either NBC or LZ-GCBC, to become a DZ-GCBC. In contrast, this phenomenon was not observed in LZ-GCBCs that further differentiate into PCs, which is the focus of the next section.

### Plasma cell differentiation and cell identity regulation in human tonsils

Antibody-secreting PCs play a key role in humoral immunity and represent the final maturation stage of the B cell lineage (Nutt et al., 2015). To properly study transitions from B cells to PCs, we added selected DZ, LZ and MBC cells to the PC compartment as well as LZ GCBCs with early PC features (Figure 5A). This last subpopulation borrowed from the GCBC cluster presented the first seeds of PC commitment by expressing the three key PC transcription factors (*IRF4, PRDM1, XBP1*) while still maintaining GCBC and LZ markers (*e*.*g. BCL6* and *CD83*; Figure 5A and 5B). Further, early PC precursors showed a remarkable drop of B cell program genes, maintained PC transcription factors and increased the expression of PC-specific genes (*SLAMF7* and *MZB1*; Figure 5A and 5B). These early PC precursors could be further divided into IgG+ or IgM+. We also identified a small PC precursor subpopulation clustering with DZ- and LZ-GCBCs. These cells showed an upregulation of DZ markers (*AICDA* and *CXCR4*), and may represent the human counterpart of a precursor PC subpopulation described in mice that migrates from the LZ to the DZ, and leaves the GC at the DZ-T interface (Kräutler et al., 2017; Zhang et al., 2018). We next identified a subpopulation with upregulated proliferation-related genes, potentially representing precursors poised towards proliferation. Gradually increasing cell cycle activity, a transitional cluster showed intermediate levels of proliferation, in contrast to proliferative plasmablasts, in which we also identified clonal expansion (Fig S7A). It is interesting to note that the increased proliferation signature is concomitant to increased expression of PC-related genes, in which G2M-phase cells show higher expression of these genes than S-phase cells (Figure S7B). These observations confirm previous findings using experimental models in which cell division is coupled to PC differentiation (Barwick et al., 2016; Caron et al., 2015).

**Figure 5.**
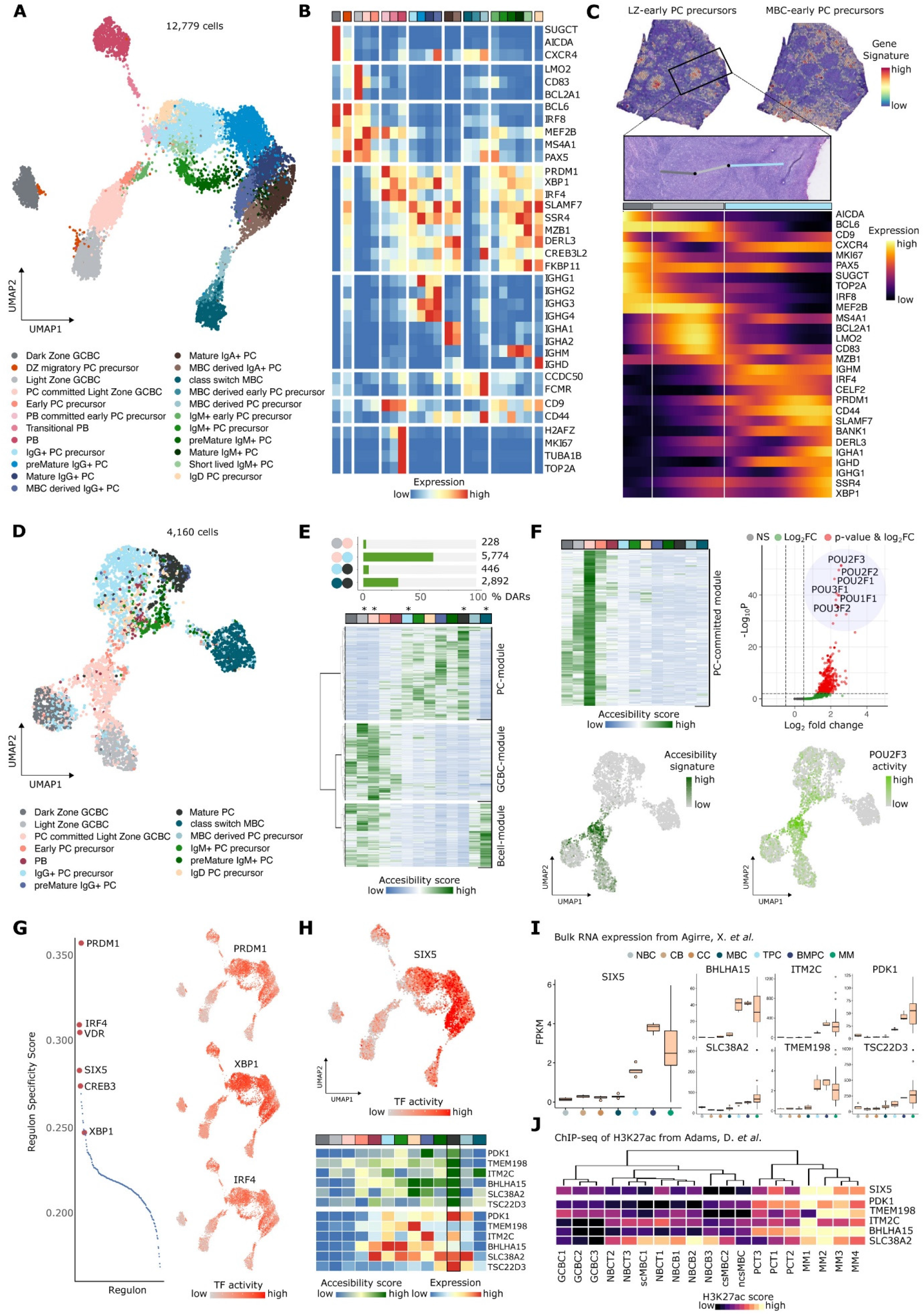
Plasma cell differentiation and cell identity regulation in human tonsils. (**A**) UMAP projection of tonsillar plasma cells (PC) colored by scRNA-seq clusters. (**B**) Heatmap showing scaled mean marker expression per PC subpopulation. (**C**) Spatial transcriptomics section from a representative slide. *Top:* Transcriptomics-based tissue localisation of LZ-derived early PC precursor (left) and MBC-derived early PC precursor (right) using the top 25 marker genes for each population (STAR Methods). *Middle:* DZ (dark grey) to LZ (light grey) to subepithelial-PC-rich zone (light blue) trajectory on an H&E image from the highlighted area. *Bottom:* Heatmap showing smoothed expression changes for genes of interest through the pre-defined trajectory. (**D**) UMAP projection of PC colored by scATAC-seq clusters. (**E**) *Top:* Proportion of differentially accessible regions (DARs) identified by a pairwise differential analysis between LZ, PC-committed, IgG PC precursor, Mature PC and csMBC. *Bottom:* Clustered heatmap representation of the normalized accessibility score from the 9,340 DARs of the three main modules. (**F**) PC-committed module analysis. *Left:* Heatmap showing normalized accessibility score of the 654 DARs and UMAP projection of their combined accessibility signature. *Right:* Motif enrichment analysis of the 654 DARs (p cutoff: 10e-03, FC cutoff: 0.5) and UMAP projection of top motif (POU2F3) activity (Fold enrichment: 2.57, Pvalue: 7.54e-52). (**G**) *Left:* Regulon specificity score for the PC subpopulation. The top 5 regulons and XBP1 are highlighted in red and labeled in the plot. *Right:* UMAP projection highlighting the activity (AUCell score) of PRDM1, XBP1 and IRF4 TFs. (**H**) *Top:* UMAP projection highlighting the activity (AUCell score) of SIX5. *Bottom:* Heatmap showing scaled mean accessibility and gene expression for each of the predicted targets of SIX5. (**I**) Boxplot of FPKM (fragments per kilobase per million fragments mapped) values for SIX5 and its target genes from Aguirre *et al*. NBC, Naïve B Cell; CB, Centroblast; CC, Centrocyte; MBC, Memory B Cell; TPC, Tonsillar Plasma Cell; BMPC, Bone Marrow Plasma Cell; MM, Multiple Myeloma. (**J**) Heatmap showing normalized mean H3K27ac signal for SIX5 targets in NBCT, tonsillar NBC (n=3); NBCB, NBC from peripheral blood (n=3); GCBC (n=3), csMBC, class-switch MBC (n=2); ncsMBC, non-class switch MBC (n=1); PC (n=3); MM (n=4).

Following the proliferative stage, two major paths emerged depending on IgG or IgM expression. This gradual maturation process was characterized by further decreased expression of B cell related genes as well as increased expression of PC markers and CD44, Ig genes, and an endoplasmic reticulum (ER) signature (Figure 5A and 5B; Figure S7C and S7D). As for IgD and IgA isotypes, we did not observe a maturation gradient. Instead, we detected a specific precursor IgD+ PC cluster and scattered IgA+ precursors within the IgG+ precursor cluster. Mature IgA+ cells showed a distinct subpopulation, while individual mature IgD+ cells dispersed within mature IgG/A+ clusters (Figure S7C). Interestingly, we also identified a small IgM+ cluster in the precursor area, showing a global gene expression downregulation with the exception of an upregulated ER signature and IgM expression, which may represent short-lived IgM PCs responsible for the early humoral response (Figure 5A and 5B; Figure S7D) (Seifert and Küppers, 2016).

Additionally, PCs can also be generated from MBCs upon antigen re-exposure (Nutt et al., 2015). We identified a first stage of early MBC-derived PC precursor cells characterized by low expression of MBC genes (*BANK1, TXNIP* and *CELF2*) and B cell markers as well as the presence of key PC genes (Figure 5A and 5B; Figure S7C). A second cluster, annotated as MBC-derived PC precursors, displayed an increase of PC markers together with a further downregulation of the MBC program (Figure 5A and 5B; Figure S7C). Finally, we identified potential IgG+ and IgA+ mature PCs derived from MBC that maintained traces of MBC-related genes (Figure S7C). Whether these cells further differentiate to reach the same phenotype of mature PCs derived from LZ-GCBCs remains an open question. To investigate spatially mapped PC transcriptional dynamics in the tonsillar tissue, we defined signatures and inferred cell proportions for PC precursors derived from LZ-GCBCs or MBCs and visualized these in the spatial context (Figure 5C; Figure S7E). In line with their single-cell derived annotation, precursor PCs derived from LZ-GCBCs mapped within the follicles, whereas those derived from MBCs were located extrafollicularly (Figure 5C; Figure S7E).

We then analyzed gene expression changes throughout a spatially defined trajectory from an intrafollicular zone to a subepithelial zone in different follicles from different tissue sections (Figure 5C; Figure S7F). Using subpopulation-specific markers, we observed the transition from the DZ to the LZ within the follicle, including initial expression of PC genes in the LZ and a strong PC signature increasing towards the subepithelial zone, where mature PCs locate (Figure 5C; Figure S7F) (Steiniger et al., 2020). Interestingly, a spatial expression correlation analysis revealed that the PC region contained distinct IgM/D and IgG/A areas (Figure S7G). Taken together, single-cell and ST results suggest that early PC differentiation in the tonsil consists of more subpopulations and states than previously appreciated (King et al., 2021b, 2021a), being the primary reaction from follicular LZ-GCBCs associated with a stepwise progression towards mature PCs, and the extrafollicular secondary reaction from MBCs being a far more direct and rapid differentiation path into mature PCs.

We next studied chromatin accessibility and transcriptional regulation during PC maturation and grouped the scRNA-seq subpopulations into 13 scATAC-derived clusters (Figure 5D). Again, the accessibility profiles were not affected by proliferation, and we confirmed that proliferation is an early event in PC differentiation, with PBs clustering together with precursor PCs. A pairwise differential accessibility analysis revealed highest differences between committed PCs and PC precursors, and from MBC to mature PCs (Figure 5E), highlighting the need of extensive chromatin programming for the cell fate transitions. Clustering all DARs, identified three main modules of chromatin dynamics related to PC, GCBC and B-cell subpopulations (Figure 5E). Individually analyzing TF binding motifs in these modules revealed overrepresentation of TF motifs related to (1) the IRF family (*IRF8, IRF4*) and *MESP1* in the PC module, (2) to *EBF1, POU2F2/F3, PAX5, NFKB1, RELA/RELB* and *MEF2C* in the GCBC module, and (3) to *EBF1, PAX5, SPIB, SPI1 and ETV3/ETV6* in the B cell module (Table S13). Remarkably, we identified 654 DARs within the GCBC module showing increased chromatin accessibility in PC-committed LZ-GCBC. These DARs were strongly enriched in POU TF binding sites (i.e. *POU2F1, POU3F1, POU2F2*), a TF family described to be crucial for PC differentiation towards an antibody secreting phenotype, particularly *POU2F1* (OCT1) and *POU2F2* (OCT2; Figure 5F) (Corcoran et al., 2014; Emslie et al., 2008; Shah et al., 1997).

To obtain a deeper insight into the regulatory mechanisms underlying PC differentiation, we complemented the epigenomic with GRN analysis (Tables S14 and S15). Compared to all other B cell populations, the key PC regulators IRF4, PRDM1 and XBP1, together with previously reported VDR (Lin et al., 2017) and CREB3 (Al-Maskari et al., 2018), defined PC-specific regulons (Figure 5G; Figure S7H; Table S16). Intriguingly, the homeobox protein SIX5 (Sarkar, 2004; Sarkar et al., 2000), a TF not yet described in PCs, showed lineage-specific regulatory activity. SIX5 regulon activity suggested the TF to be involved in later stages of PC maturation (Figure 5H). These findings were further supported using bulk RNA-seq (Agirre et al., 2019) and H3K27ac ChIP-seq data (Beekman et al., 2018a) as well with scRNA-seq from peripheral blood (Hao et al., 2021) and bone marrow (Hay et al., 2018), confirming SIX5 to be highly specific for mature PCs (Figure 5I and 5J; Figure S7I and S7J). In line, the predicted target genes of SIX5 (*PDK1, TMEM198, ITM2C, BHLHA15/*MIST1, *SLC38A2* and *TSC22D3/*GILZ) showed increased accessibility and selectively expression in the mature PC (Figure 5H and 5I). In Multiple Myeloma (MM), a PC-derived neoplasm, we identified increased expression and H3K27ac signal in SIX5 and its target genes (Figure 5I and 5J) (Ordoñez et al., 2020). While PDK1, BHLH15 and ITM2C have been described in MM, the regulatory network of SIX5 could provide further insights into disease-driving mechanisms. Together, these results indicate SIX5 to be a novel marker for mature PCs with a potential role in the regulation of MM tumorigenesis.

### Non-lymphoid cells and novel roles of slancyte subsets in human tonsils

We then charted the cellular heterogeneity within the epithelial compartment, the first tonsillar barrier against pathogens. We identified six clusters, three of which overlapped with the keratinocyte populations reported in an oral mucosa cell atlas (Figure S8A and S8B) (Williams et al., 2021). Basal and surface keratinocytes presented mutually exclusive keratins, with the former expressing *KRT5* and *KRT14* and the latter *KRT13* and *KRT80* (Figure S8A and S8B) (Alam et al., 2011). Cell type deconvolution of ST data validated the spatial location of both cell types (Figure S8C). Noteworthy, surface epithelium expressed the protease *TMPRSS2* (Figure S8B), which activates coronaviruses in other tissues (Strope et al., 2020), although we did not detect the expression of the receptor *ACE2* (Figure S8B). We annotated the remaining clusters as (1) “Outer surface” with a cornification (*CNFN, LCE3* family) and keratinization (*SPRR2D*/*E*) gene signature, (2) “Crypt” with a gene signature of inflammatory responses (*IL1B* and *S100A6*) and M cells (*SPIB* and *MARCKSL1*), specialized in antigen uptake (Kobayashi et al., 2019), and (3) “FDCSP epithelium”, expressing *FDCSP* and *KRTDAP* (Figure S8A-S8C). *FDCSP* was first described in FDC and in “leukocyte-infiltrated tonsillar crypts” (Marshall et al., 2002), although the specific population within the crypts remained unknown. Here, we provide evidence that FDSCP expressing cells represent a specific subpopulation of the tonsillar epithelium.

After M cells transport antigens to the stroma of the tonsil, they are sampled by DCs. The transcriptional heterogeneity within the tonsillar DC compartment was remarkably consistent with the one observed in blood (Villani et al., 2017). We identified all previously described DC subsets: (1) DC1, conventional DC1 (cDC1, *e*.*g. CLEC9A*), divided into precursor and mature states on the basis of *XCR1* expression (Balan et al., 2018); (2) DC2 and DC3, corresponding to cDC2 (*CD1C, FCER1A, CLEC10A*) and differing in their antigen-presenting capacity (*e*.*g. HLA-DQB1*) and inflammatory signatures (*S100A8, S100A9*), respectively; (3) DC4 (*FCGR3A, SERPINA1* and *FTL*) putatively derive from non-classical monocytes; and (4) DC5 expressing *AXL* and *SIGLEC6* (AS), the hallmark markers of AS DCs (Figure 6A and 6B; Figure S9A). AS DCs represent a continuum of cell states between antigen-presenting cDC and interferon-producing pDC (Villani et al., 2017). Accordingly, DC5 expressed *LILRA4*, which was highly expressed in pDC.

**Figure 6.**
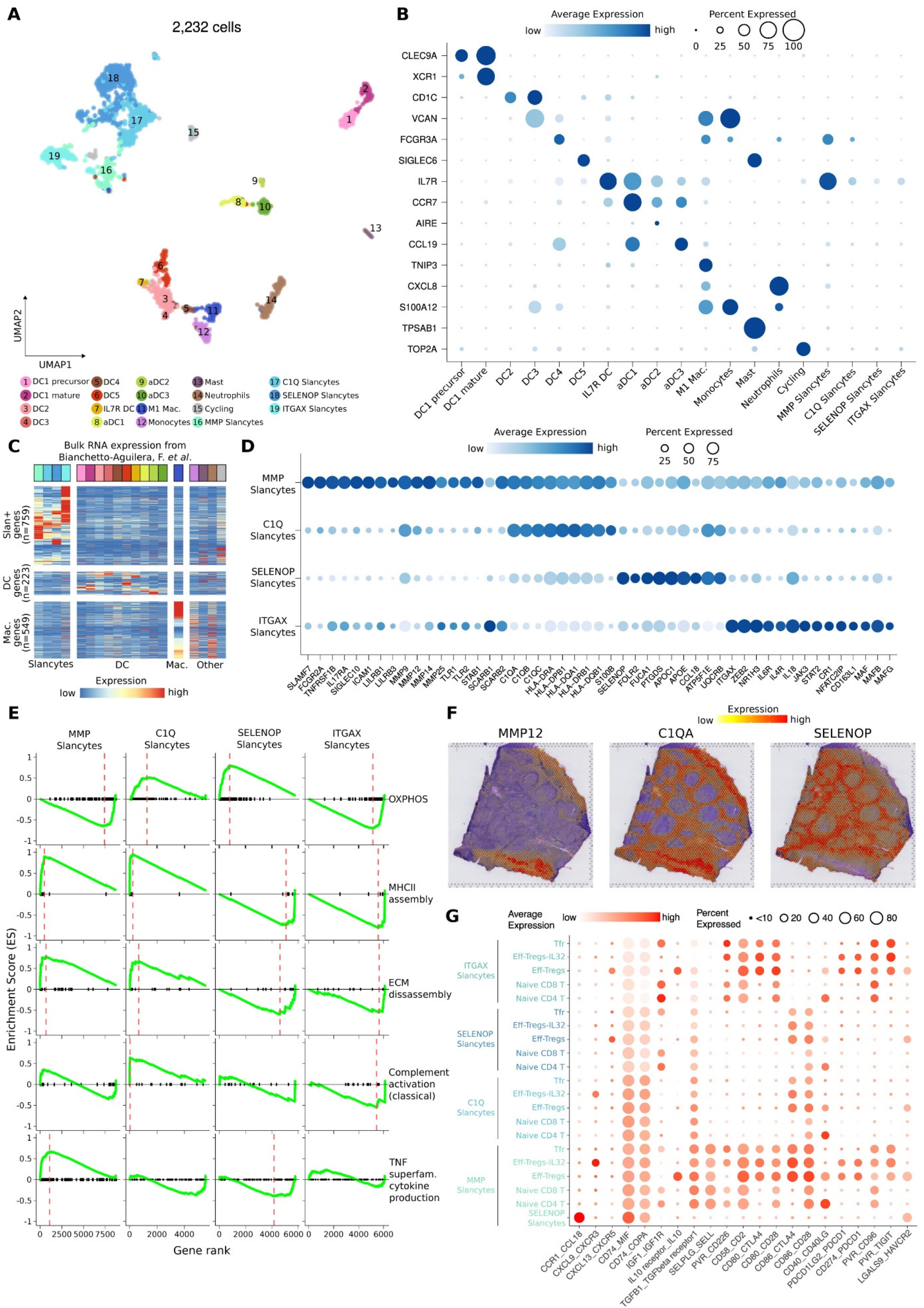
Myeloid cells and novel roles of slancytes in human tonsils. (**A**) UMAP projection of tonsillar myeloid cells colored by scRNA-seq clusters. (**B**) Dotplot showing the average gene expression for the top 15 markers *(y-axis)* for myeloid subpopulations *(x-axis*). Dot size reflects the percentage of expressing cells and the color shows the average expression level. (**C**) Heatmap showing scaled expression of slan+ vs DC vs macrophages differentially expressed genes from Bianchetto-Aguilera *et al*. in our myeloid subpopulations. (**D**) Dotplot showing expression of the top marker genes *(x-axis*) per slancyte subpopulations *(y-axis)*. (**E**) Gene Set Enrichment Analysis (GSEA) plots showing the running enrichment score (ES) for selected enrichmed GO terms across slancyte subpopulations. Vertical red dashed lines mark significant maximum ES (FDR-adjusted p-value < 0.05). Black vertical lines in the x-axis show genes in ranked gene lists found in each GO. (**F**) MAGIC-denoised (STAR Methods) expression of genes of interest identifying slancyte populations on an ST slide. (**G**) Dotplot showing potential cell-to-cell interactions between slancytes and subpopulations of naive T cells and T regulatory cells. Interacting ligand/receptor pairs (*x-axis*) show the gene expressed by slancytes followed by the gene expressed by the T cell subpopulation. Color shows the average of the expression of the ligand and receptor (each in its population) and dot size the average percentage of expressing cells.

Intriguingly, we identified a previously uncharacterized cluster that expressed *AXL* and *IL7R* (but not *SIGLEC;* Figure 6B; Figure S9A). In addition, IL7R DCs expressed *IL1RN, IL1B, CD83, AREG*, and *IFI30* (Figure S9A). We also found three populations of migratory CCR7+ DC (activated DC, aDC) that were characterized in the thymus(Park et al., 2020) and have been observed in tonsils (Figure 6A and 6B; Figure S9A) (Villar and Segura, 2020). However, in contrast to the thymus, we could not assign a specific DC origin to each aDC subset (Figure S9B). Noteworthy, aDC2 expressed shallow levels of Autoimmune Regulator (*AIRE*; Figure 6B), which has a role in upregulating tissue-specific antigens to generate peripheral tolerance(Gardner et al., 2013; Poliani et al., 2010; Wang et al., 2021). Finally, we annotated four clusters as monocytes (*VCAN, S100A12*), M1 macrophages (*TNIP3*), mast cells (*TPSAB1, TPSB2*) and neutrophils (*CXCL8, PI3*).

Tonsil 6-sulfo LacNAc+ (slan+) cells (hereafter named slancytes) have been described as a population derived from non-classical monocytes that is transcriptomically distinct from tonsillar cDC2 and macrophages (Bianchetto-Aguilera et al., 2020). In this setting, we investigated whether slancytes represent a single homogeneous population or, conversely, consist of several heterogeneous subsets. To that end, we quantified the slan+ cells, macrophage and cDC2 gene expression signatures (Bianchetto-Aguilera et al., 2020) for the 19 clusters of the myeloid compartment of our tonsil atlas. Strikingly, we identified four clusters that displayed a high expression of genes specifically upregulated in slan+ cells in comparison to tonsillar DCs and macrophages (Figure 6B and 6C). These four clusters were the most prevalent cell types of myeloid cells in tonsils (Figure S9C), and we annotated them on the basis of their specific slancyte marker genes. (1) MMP slancytes expressed metalloproteinases (*MMP* family), and toll-like receptors (*TLR1*/2); (2) C1Q slancytes expressed members of the classical complement pathway (*C1Q* family), and class II MHC genes; (3) SELENOP slancytes expressed apolipoproteins (*APOC1, APOE*) and fucosidases (*FUCA1*); and (4) ITGAX slancytes expressed scavenger receptors (*SCARB1/2*), among other specific marker genes (*e*.*g. ITGAX, CR1*) (Figure 6D). Gene set enrichment analysis (GSEA) revealed that C1Q and SELENOP slancytes have increased oxidative phosphorylation and metabolic activity, while C1Q and MMP slancytes show increased antigen presenting and extracellular matrix disassembly capacity (Figure 6E). MMP slancytes also produce more inflammatory cytokines, such as a tumor necrosis factor (TNF; Figure 6E). Because *SELENOP* was vastly specific to SELENOP slancytes across the 121 cell types and states of the tonsil atlas (Figure S9D), we used it as a proxy of the spatial location of these slancytes. Noteworthy, *SELENOP* was mostly expressed at the interfollicular/T cell zone; while it was absent in the epithelium and follicles (Figure 6F). *MMP12* was expressed subepithelial, while *C1QA* was expressed both subepithelial and at the interfollicular zones. IL7R DCs also expressed *MMP12* and *C1QA* (Figure S9D), however the low prevalence of this cell type (Figure 6A) suggests the main source of *MMP12* and *C1QA* to be the slancyte populations. Finally, the slancyte subtypes markedly differed in their cell-cell interactome (Figure 6G). Cell-cell interaction analyses confirmed the different roles in immune activation, antigen presentation, cytokine production and migration patterns of each slancyte subtype. Taken together, we have discovered four previously uncharacterized types of slancytes that might control different scenarios of immune responses and interact with distinct subtypes of immune cells, depending on their location.

We next classified cells of mesenchymal origin. Recently, single-cell transcriptomic profiling of tonsillar mesenchymal cells uncovered three major subsets: marginal reticular cells (MRC), fibroblastic reticular cells (FRC), and FDC (Heesters et al., 2021). In line, we identified MRC (*DCN, DIO2*) and a cluster which shared markers with the FRC (*PERP, CCL20*; Figure S8D and S8E). Notably, we found three subsets of FDCs: COL27A1+ FDC and a cluster of FDCs that expressed *CD55* and *CD14* (Figure S8D and S8E). Of note, CD14+ FDC are associated with poor prognosis in follicular lymphoma (Smeltzer et al., 2014). All FDCs expressed *FDCSP, CLU, CXCL13* and *VIM*, although COL27A1+ FDC at a lower level (Figure S8D and S8E). COL27A1+ FDC and MRC differed in their expression of collagens. MRC expressed higher levels of for example *COL1A1, COL1A2* and *COL3A1*, but lower levels of *COL27A1* (Figure S8D and S8E). Collagens expressed by MRC localized mostly at the interfollicular zone (Figure S8F). Intriguingly, MRC expressed *PDGFRB*, which has been shown to be specific to perivascular precursor FDC in mice (Krautler et al., 2012).

### Disseminating and leveraging the translational potential of the tonsil data atlas

A core value of the HCA is transparency and open data sharing (Regev et al., 2017). To make our data FAIR (Findable, Accessible, Interoperable and Reusable) (Wilkinson et al., 2016), we developed HCATonsilData: an R/BioConductor data package that provides modular and programmatic access to the tonsil atlas dataset. With minor efforts, users can access SingleCellExperiment (Cph, 2017) objects for different cell types and data modalities, easily convertable to AnnData (Wolf et al., 2018) objects via zellkonverter (Luke Zappia, Aaron Lun), ensuring interoperability across programming ecosystems. In addition, the dataset is extensively documented, so users can easily trace back the reasoning behind any cell type annotation. We also would like to highlight that we provide a detailed Glossary listing supportive evidence for the annotation of all the 121 cell types and states of this tonsil atlas (Glossary). For the relevant subsets presented in the manuscript, we prepared iSEE instances (Rue-Albrecht et al., 2018), enabling interactive exploration of marker genes and metadata of the tonsillar cell types (Figure 7A).

**Figure 7.**
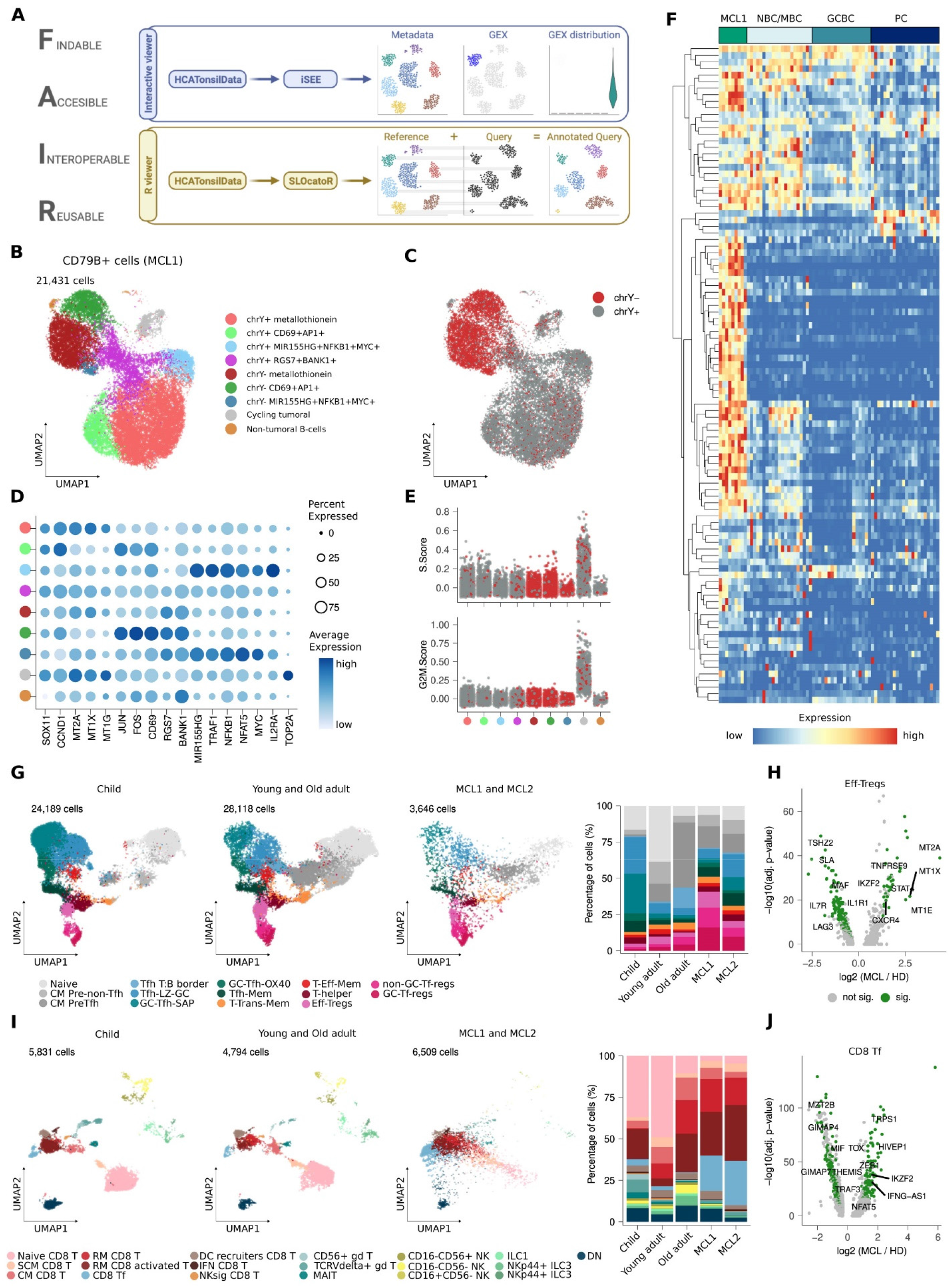
Disseminating and leveraging the translational potential of the tonsil atlas. (**A**) Schematic representation showing the computational framework to access and reuse the tonsil atlas dataset in a FAIR manner. (**B**) UMAP projection of CD79+ cells from an MCL case (MCL1) colored by scRNA-seq clusters. (**C**) UMAP projection of CD79+ cells from MCL1 colored by chrY status (+ or -). (**D**) Dotplot showing the average expression of the top 17 markers *(x-axis*) across scRNA-seq clusters from CD79+ cells from MCL1 *(y-axis)*. Dot size reflects the percentage of expressing cells and the color shows the average expression level. (**E**) Jitter plot showing the cell cycle signature score; S.Score (*top*) and G2M.Score (*bottom*) across clusters. Color represents the chrY status: + (gray) or - (red). (**F**) Heatmap showing scaled mean gene expression for 100 top genes upregulated in MCL for MCL1 and healthy B cell clusters (NBC-MBC, naive and memory B cells; GCBC, Germinal Center B cells; PC, Plasma Cells). Dendrogram represents the hierarchical cluster of the genes (rows). (**G, I**) *Left:* UMAP projection of reference [tonsillar CD4 T cells (G), Cytotoxic cells (I)] and query (MCL cells) annotated with SLOcatoR. *Right:* Stacked barplot showing the cell type proportion across age groups and disease status. (**H, J**) Volcano plot showing the log_2_ fold-change (*x-axis*) versus the significance (*y-axis*) of the differential expressed genes between MCL and healthy donors (HD) for Eff-Tregs (H) and CD8 Tf cells (J) colored by significance (adjusted p-value < 0.01, log_2_ FC > 0.3, difference in % of expressing cells > 0.25).

We then developed SLOcatoR: an R package that annotates unseen transcriptomes and open chromatin profiles from SLO using the tonsil atlas as reference. SLOcatoR generalizes the methods we developed to bridge different data modalities. Both the reference and the query can be profiled with single-cell or single-nuclei assays, even though transcriptomes largely differ between these cellular compartments (Mereu et al., 2020). To mitigate this, SLOcatoR first finds technology-independent highly variable features. Following principal component analysis (PCA), SLOcatoR integrates the query and the reference with Harmony (Korsunsky et al., 2019). Harmony scales well to atlas-level dataset and ranks among the best-performing tools for both scRNA-seq and scATAC-seq (Chazarra-Gil et al., 2021; Luecken et al., 2022; Tran et al., 2020). Finally, SLOcatoR transfers the labels and the UMAP coordinates with KNN classification and regression, respectively. Plotting the annotation probability in the learnt UMAP coordinates informs about potential misclassifications (Figure 7A).

To demonstrate SLOcatoR feasibility, we performed Multiome profiling of two tonsils from two different patients affected by conventional mantle cell lymphoma (MCL), an aggressive B cell neoplasm arising in lymphoid tissues, including the tonsil (Tashakori et al., 2021). Tumor B cells were identified by the B cell marker *CD79A* and the MCL-specific markers *CCND1* and *SOX11* (Beekman et al., 2018b), whereas the remaining cells were classified as tumor microenvironment (TME). We initially explored the neoplastic cells, which were characterized by transcriptional heterogeneity, with two major clusters containing several subclusters (Figure 7B). For each patient, we noticed that one of the two major clusters showed reduced expression of genes located on chromosome Y (*UTY, KDM5D, DDX3Y;* Figure S10A) and collapsed these into a “chrY expression score” per cell (Figure S10B). This score followed a bimodal distribution, which allowed us to classify neoplastic cells into chrY+ and chrY- (Figure 7B and 7C; Figure S10B-S10D). Of note, loss of chromosome Y is a common feature of MCL (Nieländer et al., 2008). In patient 1, we observed four subclusters, which were similar in both chromosome Y+ and Y-clusters, whose characteristic genes were (1) metallothionein genes, (2) AP1-related genes and *CD69*, (3) *MIR155HG, NFKB1* and *MYC*, and 4) *BANK1* and *RGS7* (Figure 7B-7D). Remarkably, several of these genes are also associated with normal B cell subpopulations, *i*.*e*., from activated NBCs to pre-GC cells (Figure 4A and 4B). This finding suggests that MCL cells are not frozen on a particular maturation stage, but span a particular window of normal B cell differentiation. Interestingly, we also detected a relatively small cluster containing proliferative MCL cells (Figure 7E). Overall, similar findings were observed in the second MCL patient, including the subclonal loss of chromosome Y (Figure S10C-S10F).

Our tonsil atlas can also be exploited to identify MCL-specific genes whose expression is not modulated during physiological B cell maturation. We exemplified this with 100 top genes differentially expressed in MCL as compared to B cells using bulk transcriptomics (Nadeu et al., 2020). We then created pseudo-bulk transcriptomic profiles for these 100 genes across all the clusters defined for MCL, NBC, MBC, GCBC and PC. Hierarchical clustering of these genes revealed a module of MCL-specific genes (Figure 7F; Figure S10G). We observed genes inside this module only expressed in one or a few clusters of neoplastic cells, again highlighting the discriminative power of single-cell analysis.

Another strength of the tonsil atlas is to identify potential compositional changes in the TME. Projecting the CD4 T cells onto the tonsil atlas reference revealed an enrichment of the three types of Treg cells reported in the atlas. In line, FOXP3+ Treg cells were recently shown to be enriched in the TME of aggressive MCL (Balsas et al., 2021). Here, we confirm and further expand this finding by the identification of distinct sets of Treg subpopulations (Figure 7G). In addition, we found a strong shift towards terminally differentiated CD8 T cells, specifically CD8 Tf (Figure 7I). However, because CD8 Tf express *PDCD1, TIGIT* and *ICOS*, we reannotated those cells as exhausted CD8 T cells. Thus, disease-specific cell states ought to be considered when interpreting the projected annotations. Finally, we used our tonsil atlas as a healthy reference to find expression changes associated with all annotated cell types (Figure 7H and 7J). Taken together, we show that our atlas is a FAIR resource that can be exploited to deepen our understanding of hematological malignancies.

## Discussion

We provide a detailed taxonomy of cells in the human tonsil. In addition to the annotation of cell types using single-cell transcriptomics, the multimodal nature of our atlas allowed the fine-grained interrogation of subtle cell states and their driving mechanisms through gene regulatory or spatial determinants. Supportive information was retrieved across profiling modalities, highlighting the need to generate multimodal cell atlases for a holistic understanding of organ composition and function.

The high number of profiled cells allowed the detection of extremely rare cell subtypes, such as precursor B and T cells, the latter representing <0.01% of cells, and supporting an ongoing maturation of lymphoid cells outside the bone marrow and thymus, respectively. Further, the scale of this tonsil atlas enabled us to zoom into cell types to identify novel subtypes and states. In the myeloid lineage, we describe an intriguing heterogeneity within slancytes, with four subtypes forming the largest myeloid fraction in the tonsils. Different slancyte types expressed specific markers and gene sets, which, together with their distinct spatial localization, pointed to vastly specialized functions. Similarly, zooming into the CD4 T cell compartment, we identified an early bifurcation following antigen recognition by naive T cells, resulting in two distinct precursor populations of Tfh and non-Tfh CM cells. On the other extreme of the CD4 T cell differentiation trajectory, we identified two clusters of Tfh cells. Specifically, cells expressing high levels of *SH2D1A* support GC responses and the generation of MBCs and long-lived PCs, whereas *OX40* expressing cells engage with activated B cells to enhance Ig secretion of PCs (Crotty, 2011b). Subtype heterogeneity was further observed in Treg cells that control immune homeostasis, but also play a role in different diseases such as cancer and autoimmune diseases (Zaiss et al., 2019). Our data revealed three transcriptional subtypes related to classical effector phenotypes or proinflammatory and follicular subtypes, with previous described functions in the suppression of antitumor responses and severe COVID-19 and the control of B cell antibody production, respectively (Agarwal et al., 2020; Galván-Peña et al., 2021). The multimodal study design further enabled the interrogation of regulatory circuits driving cell type specialization. We describe a *BCL6* superenhancer, specifically active in follicular T cells. Intriguingly, the same region has been previously linked to *BCL6* activation in GCBCs (Bunting et al., 2016), indicating its function as follicle-specific rather than cell type-specific cis-regulatory element. Combining gene expression and the chromatin landscape further allowed to disentangle the transcription factor hierarchy associated with the DZ entry with NF-kB followed by BATF activity. Remarkably, this TF hierarchy seems to be shared in LZ cells reentering the DZ as well as activated naïve B cells that enter the DZ for the first time, further expanding previous findings focused on MYC (Dominguez-Sola et al., 2012). Charting the regulatory landscape in PCs discovered SIX5 as a novel TF associated with the maturation, but not the commitment, of PCs. TF activity is essential for the understanding of causal chains of events driving cell type specialization. With SIX5, we present a novel candidate orchestrating PC maturation with a potential role in neoplastic transformation to MM.

In total, we describe 121 cell types and states comprising the human tonsil. Our cluster annotation was based on complementary information from multiple data modalities, interpreted using the latest literature and expert-based knowledge. However, eventually it will be a community effort to agree on cell type annotation, especially for newly reported subtypes, which we facilitate through the accessibility of data, analysis code and a thoroughly assembled Glossary. To broaden the utility, we designed HCATonsilData to accommodate the dynamic nature of cell type annotation. HCATonsilData provides programmatic and modular access to our Tonsil Atlas to ease data integration and community driven annotation, such as the efforts to generate an Immune Cell Atlas across organs, tissues and immune cell lineages (coordinated by the HCA Bionetwork).

Beyond providing an atlas as a resource and reference map of the healthy human tonsil, we provide a proof-of-concept for its utility in the context of disease. MCL cells have a clonal origin, derived from a single B cell (Navarro et al., 2020). However, we discovered that MCL cells are not homogeneous in the environment of the tonsil, but rather generate an intraclonal transcriptional ecosystem with different subclusters related to normal B cell maturation. Consequently, MCL cells are not frozen in a single maturation state, but rather seem to mimic a spectrum of B cell subpopulations ranging from activated naive B cells to pre-GC B cells. In addition, mapping to our reference revealed that cells in the TME showed a striking shift towards regulatory CD4 T cell subtypes (Balsas et al., 2021) and terminally differentiated and potentially exhausted CD8 subtypes.

While current atlases of healthy human organs and tissues focus on the analysis of mostly transcriptional data, the presented atlas integrated five modalities including spatially-resolved transcriptional profiling. However, currently available ST technologies for transcriptome-wide profiling do not provide *bona fide* single-cell resolution and require capture site deconvolution to predict cell type location by integrating single-cell and ST datasets. However, novel technologies (Chen et al., 2022; Cho et al., 2021) will soon overcome such limitation, once becoming broadly accessible, and one can foresee future cell atlases to perform single-cell resolved phenotyping directly from tissue sections, avoiding tissue dissociation and related technical artifacts that could bias cell composition or gene expression profiles.

## Supporting information

Glossary

## Acknowledgements

This work was supported by the project BCLLATLAS that has received funding from the European Research Council (ERC) under the European Union’s Horizon 2020 Research and Innovation Programme (grant agreement No 810287, to H.H., J.I.M.-S., E.C., and I.G.G.). This work was also supported by the Ministerio de Ciencia e Innovación (MCI), grant agreements PID2020-115439GB-I00 (to H.H.), PID2020-118167RB-I00 (to J.I. M.-S.) and RTI2018-094274-B-I00 (to E.C.), and FEDER: European Regional Development Fund “Una manera de hacer Europa”. We also acknowledge funding from the Spanish Instituto de Salud Carlos III, Fondo de Investigaciones Sanitarias and cofunded with ERDF funds (PI19/01772), the Spanish Ministry of Science and Innovation through the Instituto de Salud Carlos III and the 2014–2020 Smart Growth Operating Program, to the EMBL partnership and co-financing with the European Regional Development Fund (MINECO/FEDER, BIO2015-71792-P). We thank the support of the Centro de Excelencia Severo Ochoa, and the Generalitat de Catalunya through the Departament de Salut, Departament d’Empresa i Coneixement, the CERCA Programme, and Suport Grups de Recerca AGAUR (2017-SGR-1142 to E.C., 2017-SGR-736 to J.I.M.-S.). Additionally, we thank the support of the Accelerator award CRUK/AIRC/AECC joint funder-partnership (to J.I.lM.-S.), the CIBERONC (CB16/12/00225, and CB16/12/00334), and the DFG (KU1315/14-1) (to R.K.). M.M.B received support from the Swiss Cancer League (BIL KLS-5130-08-2020) and the Nuovo-Soldati Foundation for Cancer Research. E.C. is an Academia Researcher of the “Institucio Catalana de Recerca i Estudis Avancats” (ICREA) of the Generalitat de Catalunya. This work was partially developed at the Centre Esther Koplowitz (CEK, Barcelona, Spain).

## Author contributions

I.G.G., E.C., J.I.M.-S. and H.H. designed the study. R.M.-B., J.I.M.-S. and H.H. supervised the work. D.M., M.K., A.V.-Z., G.C., C.M., S.R. and P.L. performed the experiment. J.C.N., M.M.B., G.F., X.A., M.A.W., R.K. and E.C. annotated cells and slides. R.M.-B., P.S.-V., S.A.-F., M.E.-B., S.R., C.A., S.P., D.G.-C. and F.M. performed the computational analysis. I.G.G., G.L., W.B., M.K. and D.M. coordinated data generation. D.C., F.J.C.-P., P.M.B., I.V., F.P. and E.C. provided clinical material. R.M.-B., P.S.-V., S.A.-F. and J.C.N. prepared the figures. R.M.-B., P.S.-V., S.A.-F., J.C.N., J.I.M.-S. and H.H. wrote the manuscript. All authors edited or commented on the manuscript.

