## Supplementary material for "An Atlas of Cells in the Human Tonsil": Glossary

### GLOSSARY OF TONSILLAR CELL SUBPOPULATIONS AND STATES

The goal of this glossary is to help readers understand the rationale behind the designation of each cell subpopulation. It is not a detailed description of the multimodal features of each subpopulation but just the information we have used to name each cell type or state. It will be divided into ten sections representing the major cell populations identified at the UMAP level 1, and the references used to annotate them.


1. Plasmacytoid Dendritic Cells and Precursors T and B cells ……………………………….. page 2
2. CD4+ T cells ……………………………………………………………………………… page 3-4
3. CD8+ T cells/innate lymphoid cells ……………………………………………………… page 5-6
4. Naive/Memory B cells ………………………………………………………………….. page 7-9
5. Germinal center B cells ………………………………………………………………… page 10-12
6. Plasma cells ……………………………………………………………………………. page 13-16
7. Myeloid cells …………………………………………………………………………… page 17-18
8. Epithelial cells …………………………………………………………………………. page 19
9. Mesenchymal cells/follicular dendritic cells ……………………………………………. page 20
10. References ……………………………………………………………………………. page 21-24


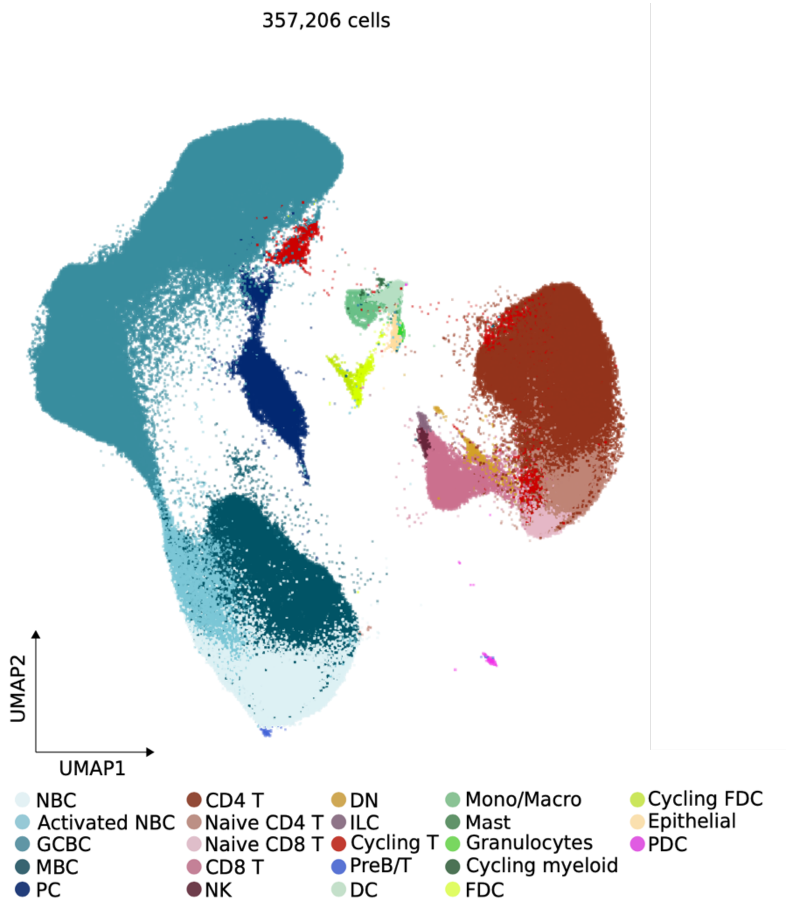

### Plasmacytoid Dendritic Cells and Precursors T and B cells

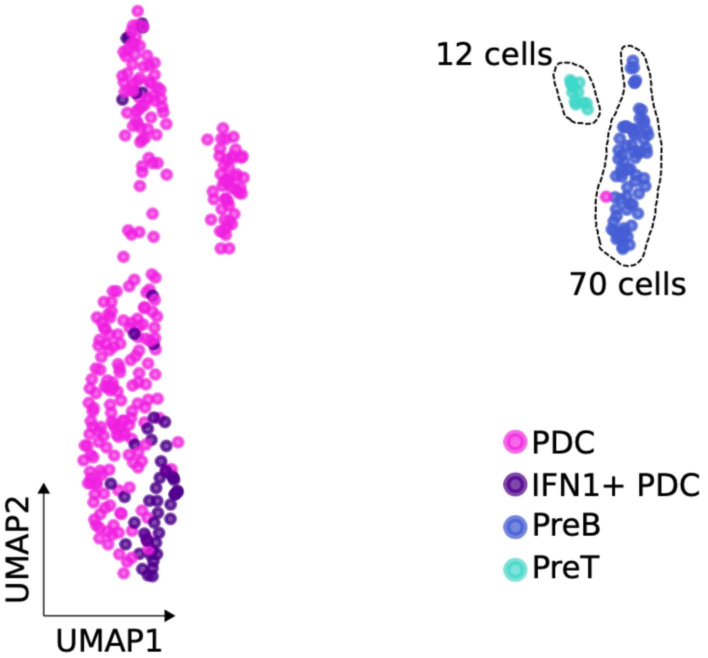


**PDC:** pDCs were previously identified by the standard protein marker CD123 (IL3RA). This cluster had higher expression of *IRF8, IL3RA* and *LILRA4* (Villani et al., 2017).

**IFN1+ PDC:** The second pDC subpopulation was identified by standard protein marker CD123 and as IFN-producing cells. This cluster had higher expression of *IRF8, IL3RA, LILRA4* and *IFI44, IFI6* (Park et al., 2020; Villani et al., 2017).

**PreB**: Precursor B cells were identified by the expression of *CD79B, CD19, PAX5, RAG1/2* (King et al., 2021).

**PreT:** Precursor T cells in tonsils expressed: *CD3G, CD8A, RAG1, DNTT, CD1A, CD1B, CD1C, CD1E* (King et al., 2021).

### CD4+ T CELLS

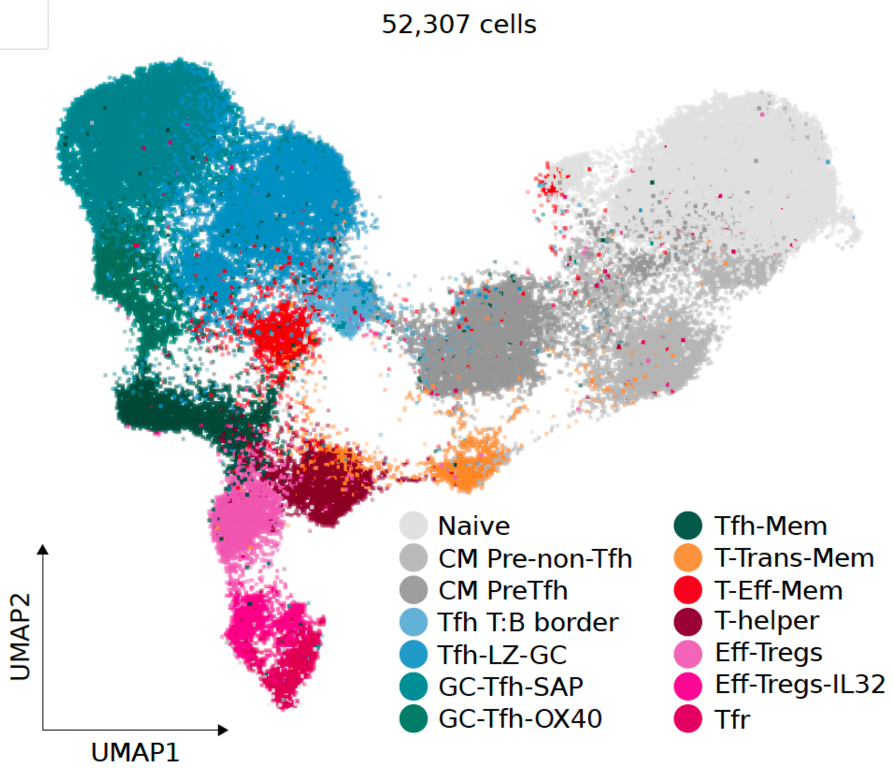


**Naive CD4 T cells**: Naïve cell is a CD4 T cell that recently migrates from the thymus towards lymph nodes. Naïve CD4 T cells expressed lymph node homing genes such as *CCR7* and naïve-associated genes: *SELL*, *LEF1, BACH2* and *NOSIP*. Also, they were negative for *PRDM1* and *BCL6* and were CD62L+ and CD45RA+ (Cano-Gamez et al., 2020).

**Central Memory Pre-non-Tfh (CM Pre-non-Tfh)**: After antigen encounter T cells acquire memory and effector functions maintaining the lymph node homing. CM showed an upregulation of *IL7R* compared to naïve T cells, and higher expression of PRDM1*, PASK, ANXA1* and *S100A4*. They had an intermediate expression of CCR7 and higher expression of CD28 confirming their migratory capacity to secondary lymph nodes and their memory profile based on their effector function. This cluster was CD45RO+, CD28+ and CD29+(Cano-Gamez et al., 2020; Nicolet et al., 2021).

**CM Pre- Tfh:** Early Tfh cell differentiation is regulated by IL-6 and ICOS. This cluster had a similar CM profile, but differential expression of follicular precursors: *TCF7, IL6ST* and *CXCR5*, and up-regulation of follicular genes such as *ICOS, PDCD1* and *TIGIT* (Qi, 2016; Sallusto et al., 2004).

**Tfh T:B border**: After antigen recognition, CD4 T cells migrate toward the border of the follicle activating B cells. This cluster showed a downregulation on the expression of *CCR7* and up-regulation of *BCL6*, *TOX*, *CXCR5 and BTLA*. The expression of *ICOS* and *CD40LG* confirmed the interaction with B cells at the border of the follicle (Crotty, 2019).

**Tfh LZ GC:** Tfh cells provide IL-21 and CD40 signals required for B cell proliferation and differentiation toward both GC and extrafollicular fates. This cluster expressed *IL21* and *CD40LG*. Also, the initial up-regulation of follicular markers as *PDCD1*. ATAC-seq analysis confirms the activity of *BCL6* in this cluster as a follicular T cell subpopulation (Crotty, 2014).

**GC Tfh SAP:** Terminally differentiated Tfh cells in GC acquired functions depending on the expression of certain markers. This cluster is expressed as a top gene *SH2D1A* (SAP) and specific genes and transcription factors associated with GC function: *CXCL13, PDCD1* and *POU2AF1*. SAP was demonstrated to be essential in CD4 T cells for GC responses and the generation of memory B cells and long-lived plasma cells (Crotty, 2011).

**GC Tfh OX40:** OX40 expression (*TNFRSF4*) on Tfh cells play a role in the maturation and Ig secretion of B cells and plasma cells during GC reaction. This cluster expressed higher levels of *TNFRSF4, CD200, IL21* and *IL4 canonical molecules of GC Tfh.* IL-21, and IL-4 are major ‘‘help’’ molecules produced by GC Tfh cells to keep GC B cells alive and induce their proliferation (Crotty, 2011). *CXCL13, IL-21* and *IL- 4* are also potent inducers of IgG1 class switch recombination for human B cells (Choi and Crotty, 2021).

**Tfh Mem:** Upon leaving a GC, the Tfh cell acquires a less activated, less polarized Tfh phenotype and can upregulate *KLRB1* and develop into resting memory Tfh cells. This cluster had the expression of follicular markers *PDCD1, MAF,* and higher expression of memory markers such as *KLRB1, ITGB1, CD28* and *IRF4* (Crotty, 2014).

**T Trans-Mem:** Transitional memory T cells appear to be more differentiated than TCM cells but not as fully differentiated as TEM cells in terms of phenotype. This cluster had higher levels of *IL7R, CD28, CCR6, RORA* and lower *KLRB1*. This cluster was CD62L-, CD45RO+ and CD28+ (Mahnke et al., 2013).

**T-Eff-Mem:** Following antigen clearance, effector cells die while a small pool of T cells ultimately develops into long-lived effector memory T cells. This cluster was positive for *KLRB1, HECW2, EGLN3* and *PRDM1.* The phenotype of this cluster included CD45R0+, CD161+ and CD62L- (Cano-Gamez et al., 2020; Mahnke et al., 2013).

**T-helper:** T helper cells play a critical role in defending infections by coordinating immune responses. CD4 T cells execute their functions mainly in peripheral tissues and various lymphoid organs. CD4 T cells receive signals from the environment and differentiate into specific subsets of T helper cells to efficiently recognize a broad range of antigens (Zhu and Zhu, 2020).

**Th1**: Th1 cells induce responses against intracellular pathogens by activating macrophages. This cluster expressed *CXCR3, TBX21, GZMK* and *CCR5,* main markers of Th1 cells (Krueger et al., 2021).

**Th2**: Th2 cells are involved in type 2 responses to fight infections by parasites such as helminths, as well as the recruitment of eosinophils, basophils, and mast cells to the sites of infections. This cluster had higher expression of *CCR4, GATA3, IL17RB* and *IL4IL* (Cano-Gamez et al., 2020; Stark et al., 2019).

**Th17:** Th17 cells mediate responses for the clearance of extracellular bacteria and fungi by orchestrating sustained neutrophil recruitment. This cluster had higher expression of *CCR6, RORC, IL17A* and *IL17F* (Ma et al., 2016).

**Th17/Th1:** Th17 cells can convert to Th1 cells based on the cytokine environment. This cluster had expressions of *IFNG*, *IL26* and *IL17A* (Damsker et al., 2010).

**Th22:** Th22 cells are players in adaptive immune responses and are identified by the production of IL-22. This cluster had higher levels of *IL22, IL10* and *TNF* (Jia and Wu, 2014).

**Eff-Tregs:** Effector T regulatory cells play an important role in maintaining the homeostasis in tissues and control immune responses. This cluster is the classical Treg subpopulation that expresses IL2RA, CTLA4, IL1R1, IL1R2 and IL10 and specific transcription factors FOXP3, MAF, IKZF1 and IKZF3 and it is CD25+ and CD28+ (Du et al., 2021).

**Eff-Tregs-IL32:** This cluster had a similar profile of Eff-Tregs, it showed a distinct transcriptional signature, with overexpression of several suppressive effectors, but also proinflammatory molecules like IL32. This cluster had as top genes: *IL32, FOXP3, CTLA4, IL2RA* and *IKZF2* (Galván-Peña et al., 2021).

Tfr: T-follicular regulatory cells in the tonsils are CD25-. These cells down-regulate effector Treg markers, and acquire a follicular T helper profile. This cluster had lower levels of FOXP3 and upregulation of BCL6, PDCD1, ICOS and TCF7*.* This cluster was CD25- (Wing et al., 2017).

### CD8+ T CELLS AND INNATE LYMPHOID CELLS

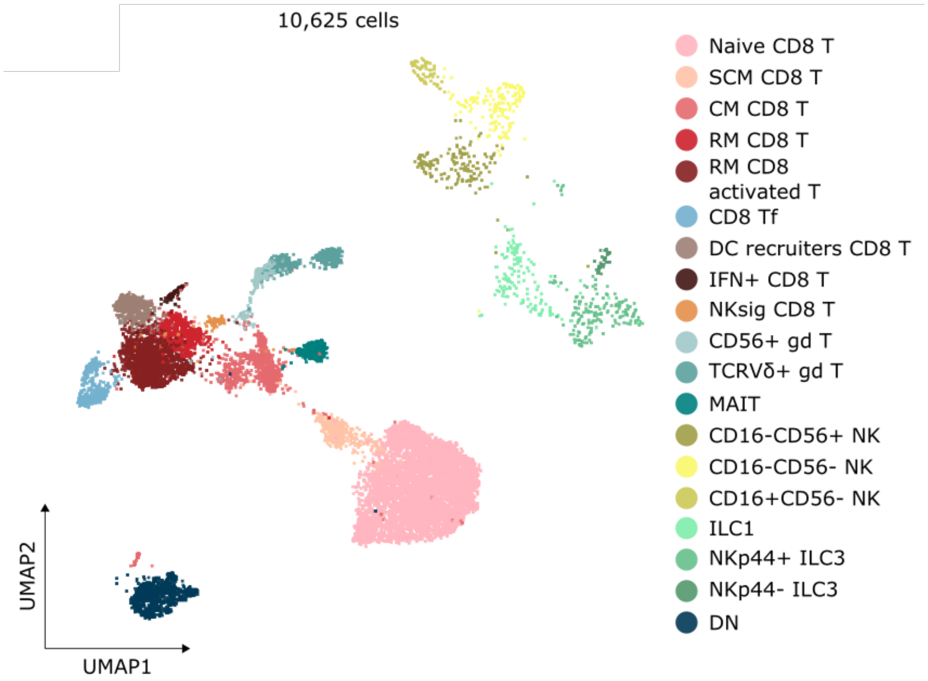


**Naïve CD8:** Naïve cell is a CD8 T cell that recently migrates from the thymus towards lymph nodes. This cluster had an expression of *BACH2, LEF1, NOSIP* and *CCR7*. Also, this cluster was CD45RA+.

**SCM CD8 T:** Stem-cell memory T cells self-renew and generate long-lived central memory T cells and short-lived effector memory T cells. Differential expression analysis reveals these genes as the main markers: *FAM117B, TCF7* and *PLAC8*. This population was CD95+ and CXCR3+ (Galletti et al., 2020).

**CM CD8 T:** After antigen encounter T cells acquire memory and effector functions maintaining the lymph node homing. CM showed an upregulation of *IL7R* compared to naïve T cells, and higher expression of *ANXA1* and *S100A4*. They had an intermediate expression of CCR7 and a higher expression of *CD28*. This cluster was CD45RO+, and CD28+.

**RM CD8 T:** Resident memory T cells are essential for immune defense against pathogens and malignancies. RM CD8 T cell is a lineage, resulting in the generation of a pool of TRM cell- poised T cells within the tonsil compartment. The main gen markers of this cluster are: *CD69, EOMES, ZEB2, ITGA1* and *TOX,* and CD69+ (Kok et al., 2022).

**RM CD8 Activated T:** This cluster had a similar profile of RM T cells but it had higher expression of *GZMH, CCL5, GZMA, GZMK, CXCR6, HLA-DRB1, HLA-DPA1* and *CD38.* These markers showed an activated profile of these RM CD8 T cells. Also, It was positive for CD103 and CD54.

**CD8 Tf:** CD8+ T cells located within the lymphoid follicle are follicular CD8+ T cells that play an important role in viral and tumor control, as well as to modulate humoral and T follicular helper cell responses. This cluster had upregulated CD200, CXCL13, ICOS, PDCD1 and IFNG, and it was CXCR5+, PD-1+ and ICOS+ (Perdomo-Celis et al., 2017).

**DC recruiters CD8 T:** CD8+ T cells can orchestrate the local recruitment of tonsil resident XCR1+ DCs via secretion of the XCL1 chemokine. Functionally, this CD8+ T cell-mediated reorganization of the local DC network allowed for the interaction and cooperation of pDCs and XCR1+ DCs. This cluster had as top markers *CCL5, XCL2, XCL1* and *CCL4*. These markers confirmed their role in the recruitment of DCs (Brewitz et al., 2017).

**IFN+ CD8 T:** This cluster had an expression of genes related with IFNg responses. The up-regulated genes were *HLA-DRB1, IFIT3, IFIT5* and *IFITM2.*

**NKsig CD8 T:** Terminally differentiated CD8 effector T cells can have NK-like features and they can be potent mediators of viral responses. This cluster had specific markers related to an NK signature. These were: *GNLY, NKG7, CX3CR1* and *FCGR3A*. This cluster was CD16+, CD158b+ and CD56+ (Mathewson et al., 2021).

**TCRVd gd-T:** γδ T cells are a distinct subgroup of T cells containing T cell receptors γ and TCR δ chains. In this cluster we identified a clear signature of gd T cells: *KLRD1, KLRC2, TRGC2* and *TRDC* (Pizzolato et al., 2019).

**CD56+ g/d T:** This cluster had a similar signature of gd T cells but expresses specific markers of NK cells. This cluster had the expression of *NCAM1, KLRC2, ZNF683,* and *XCL1* and was positive for CD56 (Nörenberg et al., 2021).

**MAIT:** Mucosal- associated invariant T cells, had higher expression of *KLRB1, IL7R, RORA* and *IL18RAP* (Pellicci et al., 2020).

**CD16- CD56-:** Immature NK-cells, without low cytotoxic activity, express *SELL,* and CD161 and CD122, but not CD16 or CD56 (Boldt et al., 2014).

**CD16-CD56+:** NK cells with low cytolytic activity. This cluster had an expression of *NCAM* and higher levels of CD56 (Boldt et al., 2014).

**CD16+ CD56-:** An intermediate NK cell subtype with migratory capacity. This cluster had an expression of *CX3CR1, FCGR3A,* and *TBX21.*

**ILC1:** ILC1 are characterized by their capacity to produce IFNγ and they depend on the transcription factor T- bet. This cluster had higher levels of *CD200R1, CD69* and *ANXA1* (Boldt et al., 2014).

**NKp44+ ILC3 and NKp44- ILC3:** ILC3s depend on RORC for their development and function, and comprise subsets that can be distinguished on the basis of expression of the natural cytotoxicity receptors NKp46 and NKp44. The main cytokine produced by ILC3s is IL-22 and a proportion of ILC3s also produce IL-17. We identified two subtypes of ILC3s that expressed RORC and were NKp44 positive and negative (Bal et al., 2020).

**DN:** double negative T cells express the αβ T cell receptor but do not express CD4, CD8, or natural killer cell markers. This cluster had a profile of inflammatory and activated DN T cells expressing *HLA-DRB1, HLA-DPA1, CD38* and *GPR183* (Wu et al., 2022b).

**Cycling T:** This cluster is a T cell cluster (CD3+) that had a higher score of cell cycle and had higher expression of cell cycle genes: *STMN1, HIST1H4C, TUBA1B, HMGB2, MKI67, TOP2A* and *TUBB*. As we observed that the proliferative signature was a major driver of variance in scRNA-seq data, which caused (1) cells from different lineages to cluster together and (2) hindendered biological heterogeneity in the other cluster. Thus we show this population in Figure 1B, but exclude it from downstream analysis.

### NAIVE AND MEMORY CELLS

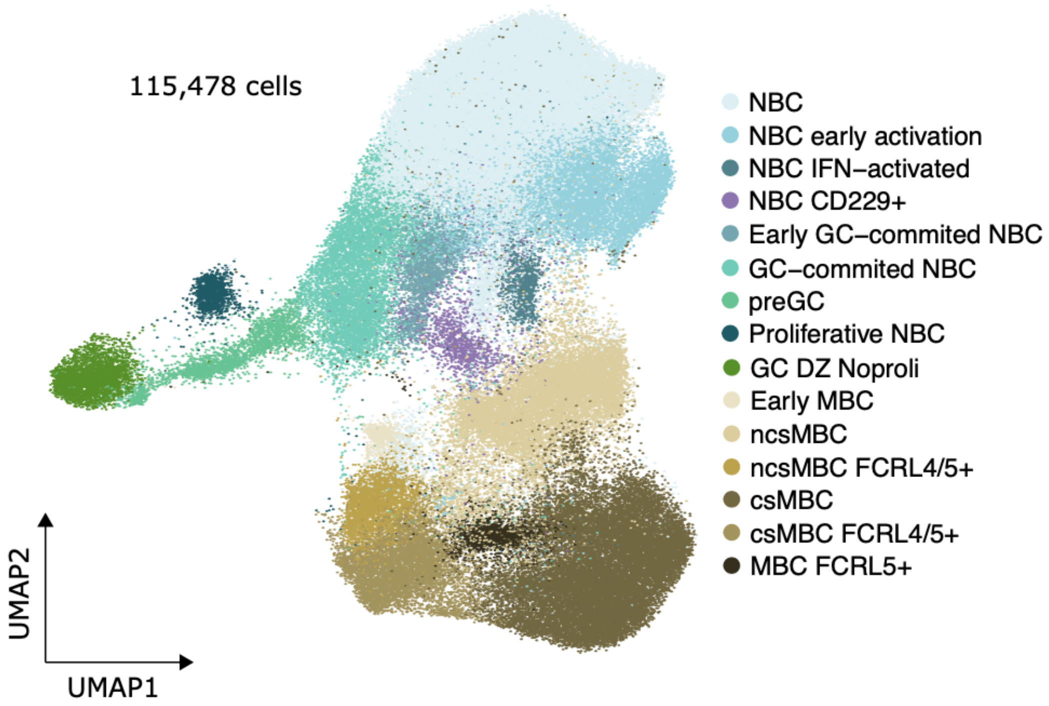


* For the sake of mapping cell transitions, in this analysis we included *DZ non proliferative GCBCs* from the GCBC level 1 cluster. This subpopulation is not described in this section of the glossary but in the GCBC section.

**NBC**: This is the main cluster of NBCs, with a clear and homogenous expression of FCER2 (CD23), TCLA1, IGHM and IGHD. This cluster seems to represent standard NBC lacking the expression of genes related to cell activation or commitment towards GC reaction. We would like to clarify that we performed a targeted search for two known NBC subpopulations that are expected to be present in tonsillar tissue, i.e. CD5 Mature NBCs and transitional B cells. However, CD5, a key marker to define these subpopulations, could not be detected by scRNAseq. We also used alternative gene signatures defined by Seifert et al. (Seifert et al., 2012) but we could not detect any particular subcluster. Therefore, although we cannot rule out that CD5 Mature NBCs and transitional B cells are present in our dataset -probably within the standard NBC cluster-, we could not identify them.

**NBC early activation**: Cells in this cluster show expression of typical NBC markers such as FCER2 (CD23), TCLA1, IGHM and IGHD. This cluster is transcriptionally very similar to standard NBCs, with the exception of a few genes related to B cell activation, such as CD69 and IL4R (Cibrián and Sánchez-Madrid, 2017; Ferrer et al., 2014). Therefore, these cells seem to represent the first step of NBC activation.

**NBC INF-activated**: Cells in this cluster show expression of typical NBC markers such as FCER2 (CD23), TCLA1, IGHM and IGHD. Several genes expressed exclusively by the cells in these clusters are associated with interferon induction, as for example ISG15, the IFIT gene family (i.e. or IFIT1, IFIT2 or IFIT3) family, IFI44, IFI44L and MX1. Thus, this cluster represents NBC cells that have been activated in response to interferon stimulation.

**NBC CD229+**: Cells in this cluster show expression of typical NBC markers such as FCER2 (CD23), TCLA1, IGHM and IGHD. As compared to the standard NBC cluster, only few genes were differentially expressed. We identified two main markers, related to immune system and B cell biology that were specifically expressed within this cluster, i.e. LY9 (CD229), which has been shown to be highly expressed in marginal zone B cells (Cuenca et al., 2016) and LILRA4 (Cao and Bover, 2010), which actually is more relevant for plasmacytoid dendritic cells. Unfortunately, the lack of literature related to this particular combination of markers precluded us from assigning a biological function to this cluster.

**Early GC-committed NBC:** Cells in this cluster show expression of typical NBC markers such as FCER2 (CD23), TCLA1, IGHM and IGHD. This cluster is characterized by the expression of very specific genes that seem to correspond to early GC commitment due to the transient MYC upregulation (Dominguez-Sola et al., 2012). For instance, we observed high expression of EGR family (i.e. EGR1, EGR2, EGR3), transcription factors that are rapidly induced upon stimulation of the BCR; genes related to NF-kB pathway (i.e. NFKB1, NFKB2, NFKBIA, NFKBID, RELB); or proinflammatory chemokines i.e. CCL3 and CCL4. Furthermore, in this cluster we identified high levels of bone-fide activation marker, i.e. CD69. As this marker was also present at lower levels in the NBC early activation cluster, we may hypothesize that these two clusters represent sequential steps of the NBC activation process towards the GC reaction.

**GC-committed NBC**: Cells in this cluster show expression of typical NBC markers such as FCER2 (CD23), TCLA1, IGHM and IGHD. This cluster seems to follow the Early GC-committed NBCs in the differentiation path between NBC and GCBCs. For instance, one of the main markers of this cluster is CCND2, which is a bona fide downstream target of MYC (a marker for early GC commitment, (Dominguez-Sola et al., 2012). It also maintains the expression of NF-kB pathway related genes. Furthermore, the cells in this cluster express genes associated with signaling pathways related to response to antigens, such as MIR155HG, TRAF1/4, EBI3, and BCL2A1. Finally, we observed upregulation of BATF, a gene known to be involved in the regulation of AID (AICDA) expression and thus antibody diversification in DZ-GCBCs (Ise et al., 2011). Although this cluster lacks any GC-related marker, it seems to represent a cell stage that is already committed to enter the GC reaction.

**preGC**: Cells in this cluster show expression of typical NBC markers such as FCER2 (CD23), TCLA1, IGHM and IGHD. As compared to the preceding clusters related to NBC activation, within this cluster we could detect the expression of GC-specific genes, such as RGS13 or MEF2B. However, these cells still partially maintain the NBC transcriptome and transcriptional programs started in the previous steps of NBC activation process (i.e. GC-committed NBC) such as the BATF expression. Additionally, this cluster still lacks other GC phenotypic genes, such as AICDA or BCL6. Therefore, we conclude that this subcluster may represent the last step of NBC activation before entering into the DZ and starting a proper GC reaction.

**Proliferative NBC:** Cells in this cluster show expression of typical NBC markers such as FCER2 (CD23), TCLA1, IGHM and IGHD. This cluster is characterized by a proliferation signature, marked by high expression of MKI67, TOP2A, histone genes and genes related with chromosome segregation. Intriguingly, this cluster does not contain any marker related to GC, i.e. lack of expression of RGS13, MEF2B (the two expressed in preGC cluster), AICDA and BCL6. We therefore discarded that these cells were related to the GC-dependent maturation pathway of NBCs. Instead, we hypothesize that this population may correspond to primary focal reaction upon antigen stimulation (Jacob and Kelsoe, 1992), which leads to rapid generation of antibody-producing cells in a GC-independent manner.

**Early MBC:** This cluster of the NBC/MBC level 1 cluster is CD27 positive, and therefore, it seems to be a MBC. However, this cluster does not have many specific markers, with the exception of e.g. low-medium levels of MEF2B, a GC marker. This might suggest that this cluster represents MBCs freshly generated from the GC reaction, and still contains traces of GC markers such as MEF2B.

**ncsMBC:** This is one of the main clusters of MBCs, with clear expression of known markers such as CD27 and CD267. It is also characterized by high expression of IGHD and IGHM, and therefore, this cluster is classified as ncsMBCs.

**csMBC**: This is one of the main clusters of MBCs, with clear expression of known markers such as CD27 and CD267. It is also characterized by high expression of IGHA1 and IGHG1, and therefore it is classified as csMBCs.

**ncsMBC FCRL4/5+:** This is a subcluster of ncsMBC (positive for CD27, CD267, IGHD and IGHM), characterized by the expression of FCRL4 and FCRL5. This subpopulation is typically found in tonsils but not in peripheral blood or bone marrow. In concordance with the available literature related to FCRL4+ MBCs, this cluster expresses high levels of GSN, SOX5, FGR or HCK, among others (Li et al., 2020).

**csMBC FCRL4/5+:** This is a subcluster of csMBC (positive for CD27, CD267, IGHA and IGHG), characterized by the expression of FCRL4 and FCRL5. This subpopulation is typically found in tonsils but not in peripheral blood or bone marrow. In concordance with the available literature related to FCRL4+ MBCs, this cluster expresses high levels of GSN, SOX5, FGR or HCK, among others (Li et al., 2020).

**MBC FCRL5+**: This is a low abundant cluster of MBC cells (positive for CD27 and CD267) that present and intermediate transcriptomic signature between csMBC and csMBC FCRL4+/FCRL5+. These cells express only FCRL5 with the absence of FCRL4. This subpopulation has been described as MBC FCRL5-only, which indeed was shown to express many genes at intermediate level between FCRL4/FCRL5 MBCs and MBC (Li et al., 2020).

### GERMINAL CENTER B CELLS

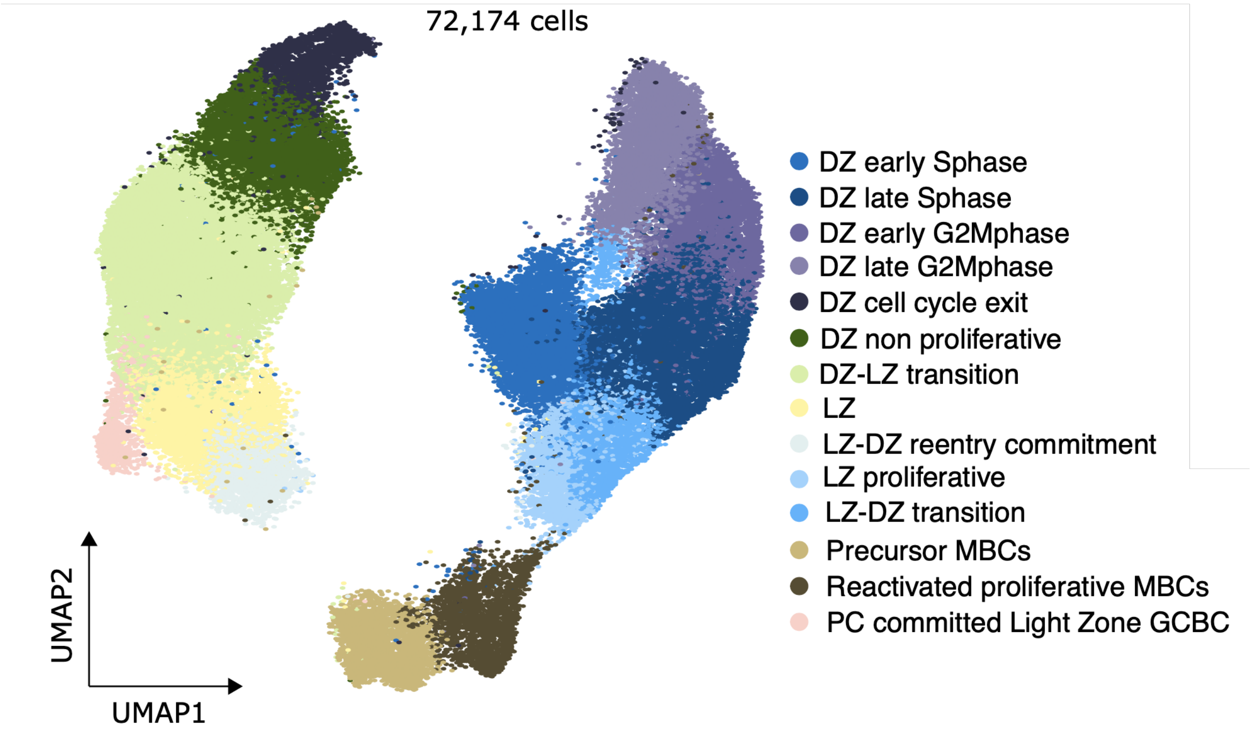


*The *PC committed light zone GCBC* subpopulation will be described in the plasma cell section.

**DZ-early S phase:** Cluster with high expression of CXCR4 (a marker defining DZ GCB cells) and additional expression of DZ markers, such as FOXP1 and MME, and moderate levels of AICDA. This cluster displays a clear proliferation signature and high S-phase score. Cells express typical cell cycle genes, such as MCM family, POLA1 or PCNA, which are associated with DNA replication and strand elongation as well as G1/S transition GO terms. This cluster lacks expression of histone genes (i.e. HIST1H4C, HIST1H1B, HIST2H2AH, HIST1H3D), which are related to a more advanced S phase. Therefore, the present cluster most likely represents the early steps of the S phase of the cell cycle.

**DZ-late S phase**: Cluster with high expression of CXCR4 (a marker defining DZ GCB cells) and additional expression of DZ markers, such as FOXP1 and MME, and moderate levels of AICDA. This cluster displays a clear proliferation signature and high S-phase score. Cells express typical cell cycle genes, such as MCM family, POLA1 or PCNA, which are associated with DNA replication and strand elongation as well as G1/S transition GO terms. This cluster shows upregulation of histone genes (i.e. HIST1H4C, HIST1H1B, HIST2H2AH, HIST1H3D), suggesting that it contains cells associated with the later steps of the S phase. Moreover, the cells within this cluster start expressing the first traces of genes related to more advanced phases of the cell cycle, such as AURKB, TOP2A or MKI67.

**DZ-early G2M phase**: Cluster with high expression of CXCR4 (a marker defining DZ GCB cells) and additional expression of DZ markers, such as FOXP1 and MME, and intermediate levels of AICDA. This cluster has a clear G2M phase score. It is characterized by high expression of genes related to spindle organization, chromosome segregation, mitotic cell cycle and division, i.e. BUB1, TOP2A, centromere proteins family, tubulin genes, AURKA, CDK1 and MKI67. However, cells from this cluster still have high expression of histone genes and do not present the upregulation of cyclin B and CDC20, suggesting that they are in early G2M phase.

**DZ-late G2M phase**: Cluster with high expression of CXCR4 (a marker defining DZ GCB cells) and additional expression of DZ markers, such as FOXP1 and MME, as well as high expression of AICDA. This cluster has a clear G2M phase score. It is characterized by high expression of genes related to spindle organization, chromosome segregation, mitotic cell cycle and division, i.e. BUB1, TOP2A, centromere proteins family, tubulin genes, AURKA, CDK1 and MKI67. Cells from this cluster start to downregulate the expression of histone genes concomitantly with the upregulation of cyclin B genes and CDC20, indicating full entry into mitosis.

**DZ-cell cycle exit**: Cluster with high expression of CXCR4 (a marker defining DZ GCB cells) and additional expression of DZ markers, such as FOXP1 and MME, as well as moderate expression of AICDA. Cells from this cluster start to downregulate the expression levels of cell cycle program genes. For instance, they lack CDK1 but still contain lower levels of CCNB1, CDC20, TOP2A and MKI67 as compared to the cells from preceding DZ-G2M phase clusters. The cells still maintain clear expression of other proliferation-related genes such as HMG family genes, STMN1 or BIRC5. This cluster was not associated with any significant upregulation as compared to the preceding cluster (DZ-late G2M phase). Therefore, we defined this cluster as dark zone cells that have started to exit the cell cycle.

**DZ-non-proliferative:** Cluster with high expression of CXCR4 (a marker defining DZ GCB cells) and additional expression of DZ markers, such as FOXP1 and MME, as well as moderate expression of AICDA. Cells from these clusters are clearly non-proliferative, lacking the detectable expression of any cell cycle gene. Therefore, we hypothesize that cells in this cluster, although they maintain the expression of dark zone-related genes, have already finished the cell cycle program and are progressing towards the next maturation stage.

**DZ-LZ transition:** This relatively large non-proliferative cluster is characterized by rather intermediate or low expression of DZ markers, such as CXCR4. The cells start to express some LZ markers such as CD83 or LMO2, but still are missing other LZ-genes such as BCL2A1 or DUSP2. Furthermore, we could not define clear markers specific and unique for this cluster. From the chromatin perspective, this cluster contained intermediate levels of regions differentially accessible between the DZ and the LZ. Therefore, we believe that these cells represent a transitional subpopulation moving from the DZ to the LZ.

**LZ**: Cells in this non-proliferative cluster show expression of well-known light zone markers, i.e. CD83, LMO2, and downregulation of DZ genes such as CXCR4 or AICDA. Within this cluster, genes related to CD40 signaling and response to BCR stimulation are starting to be expressed (e.g. BCL2A1, BCL2L1, TRAF1/4 or EBI3). However, the expression levels of these genes were not homogeneous, suggesting the presence of different levels of activation of LZ cells.

**LZ-DZ re-entry commitment**: Cluster with high expression of CD83 and low expression of CXCR4, as well as positive for LMO2, which indicate that this cluster contains LZ-GCB cells. Within this cluster, apart from the high expression of genes associated with the previously described LZ cluster, the cells start to express very specific genes such as MYC, MIR155HG or genes related to NF-kB pathway (i.e. NFKB1, NFKB2, NFKBIA, NFKBID, RELB). This transient expression of MYC has been described in LZ cells re-entering into the DZ, to continue with additional rounds of affinity maturation (Dominguez-Sola et al., 2012). Additionally, as compared to the LZ cluster, this cluster shows higher expression of BATF, a gene known to be involved in regulating AIDCA (a DZ marker) (Ise et al., 2011). From the scATAC-seq perspective, this LZ-DZ re-entry cluster is associated with epigenetic programming, with transiently increased chromatin accessibility associated with TFs of the NF-kB family as well as BATF. Altogether, these observations suggest that this cluster contains LZ cells that are already committed and preparing to return to the DZ.

**LZ-proliferative:** This cluster has a proliferative signature, marked for example by upregulation of MCM gene family, PCNA, POLA1, while still maintaining the expression of standard light zone markers, like CD83 and LMO2. Furthermore, these cells maintain the expression of specific genes related to the LZ-DZ re-entry commitment cluster, such as BCL2A1, MIR155HG, NF-kB pathway genes or BATF. Therefore, this cluster may represent LZ cells that are starting to proliferate.

**LZ-DZ transition**: This proliferative cluster presents intermediate expression of both LZ and DZ markers, i.e. containing traces of both CD83 and CXCR4 expression. It also downregulates previously mentioned signature related to LZ-DZ re-entry commitment, but cell cycle genes are highly expressed. Therefore, these cells may represent the intermediate transitional step from proliferative LZ to proliferative DZ cells. It also suggests that the phenotypic switch from LZ to DZ occurs while cells proliferate.

**Precursor MBC**: Within the level 1 cluster containing GCBCs, we also identified a subset of cells that were characterized by MBC-related markers while maintaining a non-proliferative GC signature. Specifically, this cluster was characterized by high expression of CCR6 or CELF2, genes that are commonly used in literature to define memory B cell precursors (Holmes et al., 2020). Although this cluster likely represents early MBCs derived from the LZ, we cannot exclude some technical issues or even MBC-GC doublet cells.

**Reactivated proliferative MBC**: This is a second cluster within the GC compartment that contains MBC markers. In this case, cells show high expression of BANK1 and TNFRSF13B (CD267). However, in the contrary to precursor MBC (that supposedly are generated from non-proliferative LZ cells), this cluster is characterized by high proliferation signature (i.e. PCNA, MKi67 or TOP2A expression). Therefore, we hypothesize that these cells may be derived from secondary immune responses, in which MBC are reactivated, start to rapidly proliferate and may eventually differentiate into plasma cells. However, the overlap between some GC and MBC markers does not allow us to unequivocally exclude doublets.

### PLASMA CELLS

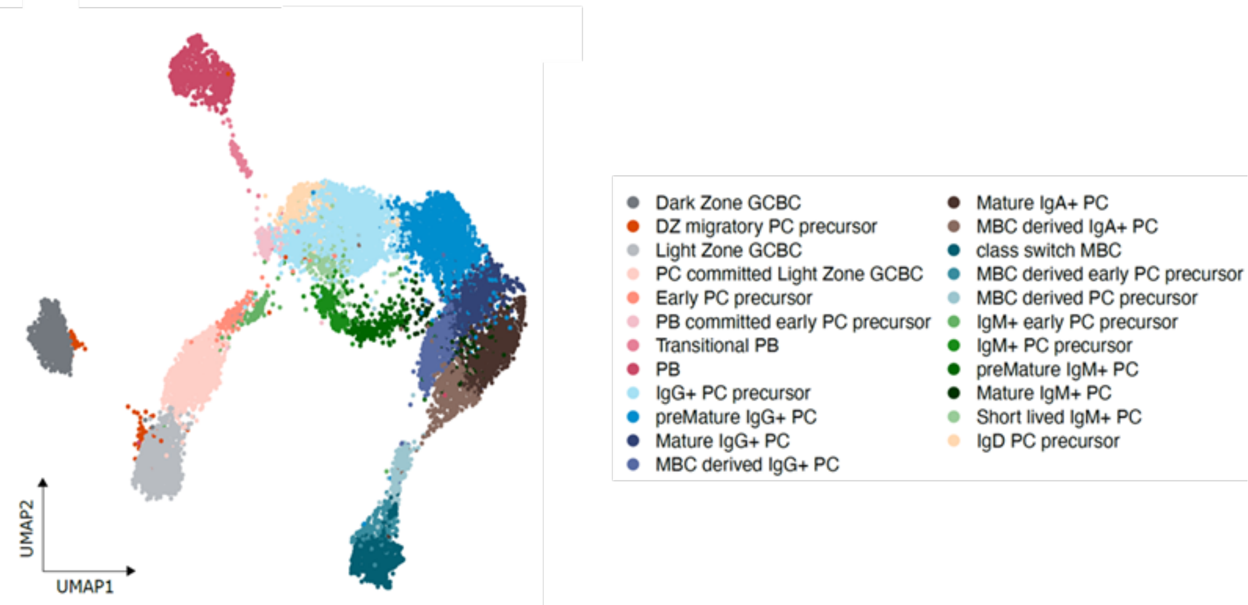


*For the sake of mapping cell transitions, in this analysis we included *Dark-Zone* and *Light-Zone GCBCs* as well as *class-switched MBCs* from the GCBC and NBC/MBC level 1 clusters, respectively. These subpopulations will not be described in this section of the glossary but in their respective clusters.

**DZ migratory PC precursor:** This cluster is composed by a relatively small number of cells that belong to the PC cluster at level 1, but in the PC-specific UMAP unexpectedly cluster close to both non-proliferative DZ-GCBCs and LZ-GCBCs (which were borrowed from the GCBC cluster). This intriguing cluster shows expression of GC markers such as BCL6, IRF8 and MEF2B, as well as intermediate levels of DZ (e.g.CXCR4 and AICDA) and LZ markers (e.g. CD83 and LMO2). Additionally, expression of B cell markers such as MS4A1 and PAX5 can be observed. These observations may be consistent with published evidence in mice suggesting that there are some maturing PCs migrating from the LZ to the DZ before leaving the GC (Kräutler et al., 2017). Although this hypothesis is attractive, we cannot formally rule out that these cells may represent doublets from PCs and GCBCs.

**PC committed light zone GCBC:** This subpopulation still belongs to the GCBCs level 1 cluster (it is described within this plasma cell section of the glossary). It maintains high levels of B cell markers such as MS4A1 and PAX5, as well as high levels of GC markers such as BCL6, IRF8 and MEF2B, and LZ markers (MME (CD10), and CD83). However, it shows the first seeds of commitment towards PC differentiation, which are associated mostly with the expression, still at low levels, of the three key PC-related transcription factors PRDM1, XBP1 and IRF4.

**Early PC precursor**: This cluster belongs to the PC level 1 cluster, and is located next to the PC-committed LZ-GCBCs. Although it still maintains the expression of B cell and GC markers, their levels are reduced as compared to the PC committed cells (e.g. MS4A1, BCL6, IRF8, CD83). On the contrary, the expression of the 3 PC transcription factors (PRDM1, XBP1 and IRF4) are increased as compared to PC committed cells, and the first traces of PC phenotypic markers can be detected, e.g. MZB1, SLAMF7, SSR4, CREB3L2 and FKBP11.

**PB committed early PC precursor:** Similar expression profile to the Early PC precursors, with low levels of B cell and GC markers (e.g. MS4A1, BCL6, IRF8, CD83), high expression expression levels of the 3 PC transcription factors (PRDM1, XBP1 and IRF4) and traces of PC phenotypic markers, e.g. MZB1, SLAMF7, SSR4, CREB3L2 and FKBP11. CD9, a marker for precursor PCs (Yoon et al., 2013), can be detected. However, in this cluster, low levels of some cell division-related genes such as H2AFZ and TUBA1B. Therefore, we interpret that these early PC precursors are already committed to become proliferative plasmablasts.

**Transitional PB**: Cells in this cluster express the 3 PC transcription factors (PRDM1, XBP1 and IRF4) and PC phenotypic markers such as MZB1, SLAMF7, SSR4, CREB3L2 and FKBP11. CD9, a marker for precursor PCs (Yoon et al., 2013), can be detected. This is a peculiar cluster with intermediate levels of genes characteristic for proliferative plasmablasts such as H2AFZ, TUBA1B, MKI67 and TOP2A. An analysis of cell cycle phases indicates that this cluster has cells both in S phase and G2M phase, which may suggest that it contains cells transitioning from PB committed early PC precursors to proliferative PBs and vice versa.

**PB:** Cells within this cluster are clearly proliferative plasmablasts with high expression levels of proliferation-related genes (H2AFZ, TUBA1B, MKI67 and TOP2A, among others). Otherwise have an expression pattern of PCs (the 3 transcription factors PRDM1, IRF4 and XBP1, as well as phenotypic markers MZB1, SLAMF7, SSR4, CREB3L2 and FKBP11). CD9, a marker for precursor PCs (Yoon et al., 2013), can be detected. As compared to transitional PBs, the expression levels of PC phenotypic markers are slightly increased. Traces of B cell markers can still be detected (e.g. MS4A1 and PAX5) and even some GC-related markers such as MEF2B.

**IgG+ PC precursor**: Cells within this cluster have a typical PC markers (the 3 transcription factors PRDM1, IRF4 and XBP1, as well as phenotypic markers MZB1, SLAMF7, SSR4, CREB3L2 and FKBP11). CD9, a marker for precursor PCs (Yoon et al., 2013), can be detected.. Although the typical B cell-related genes (MS4A1 and PAX5, among others) are downregulated, some traces of MEF2B, a GC marker, can still be observed. Finally, as compared to early precursors, higher expression levels of IGHG can be detected in this cluster.

**PreMature IgG+ PC:** Cells within this cluster have a typical PC markers (the 3 transcription factors PRDM1, IRF4 and XBP1, as well as phenotypic markers MZB1, SLAMF7, SSR4, CREB3L2 and FKBP11). CD9, a marker for precursor PCs (Yoon et al., 2013) can still be detected. Although the typical B cell-related genes (MS4A1 and PAX5, among others) are downregulated, some traces of MEF2B, a GC marker, can still be observed. Finally, as compared to IgG+ PC precursors, higher expression levels of IGHG can be detected in this cluster.

**Mature IgG+ PC**: Cells within this cluster have a typical PC markers (the 3 transcription factors PRDM1, IRF4 and XBP1, as well as phenotypic markers MZB1, SLAMF7, SSR4, CREB3L2 and FKBP11). CD9, a marker for precursor PCs (Yoon et al., 2013) is downregulated, and CD44, a marker for mature PCs (Khodadadi et al., 2019), is expressed. Traces of MEF2B, a GC marker, or typical B-cell related genes (e.g. MS4A1) cannot be detected anymore. Finally, this subpopulation shows high expression levels of IGHG.

**MBC-derived IgG+ PC:** Cells within this cluster have a typical PC markers (the 3 transcription factors PRDM1, IRF4 and XBP1, as well as phenotypic markers MZB1, SLAMF7, SSR4, CREB3L2 and FKBP11). CD9, a marker for precursor PCs (Yoon et al., 2013) is downregulated, and CD44, a marker for mature PCs (Khodadadi et al., 2019), is expressed. This subpopulation shows high expression levels of IGHG. Remarkably, cells in this cluster show expression of some markers expressed in MBCs, and in MBC-derived precursor PCs, such as CXCR4, FCMR and CCDC50. Therefore, the mature IgG+ PC phenotype together with traces of MBC-related genes made us think that this cluster represents mature IgG+ PCs derived from the reactivation of MBCs.

**Mature IgA+ PC**: Cells within this cluster have a typical PC markers (the 3 transcription factors PRDM1, IRF4 and XBP1, as well as phenotypic markers MZB1, SLAMF7, SSR4, CREB3L2 and FKBP11). CD9, a marker for precursor PCs (Yoon et al., 2013) is downregulated, and CD44, a marker for mature PCs (Khodadadi et al., 2019), is expressed. Traces of MEF2B, a GC marker, or typical B-cell related genes (e.g. MS4A1) cannot be detected anymore. Finally, this subpopulation shows high expression levels of IGHA.

**MBC-derived IgA+ PC:** Cells within this cluster have a typical PC markers (the 3 transcription factors PRDM1, IRF4 and XBP1, as well as phenotypic markers MZB1, SLAMF7, SSR4, CREB3L2 and FKBP11). CD9, a marker for precursor PCs is downregulated, and CD44, a marker for mature PCs, is expressed. This subpopulation shows high expression levels of IGHA. Remarkably, cells in this cluster show expression of some markers expressed in MBCs, and in MBC-derived precursor PCs, such as CXCR4, FCMR and CCDC50. Therefore, the mature IgA+ PC phenotype together with traces of MBC-related genes made us think that this cluster represents mature IgA+ PCs derived from the reactivation of MBCs.

**MBC-derived early PC precursor**: This subpopulation clusters next to the csMBCs that we borrowed from the MBC cluster at level 1. As compared to csMBCs, these cells show low levels of PC related genes (PRDM1, IRF4, XBP1, MZB1, SLAMF7, SSR4, CREB3L2 and FKBP11), and increased levels of IG genes expression (although still low). Traces of B cell related genes are expressed such as MS4A1 and PAX5. Additionally, MBC-related genes are also expressed (i.e. CXCR4, FCMR and CCDC50). Therefore, we interpret that this subpopulation represents the early PC precursors derived from the reactivation of MBCs.

**MBC-derived PC precursor:** This subpopulation clusters between MBC-derived early PC precursor and mature PCs. As compared to MBC-derived early PC precursor, these cells show higher levels of PC related genes (PRDM1, IRF4, XBP1, MZB1, SLAMF7, SSR4, CREB3L2 and FKBP11), and increased levels of IG genes expression. Traces of B cell related genes are expressed such as MS4A1 and PAX5. Additionally, MBC-related genes are also expressed (i.e. CXCR4, FCMR and CCDC50). Therefore, due to the increased expression of PC and IG genes as compared to the MBC-derived early PC precursor, we interpret that this cluster represents a further step towards the phenotypic transformation of a MBC into a PC.

**IgM+ early PC precursor**: This cluster maps close to the early PC precursor cells but consistently show low expression levels of IGHM. The global gene expression pattern is similar to early PC precursor cells. It maintains low expression levels of B cell and GC markers, (e.g. MS4A1, BCL6, IRF8, CD83), and shows expression of the 3 PC transcription factors (PRDM1, XBP1 and IRF4) and the first traces of PC phenotypic markers can be detected, e.g. MZB1, SLAMF7, SSR4, CREB3L2 and FKBP11.

**IgM+ PC precursor:** Cells within this cluster express IGHM and have a typical PC markers (the 3 transcription factors PRDM1, IRF4 and XBP1, as well as phenotypic markers MZB1, SLAMF7, SSR4, CREB3L2 and FKBP11). CD9, a marker for precursor PCs (Yoon et al., 2013), can be detected. Although the typical B cell-related genes (MS4A1 and PAX5, among others) are downregulated, some traces of MEF2B, a GC marker, can still be observed. Finally, as compared to IgM+ early PC precursor, higher expression levels of IGHM can be detected.

**Premature IgM+ PC**: This IGHM-positive cluster show the expression of typical PC markers (the 3 transcription factors PRDM1, IRF4 and XBP1, as well as phenotypic markers MZB1, SLAMF7, SSR4, CREB3L2 and FKBP11). CD9, a marker for precursor PCs (Yoon et al., 2013) is downregulated, and the expression of CD44, a marker for more mature PCs (Khodadadi et al., 2019), is increased. Finally, as compared to IgM+ PC precursor, higher expression levels of IGHM can be detected .

**Mature IgM+ PC**: Cells within this cluster express IGHM and have a typical PC markers (the 3 transcription factors PRDM1, IRF4 and XBP1, as well as phenotypic markers MZB1, SLAMF7, SSR4, CREB3L2 and FKBP11). CD9, a marker for precursor PCs (Yoon et al., 2013) is downregulated. Traces typical B-cell related genes (e.g. MS4A1) cannot be detected. Finally, among the IgM+ clusters, this subpopulation shows the highest expression levels of IGHM.

**Short lived IgM+ PCs**: Cells within this cluster express IGHM and have a typical PC markers (the 3 transcription factors PRDM1, IRF4 and XBP1, as well as phenotypic markers MZB1, SLAMF7, SSR4, CREB3L2 and FKBP11). CD9, a marker for precursor PCs (Yoon et al., 2013), can also be detected. Although relatively similar to IgM+ PC precursor, this cluster shows a global gene downregulation with the exception of few upregulated genes strongly enriched in the endoplasmic reticulum cellular component concomitantly to high IgM expression. Therefore, we conclude that this may represent a subpopulation of short lived IgM PC cells that shut down most of the cellular functions with the exception of high production of IgM which is processed through the endoplasmic reticulum.

**IgD+ PC precursors**: Cells within this cluster express IGHD and have a typical PC markers (the 3 transcription factors PRDM1, IRF4 and XBP1, as well as phenotypic markers MZB1, SLAMF7, SSR4, CREB3L2 and FKBP11). CD9, a marker for precursor PCs (Yoon et al., 2013), can be detected. Although the typical B cell-related genes (MS4A1 and PAX5, among others) are downregulated, some traces of MEF2B, a GC marker, can still be observed.

#### MYELOID CELLS

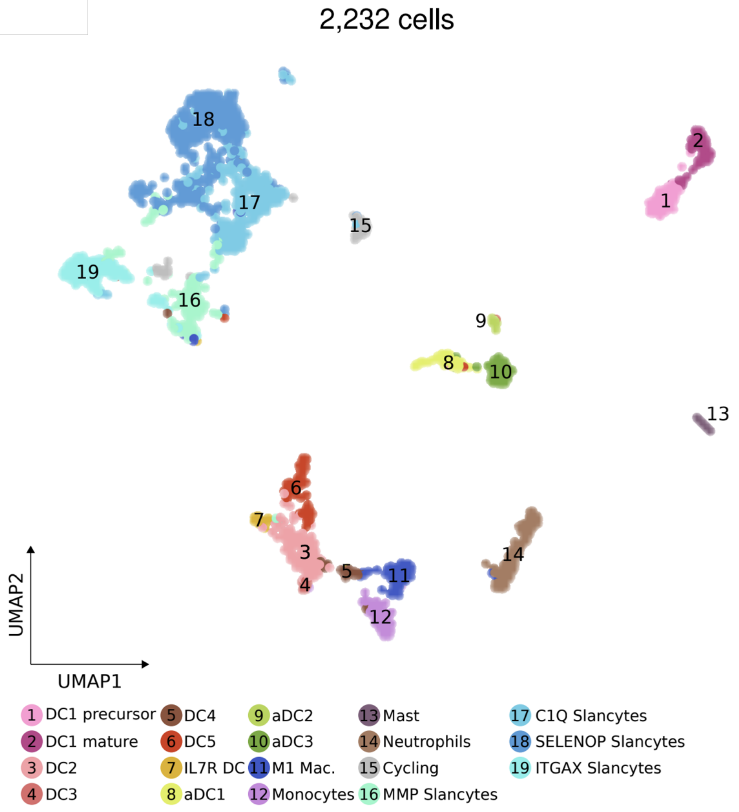


**DC1 precursor:** DC1 corresponds to the cross-presenting conventional DC. This cluster is best marked by *CLEC9A,* and the top gene markers are: *CST3, SNX3, C1orf54, HLA-DPB1, IRF8, CCND1* and *HLA-DPA1.* The low levels of XCR1 identify this population as a DC1 precursor (Balan et al., 2018).

**DC1 mature:** This cluster had a similar profile of DC1 precursor expressing higher levels of *CLEC9A* and higher levels of XCR1 acquiring a mature phenotype (Balan et al., 2018).

**DC2:** This cluster corresponds to "non-inflammatory" CD1C+ DC. CD1C was the main marker and unique that identified both DC2 and DC3 clusters. This subpopulation had an expression of *CLEC10A, FCER1A, HLA-DQB1, HLA-DRB1, HLA-DQA1 and HLA-DPB1.* This higher expression of MHC class II genes differentiates this population from DC3 (Villani et al., 2017).

**DC3:** This cluster corresponds to "inflammatory" CD1C+ DC. The higher expression CD1C identified these populations as the second cluster of CD1C DC cells. This cluster expressed *S100A8, S100A9, VCAN, LYZ* and *ANXA1.* This inflammatory signature classified this subpopulation as a different one compared with DC2 that had more antigen presentation features (Villani et al., 2017).

**DC4:** DC4 corresponds to CD1C–CD141–CD11C+ DC, which is best marked by CD16 and shares signatures with monocytes. This cluster expressed *FCGR3A, FTL, SERPINA1, LST1* and *AIF1* and it was CD1C- (Villani et al., 2017).

**DC5:** This cluster represents "AS DCs" AXL SIGLEC DCs this subpopulation has a similar profile to monocytes but displays a pDC signature. This subpopulation had as a top marker genes *SIGLEC6, AXL, CDH1, CD22* and *LILRA4* (Villani et al., 2017).

**IL7R DC:** This cluster was identified for its higher expression of *IL7R* and the expression of: *NR4A3, CLDN1, NRARP, CLEC10A* and *AXL*. This cluster was described as a pDC or DC5 precursor based on the gene expression profile (Rodrigues et al., 2018).

**aDC1, aDC2 and aDC3:** These three subtypes of activated DC cells were recently described by Park et al. These three subtypes were characterized by the expression of *LAMP3* and *CCR7*. aDC1 is a cluster that shares the profile of DC1 cells with an activated profile. This cluster has top genes: *SLCO5A1, CCR7, DUSP5, LAD1, TREML1* and *LAMP3*. aDC2 is derived from DC2 cells with a specific chemokine pattern compared with DC1. aDC2 expresses *PRDM16, CCNA1, SLC16A2, S100A2, AIRE* and *CCL22.* aDC3 induces the recruitment via CCR7. This cluster expressed *CCL19, MARCKSL1, FSCN1, ACTG1, SYNPO* and *TMSB10* (Park et al., 2020).

**M1 Mac:** This population had a higher expression of *CD68* and the top gene markers are an inflammatory signature of M1 profile. This cluster has top gene markers: *CCL4, IL1B, CCL3, TNF, IL6, IL1A* and *PTX3*.

**Monocytes:** This cluster was identified by its expression of *CD14* and the expression of canonical genes well described for monocytes. Top markers of this cluster: *VCAN, S100A9, S100A8, FCN1, S100A4* and *S100A12* (Villani et al., 2017).

**Mast cells:** This cluster has expressions of classical markers for Mast cells *KIT*, *TPSAB1* and *TPSB2*. Top markers of this cluster were: *TPSAB1, TPSB2, CPA3, MS4A2, LTC4S, HPGD, HPGDS* and *LMO4* (Wu et al., 2022a).

**Granulocytes (Neutrophils):** This cluster had expression of specific markers previously described for granulocytes. Top gene markers: *CXCL8, G0S2, PI3, CCL3L1, FCGR3B, PTGS2,* and *IFITM2* (Travaglini et al., 2020; Williams et al., 2021).

**Cycling:** This cluster is characterized by high proliferation signature expressing *CCNB2, PCNA, MKi67* and *TOP2A.*

**MMP Slancytes:** This cluster had a profile of tonsil slan+ cells expressing higher levels of *MMP9, MMP12, MMP14, HLA-DRA, HLA-DRB, CCR1, FCGR2A, APOE, ADAMDEC1* and *C1QC,* and intermediate levels of *NECTIN2* and *FUCA1* (Bianchetto-Aguilera et al., 2020). Based on GO terms, these MMP slancytes could play a role in antigen presentation, activation and matrix remodeling (Hofer et al., 2015).

**C1Q Slancytes**: This population had higher levels of tonsil slan+ cells such as *C1QC, APOE,* and *MMP9,* and intermediate levels of *FUCA1, APOC1* and *ADAMDEC1*. This cluster had a profile to respond to classical complement activation with higher expression of C1Q family genes: *C1QC, C1QB* and *C1QA*. Also, This cluster had a higher expression of MHC class II genes *HLA-DQA1, HLA-DRA, HLA-DRB1, HLA-DPA1* and *HLA-DPB1*, demonstrating their role in antigen presentation (Bianchetto-Aguilera et al., 2020) .

**SELENOP Slancytes:** These SELENOP slancytes expressed a tonsil slan+ cell signature with higher levels of *FUCA1, APOE* and *APOC1*, and lower levels of *C1QC, MMP9* and *ADAMDEC1*. This cluster expressed higher levels of *CCL18* suggesting it can be involved in the recruitment of MMP slancytes via CCL18-CCR1 axis. This cluster had an OXPHOS related signature pointing out their possible metabolic role in tonsils (Bianchetto-Aguilera et al., 2020).

**ITGAX Slancytes:** This cluster expressed tonsil slan+ cell markers: *FUCA1, NR1H3*, and *APOE.* This cluster has an inflammatory signature expressing *ITGAX, ZEB2, IL6R* and *IL18.*

### EPITHELIAL CELLS

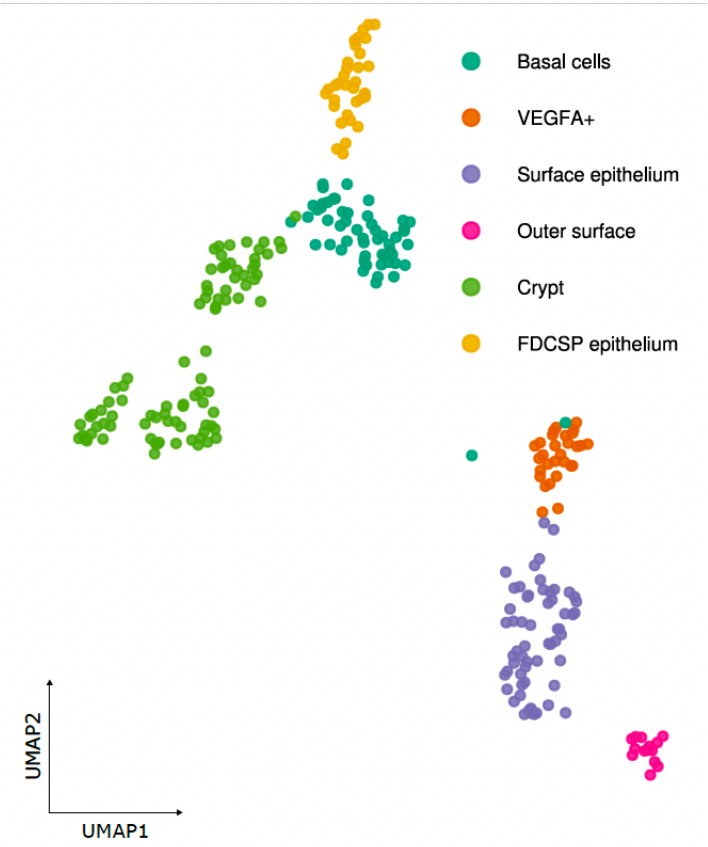


**Basal cells:** This cluster represents the basal layer of the squamous epithelium, characterized by the expression of the basal cytokeratins *KRT5* and *KRT14* (Williams et al., 2021).

**Surface keratinocytes:** Cells in this cluster represent the main bulk of the surface epithelium with expression of several cytokeratins (*KRT4, KRT13, KRT78, KRT80*), *SPRR3, TMPRSS11B* and *TMPRSS2* (Williams et al., 2021) .

**VEGFA+**: Cells in this cluster show an expression of *VEGFA*, which has been shown to be expressed by subsets of epithelial cells in other organs.

**Outer surface:** These cells express several genes associated with cornification/keratinization like *CNFN* and *SPRR2* genes and represent the outermost layer of the surface epithelium.

**FDCSP epithelium:** This cluster was named after the high expression of *FDCSP*, which was first described to be expressed in both FDCs and part of the crypt epithelium of tonsils.

**Crypt:** Cells in this cluster express *KRT8* and genes involved in inflammatory responses (IL1B, IFI27) and represent the epithelial crypts of the tonsils. Also in these clusters are cells which express *SPIB* and *MARCKSL1*, two genes associated with microfold cells (M cells).

### MESENCHYMAL CELLS/FOLLICULAR DENDRITIC CELLS

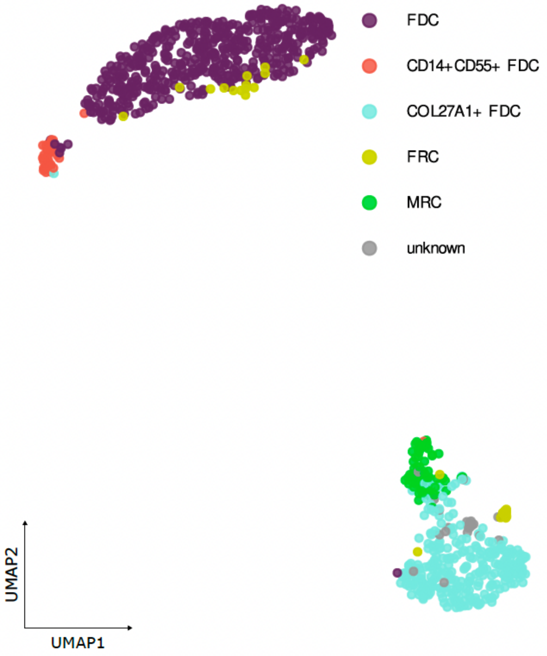


**FDC**: This is the main cluster of follicular dendritic cells, with high expression of the classical FDC markers *FDCSP*, *CLU* and *CXCL13* (Heesters et al., 2021).

**CD14+ CD55+ FDC**: This cluster, besides high expression levels of *FDCSP*, *CLU* and *CXCL13*, additionally expresses CD14 and CD55. CD14 is described in the literature as being expressed by a subset of follicular dendritic cells (Smeltzer et al., 2014).

**COL27A1 FDC**: Cells in this cluster show lower expression levels of FDCSP, CLU and CXCL13 compared to the main FDC cluster and high expression of COL27A1 (Rodda et al., 2018).

**FRC**: This cluster of cells shows expression of *CCL20*, *PERP*, *S100A2* and *S100A9* markers associated with fibroblastic reticular cells (Rodda et al., 2018).

**MRC**: Cells in this cluster express several collagen genes, as well as *DCN* and *PDGFRB*. This cluster was annotated as marginal reticular cells according to recently published literature. Based on *PDGFRB* expression at least part of these cells might represent perivascular precursors, as described in the literature in murine models (Jiang et al., 2021) .

**Unknown**: Cells in this cluster had as marker several genes from the KLF family (KLF2/4/6) as well as other markers such as EGR1, DUSP1, ZFP36, NFKBIZ, SELE, VMP1, ADAMTSL4-AS1, AC020916.1, AC007952.4. However, this cluster also showed as markers distinct stress signature responses such as FOS, JUN or NEAT1 and hence we did not know whether to exclude this cluster as it may represent a low-quality cluster.

**Cycling FDC**: This a cluster of follicular dendritic cells, with high expression of the classical FDC markers and a specific gene signature of cycling. The top expressed genes are: *HIST1H4C, HMGB2, STMN1, H2AFZ, HMGB1, HMGN2, NUSAP1, TUBA1B* and *PTTG1*. As we observed that the proliferative signature was a major driver of variance in scRNA-seq data, which caused (1) cells from different lineages to cluster together and (2) hindendered biological heterogeneity in the other cluster. Thus we show this population in Figure 1B, but exclude it from downstream analysis.

### *GLOSSARY REFERENCES*


Bal, S.M., Golebski, K., and Spits, H. (2020). Plasticity of innate lymphoid cell subsets. Nat. Rev. Immunol. *20*, 552–565. https://doi.org/10.1038/s41577-020-0282-9.

Balan, S., Arnold-Schrauf, C., Abbas, A., Couespel, N., Savoret, J., Imperatore, F., Villani, A.-C., Vu Manh, T.-P., Bhardwaj, N., and Dalod, M. (2018). Large-Scale Human Dendritic Cell Differentiation Revealing Notch-Dependent Lineage Bifurcation and Heterogeneity. Cell Rep. *24*, 1902-1915.e6. https://doi.org/10.1016/j.celrep.2018.07.033.

Bianchetto-Aguilera, F., Tamassia, N., Gasperini, S., Calzetti, F., Finotti, G., Gardiman, E., Montioli, R., Bresciani, D., Vermi, W., and Cassatella, M.A. (2020). Deciphering the fate of slan $^{+}$ -monocytes in human tonsils by gene expression profiling. FASEB J. *34*, 9269–9284. https://doi.org/10.1096/fj.202000181R.

Boldt, A., Borte, S., Fricke, S., Kentouche, K., Emmrich, F., Borte, M., Kahlenberg, F., and Sack, U. (2014). Eight-color immunophenotyping of T-, B-, and NK-cell subpopulations for characterization of chronic immunodeficiencies. Cytometry B Clin. Cytom. *86*, 191–206. https://doi.org/10.1002/cyto.b.21162.

Brewitz, A., Eickhoff, S., Dähling, S., Quast, T., Bedoui, S., Kroczek, R.A., Kurts, C., Garbi, N., Barchet, W., Iannacone, M., et al. (2017). CD8 + T Cells Orchestrate pDC-XCR1 + Dendritic Cell Spatial and Functional Cooperativity to Optimize Priming. Immunity *46*, 205–219. https://doi.org/10.1016/j.immuni.2017.01.003.

Cano-Gamez, E., Soskic, B., Roumeliotis, T.I., So, E., Smyth, D.J., Baldrighi, M., Willé, D., Nakic, N., Esparza-Gordillo, J., Larminie, C.G.C., et al. (2020). Single-cell transcriptomics identifies an effectorness gradient shaping the response of CD4+ T cells to cytokines. Nat. Commun. *11*, 1801. https://doi.org/10.1038/s41467-020-15543-y.

Cao, W., and Bover, L. (2010). Signaling and ligand interaction of ILT7: receptor-mediated regulatory mechanisms for plasmacytoid dendritic cells. Immunol. Rev. *234*, 163–176. https://doi.org/10.1111/j.0105-2896.2009.00867.x.

Choi, J., and Crotty, S. (2021). Bcl6-Mediated Transcriptional Regulation of Follicular Helper T cells (TFH). Trends Immunol. *42*, 336–349. https://doi.org/10.1016/j.it.2021.02.002.

Cibrián, D., and Sánchez-Madrid, F. (2017). CD69: from activation marker to metabolic gatekeeper. Eur. J. Immunol. *47*, 946–953. https://doi.org/10.1002/eji.201646837.

Crotty, S. (2011). Follicular Helper CD4 T Cells ($T_{FH}$). Annu. Rev. Immunol. *29*, 621–663. https://doi.org/10.1146/annurev-immunol-031210-101400.

Crotty, S. (2014). T Follicular Helper Cell Differentiation, Function, and Roles in Disease. Immunity *41*, 529–542. https://doi.org/10.1016/j.immuni.2014.10.004.

Crotty, S. (2019). T Follicular Helper Cell Biology: A Decade of Discovery and Diseases. Immunity *50*, 1132–1148. https://doi.org/10.1016/j.immuni.2019.04.011.

Cuenca, M., Romero, X., Sintes, J., Terhorst, C., and Engel, P. (2016). Targeting of Ly9 (CD229) Disrupts Marginal Zone and B1 B Cell Homeostasis and Antibody Responses. J. Immunol. Baltim. Md 1950 *196*, 726–737. https://doi.org/10.4049/jimmunol.1501266.

Damsker, J.M., Hansen, A.M., and Caspi, R.R. (2010). Th1 and Th17 cells: Adversaries and collaborators. Ann. N. Y. Acad. Sci. *1183*, 211–221. https://doi.org/10.1111/j.1749-6632.2009.05133.x.

Dominguez-Sola, D., Victora, G.D., Ying, C.Y., Phan, R.T., Saito, M., Nussenzweig, M.C., and Dalla-Favera, R. (2012). The proto-oncogene MYC is required for selection in the germinal center and cyclic reentry. Nat. Immunol. *13*, 1083–1091. https://doi.org/10.1038/ni.2428.

Du, Y., Fang, Q., and Zheng, S.-G. (2021). Regulatory T Cells: Concept, Classification, Phenotype, and Biological Characteristics. In T Regulatory Cells in Human Health and Diseases, S.-G. Zheng, ed. (Singapore: Springer Singapore), pp. 1–31.

Ferrer, G., Bosch, R., Hodgson, K., Tejero, R., Roué, G., Colomer, D., Montserrat, E., and Moreno, C. (2014). B cell activation through CD40 and IL4R ligation modulates the response of chronic lymphocytic leukaemia cells to BAFF and APRIL. Br. J. Haematol. *164*, 570–578. https://doi.org/10.1111/bjh.12645.

Galletti, G., De Simone, G., Mazza, E.M.C., Puccio, S., Mezzanotte, C., Bi, T.M., Davydov, A.N., Metsger, M., Scamardella, E., Alvisi, G., et al. (2020). Two subsets of stem-like CD8+ memory T cell progenitors with distinct fate commitments in humans. Nat. Immunol. *21*, 1552–1562. https://doi.org/10.1038/s41590-020-0791-5.

Galván-Peña, S., Leon, J., Chowdhary, K., Michelson, D.A., Vijaykumar, B., Yang, L., Magnuson, A.M., Chen, F., Manickas-Hill, Z., Piechocka-Trocha, A., et al. (2021). Profound Treg perturbations correlate with COVID-19 severity. Proc. Natl. Acad. Sci. *118*, e2111315118. https://doi.org/10.1073/pnas.2111315118.

Heesters, B.A., van Megesen, K., Tomris, I., de Vries, R.P., Magri, G., and Spits, H. (2021). Characterization of human FDCs reveals regulation of T cells and antigen presentation to B cells. J. Exp. Med. *218*, e20210790. https://doi.org/10.1084/jem.20210790.

Hofer, T.P., Zawada, A.M., Frankenberger, M., Skokann, K., Satzl, A.A., Gesierich, W., Schuberth, M., Levin, J., Danek, A., Rotter, B., et al. (2015). slan-defined subsets of CD16-positive monocytes: impact of granulomatous inflammation and M-CSF receptor mutation. Blood *126*, 2601–2610. https://doi.org/10.1182/blood-2015-06-651331.

Holmes, A.B., Corinaldesi, C., Shen, Q., Kumar, R., Compagno, N., Wang, Z., Nitzan, M., Grunstein, E., Pasqualucci, L., Dalla-Favera, R., et al. (2020). Single-cell analysis of germinal-center B cells informs on lymphoma cell of origin and outcome. J. Exp. Med. *217*. https://doi.org/10.1084/JEM.20200483.

Ise, W., Kohyama, M., Schraml, B.U., Zhang, T., Schwer, B., Basu, U., Alt, F.W., Tang, J., Oltz, E.M., Murphy, T.L., et al. (2011). The transcription factor BATF controls the global regulators of class-switch recombination in both B cells and T cells. Nat. Immunol. *12*, 536–543. https://doi.org/10.1038/ni.2037.

Jacob, J., and Kelsoe, G. (1992). In situ studies of the primary immune response to (4-hydroxy-3-nitrophenyl)acetyl. II. A common clonal origin for periarteriolar lymphoid sheath-associated foci and germinal centers. J. Exp. Med. *176*, 679–687. https://doi.org/10.1084/jem.176.3.679.

Jia, L., and Wu, C. (2014). The Biology and Functions of Th22 Cells. In T Helper Cell Differentiation and Their Function, B. Sun, ed. (Dordrecht: Springer Netherlands), pp. 209–230.

Jiang, L., Yilmaz, M., Uehara, M., Cavazzoni, C.B., Kasinath, V., Zhao, J., Naini, S.M., Li, X., Banouni, N., Fiorina, P., et al. (2021). Characterization of Leptin Receptor+ Stromal Cells in Lymph Node. Front. Immunol. *12*, 730438. https://doi.org/10.3389/fimmu.2021.730438.

Khodadadi, L., Cheng, Q., Radbruch, A., and Hiepe, F. (2019). The Maintenance of Memory Plasma Cells. Front. Immunol. *10*, 721. https://doi.org/10.3389/fimmu.2019.00721.

King, H.W., Orban, N., Riches, J.C., Clear, A.J., Warnes, G., Teichmann, S.A., and James, L.K. (2021). Single-cell analysis of human B cell maturation predicts how antibody class switching shapes selection dynamics. Sci. Immunol. *6*. https://doi.org/10.1126/sciimmunol.abe6291.

Kok, L., Masopust, D., and Schumacher, T.N. (2022). The precursors of CD8+ tissue resident memory T cells: from lymphoid organs to infected tissues. Nat. Rev. Immunol. *22*, 283–293. https://doi.org/10.1038/s41577-021-00590-3.

Kräutler, N.J., Suan, D., Butt, D., Bourne, K., Hermes, J.R., Chan, T.D., Sundling, C., Kaplan, W., Schofield, P., Jackson, J., et al. (2017). Differentiation of germinal center B cells into plasma cells is initiated by high-affinity antigen and completed by Tfh cells. J. Exp. Med. *214*, 1259–1267. https://doi.org/10.1084/jem.20161533.

Krueger, P.D., Goldberg, M.F., Hong, S.-W., Osum, K.C., Langlois, R.A., Kotov, D.I., Dileepan, T., and Jenkins, M.K. (2021). Two sequential activation modules control the differentiation of protective T helper-1 (Th1) cells. Immunity *54*, 687-701.e4. https://doi.org/10.1016/j.immuni.2021.03.006.

Li, H., Dement-Brown, J., Liao, P.-J., Mazo, I., Mills, F., Kraus, Z., Fitzsimmons, S., and Tolnay, M. (2020). Fc receptor-like 4 and 5 define human atypical memory B cells. Int. Immunol. *32*, 755–770. https://doi.org/10.1093/intimm/dxaa053.

Ma, C.S., Wong, N., Rao, G., Nguyen, A., Avery, D.T., Payne, K., Torpy, J., O'Young, P., Deenick, E., Bustamante, J., et al. (2016). Unique and shared signaling pathways cooperate to regulate the differentiation of human CD4+ T cells into distinct effector subsets. J. Exp. Med. *213*, 1589–1608. https://doi.org/10.1084/jem.20151467.

Mahnke, Y.D., Brodie, T.M., Sallusto, F., Roederer, M., and Lugli, E. (2013). The who's who of T-cell differentiation: Human memory T-cell subsets: HIGHLIGHTS. Eur. J. Immunol. *43*, 2797–2809. https://doi.org/10.1002/eji.201343751.

Mathewson, N.D., Ashenberg, O., Tirosh, I., Gritsch, S., Perez, E.M., Marx, S., Jerby-Arnon, L., Chanoch-Myers, R., Hara, T., Richman, A.R., et al. (2021). Inhibitory CD161 receptor identified in glioma-infiltrating T cells by single-cell analysis. Cell *184*, 1281-1298.e26. https://doi.org/10.1016/j.cell.2021.01.022.

Nicolet, B.P., Guislain, A., and Wolkers, M.C. (2021). CD29 Enriches for Cytotoxic Human CD4 $^{+}$ T Cells. J. Immunol. *207*, 2966–2975. https://doi.org/10.4049/jimmunol.2100138.

Nörenberg, J., Jaksó, P., and Barakonyi, A. (2021). Gamma/Delta T Cells in the Course of Healthy Human Pregnancy: Cytotoxic Potential and the Tendency of CD8 Expression Make CD56+ γδT Cells a Unique Lymphocyte Subset. Front. Immunol. *11*, 596489. https://doi.org/10.3389/fimmu.2020.596489.

Park, J.E., Botting, R.A., Conde, C.D., Popescu, D.M., Lavaert, M., Kunz, D.J., Goh, I., Stephenson, E., Ragazzini, R., Tuck, E., et al. (2020). A cell atlas of human thymic development defines T cell repertoire formation. Science *367*. https://doi.org/10.1126/science.aay3224.

Pellicci, D.G., Koay, H.-F., and Berzins, S.P. (2020). Thymic development of unconventional T cells: how NKT cells, MAIT cells and γδ T cells emerge. Nat. Rev. Immunol. *20*, 756–770. https://doi.org/10.1038/s41577-020-0345-y.

Perdomo-Celis, F., Taborda, N.A., and Rugeles, M.T. (2017). Follicular CD8+ T Cells: Origin, Function and Importance during HIV Infection. Front. Immunol. *8*, 1241. https://doi.org/10.3389/fimmu.2017.01241.

Pizzolato, G., Kaminski, H., Tosolini, M., Franchini, D.-M., Pont, F., Martins, F., Valle, C., Labourdette, D., Cadot, S., Quillet-Mary, A., et al. (2019). Single-cell RNA sequencing unveils the shared and the distinct cytotoxic hallmarks of human TCRVδ1 and TCRVδ2 γδ T lymphocytes. Proc. Natl. Acad. Sci. *116*, 11906–11915. https://doi.org/10.1073/pnas.1818488116.

Qi, H. (2016). T follicular helper cells in space-time. Nat. Rev. Immunol. *16*, 612–625. https://doi.org/10.1038/nri.2016.94.

Rodda, L.B., Lu, E., Bennett, M.L., Sokol, C.L., Wang, X., Luther, S.A., Barres, B.A., Luster, A.D., Ye, C.J., and Cyster, J.G. (2018). Single-Cell RNA Sequencing of Lymph Node Stromal Cells Reveals Niche-Associated Heterogeneity. Immunity *48*, 1014-1028.e6. https://doi.org/10.1016/j.immuni.2018.04.006.

Rodrigues, P.F., Alberti-Servera, L., Eremin, A., Grajales-Reyes, G.E., Ivanek, R., and Tussiwand, R. (2018). Distinct progenitor lineages contribute to the heterogeneity of plasmacytoid dendritic cells. Nat. Immunol. *19*, 711–722. https://doi.org/10.1038/s41590-018-0136-9.

Sallusto, F., Geginat, J., and Lanzavecchia, A. (2004). Central Memory and Effector Memory T Cell Subsets: Function, Generation, and Maintenance. Annu. Rev. Immunol. *22*, 745–763. https://doi.org/10.1146/annurev.immunol.22.012703.104702.

Seifert, M., Sellmann, L., Bloehdorn, J., Wein, F., Stilgenbauer, S., Dürig, J., and Küppers, R. (2012). Cellular origin and pathophysiology of chronic lymphocytic leukemia. J. Exp. Med. *209*, 2183–2198. https://doi.org/10.1084/jem.20120833.

Smeltzer, J.P., Jones, J.M., Ziesmer, S.C., Grote, D.M., Xiu, B., Ristow, K.M., Yang, Z.Z., Nowakowski, G.S., Feldman, A.L., Cerhan, J.R., et al. (2014). Pattern of CD14+ follicular dendritic cells and PD1+ T cells independently predicts time to transformation in follicular lymphoma. Clin. Cancer Res. Off. J. Am. Assoc. Cancer Res. *20*, 2862–2872. https://doi.org/10.1158/1078-0432.CCR-13-2367.

Stark, J.M., Tibbitt, C.A., and Coquet, J.M. (2019). The Metabolic Requirements of Th2 Cell Differentiation. Front. Immunol. *10*, 2318. https://doi.org/10.3389/fimmu.2019.02318.

Travaglini, K.J., Nabhan, A.N., Penland, L., Sinha, R., Gillich, A., Sit, R.V., Chang, S., Conley, S.D., Mori, Y., Seita, J., et al. (2020). A molecular cell atlas of the human lung from single-cell RNA sequencing. Nature *587*, 619–625. https://doi.org/10.1038/s41586-020-2922-4.

Villani, A.-C., Satija, R., Reynolds, G., Sarkizova, S., Shekhar, K., Fletcher, J., Griesbeck, M., Butler, A., Zheng, S., Lazo, S., et al. (2017). Single-cell RNA-seq reveals new types of human blood dendritic cells, monocytes, and progenitors. Science *356*, eaah4573. https://doi.org/10.1126/science.aah4573.

Williams, D.W., Greenwell-Wild, T., Brenchley, L., Dutzan, N., Overmiller, A., Sawaya, A.P., Webb, S., Martin, D., NIDCD/NIDCR Genomics and Computational Biology Core, Hajishengallis, G., et al. (2021). Human oral mucosa cell atlas reveals a stromal-neutrophil axis regulating tissue immunity. Cell *184*, 4090-4104.e15. https://doi.org/10.1016/j.cell.2021.05.013.

Wing, J.B., Kitagawa, Y., Locci, M., Hume, H., Tay, C., Morita, T., Kidani, Y., Matsuda, K., Inoue, T., Kurosaki, T., et al. (2017). A distinct subpopulation of CD25 ⁻ T-follicular regulatory cells localizes in the germinal centers. Proc. Natl. Acad. Sci. *114*. https://doi.org/10.1073/pnas.1705551114.

Wu, C., Boey, D., Bril, O., Grootens, J., Vijayabaskar, M.S., Sorini, C., Ekoff, M., Wilson, N.K., Ungerstedt, J.S., Nilsson, G.P., et al. (2022a). Single-cell transcriptomics reveals the identity and regulators of human mast cell progenitors. Blood Adv. bloodadvances.2022006969. https://doi.org/10.1182/bloodadvances.2022006969.

Wu, Z., Zheng, Y., Sheng, J., Han, Y., Yang, Y., Pan, H., and Yao, J. (2022b). CD3+CD4-CD8- (Double-Negative) T Cells in Inflammation, Immune Disorders and Cancer. Front. Immunol. *13*, 816005. https://doi.org/10.3389/fimmu.2022.816005.

Yoon, S.-O., Zhang, X., Lee, I.Y., Spencer, N., Vo, P., and Choi, Y.S. (2013). CD9 is a novel marker for plasma cell precursors in human germinal centers. Biochem. Biophys. Res. Commun. *431*, 41–46. https://doi.org/10.1016/j.bbrc.2012.12.102.

Zhu, X., and Zhu, J. (2020). CD4 T Helper Cell Subsets and Related Human Immunological Disorders. Int. J. Mol. Sci. *21*, 8011. https://doi.org/10.3390/ijms21218011.
