## Supplementary Figures for "An Atlas of Cells in the Human Tonsil"

**Supplementary Figure Legends (S1-S10)**

**Supplementary Figures (S1-S10)**

**Figure S1. A single-cell multiomic atlas of human tonsillar cells**. ***Related to Figure 1.*** (**A**,**F**) UMAP projection of unintegrated (*left*) and harmony-based integrated single-cell datasets (*right*) splitted and colored by data modality (scRNA-seq/scATAC-seq or Multiome) or external dataset (Hamish King *et al.)* (**B, G**) Boxplot comparing local inverse simpson index (LISI) across confounders of integrated and unintegrated scRNA-seq (B) and scATAC-seq (G) datasets. (**C**) UMAP projection of the King *et al.* dataset colored by the original publication. **(D)** UMAP projection of the 357,206 tonsillar cells analyzed colored by the main 9 populations identified: Naive and Memory B cells (NBC_MBC), Germinal center B cells (GCBC), plasma cells (PC), CD4 T cells, cytotoxic (CD8 T cells, NK, ILC, double negative T cells (DN), myeloid cells (DC, macrophages, monocytes, granulocytes, mast cells), follicular dendritic cells (FDC), epithelial cells and plasmacytoid dendritic cells (PDC). **(E)** UMAP projection of tonsillar cells colored by cell cycle phase; *left*: S Score and *right*: G2M Score **(H)** UMAP projection of the prediction score probability inferred for CITE-seq (*left*) and scATAC-seq (*right*) after the label and coordinate transfer.

**Figure S2. Tonsil Visium slides histology, clustering and cell type deconvolution**. ***Related to Figure 1.*** *(***A**) H&E slide (*left*) next to gene expression-based spot clustering annotation (*right*). (**B**) Spatial scatter pie plot showing each spot as a pie chart representing the predicted proportion of each cell type.

**Figure S3. CD4 T follicular and non-follicular cell fate decision in the human tonsil.** ***Related to Figure 2.*** (**A**) Dotplot showing the average expression of markers for Naive, CM pre-non-Tfh and CM Pre Tfh. Dot size reflects the percentage of cells in a cluster expressing each gene and the color the average expression level **(B)** Violinplot of BCL6 accessibility values considering the gene body and 2,000 bp upstream (*top*) and the predicted distal cis-regulatoy region using Cicero as a portential enhancer (*bottom*). **(C)** Dotplot showing the average expression of the top 20 markers that described the main CD4 T subpopulations identified. **(D)** Violinplot of PRDM1 TF activity (AUCell score) across Naive, CM pre-non-Tfh and CM Pre Tfh. **(E)** Violinplot of CD28 (*top*) and CD29 (*bottom*) protein expression for each of the CD4 T sub-populations identified. **(F)** UMAPs projection colored by the estimated Nebulosa density expression (STAR Methods) for key interleukin and chemokine receptors. **(G)** Violinplot of FOXP3 (*top*) and PRDM1 (*bottom*) TF activity (AUCell score) across Treg subtypes. **(H)** Violinplot showing the effector Treg (eTreg, *top*) and circulating Tfr (cTfr, *bottom*) gene expression signatures from Wing JB, *et al.* computed using UCell R package for Treg subtypes.

**Figure S4. Cell type deconvolution of CD4 T cell subpopulations.** ***Related to Figure 2*** (**A**) Cell type deconvolution carried out with SPOTlight. We deconvoluted the spots using CD4 T cell subpopulations annotations along with the general annotation of other cell types ensuring we captured the biological signal of all our cell types. (**B**) Topic profiles of the deconvolution showing the model learned unique gene signatures for each cell type.

**Figure S5. Landscape of CD8 and innate lymphoid cells in the human tonsil.** ***Related to Figure 3.*** (**A**) Dotplot showing the average expression of the top markers for CD8 T cells and innate lymphoid cells (ILC). Dot size reflects the percentage of cells in a cluster expressing each gene and the color the average expression level. **(B)** Violinplot showing the protein expression of seven representative markers for different subpopulations in CD8 T and ILC.

**Figure S6. B cell activation and GC dynamics.** ***Related to Figure 4.*** (**A**) Dotplot showing marker expression for NBC and MBC subpopulations. (**B**) Dotplot showing marker expression per GCBC subpopulations. Dot size reflects the percentage of cells in a cluster expressing each gene and the color the average expression level. (**C**) UMAP projection of GCBC cells colored by cell cycle phase. (**D**) UMAP projection of NFKB signature computed using *UCell* R package with seven genes (*NFKBIA*, *NFKBID*, *NFKB1*, *NFKB2*, *REL*, *RELA*, *RELB* for NBC and MBC. (**E**) UMAP projection of *BATF* expression in NBC and MBC.

**Figure S7. Plasma cell differentiation and cell identity regulation in human tonsils. *Related to Figure 5.*** (**A**) PC BCR analysis. Top: CITE-seq/BCR cells projected in scRNA-seq UMAP coordinates. Cluster labels and coordinates are transferred via KNN classification and regression respectively. Bottom: Barplot representation of PC clonality. Blue, clonal expansion when >= 3 cells had identical CDR3 sequence; Green, two cells share identical sequence; Orange, cells with disting CD3 sequence. (**B**) Proliferative cells (PB committed, Transitional PB, PB) differentiation analysis. Top: UMAP projection of proliferative cells colored by cell cycle phase. Bottom: Barplot representation of PC-phenotypic markers expression across S and G2M cells from PB cluster (most representative proliferative cluster). (**C**) UMAPs projection of the expression of Ig and MBC genes (BANK1, CELF2, TXNIP). (**D**) Violin Plot of the Endoplasmic Reticulum (ER) signature across PC scRNA-seq clusters. Signature was obtained using the *UCell* R package with 70 genes obtained from the DAVID KEGG pathway analysis: Protein processing in endoplasmic reticulum. (**E**) Cell type proportions deconvoluted using SPOTlight (see Methods). (**F**) Spatially defined trajectories on H&E stained images from BCLL-10-T (left) and BCLL-12-T (right) patients and heatmap showing smoothed expression changes for specific genes across defined trajectories. BCLL-10-T trajectory starts in a LZ zone and includes an interfollicular zone. (**G**) Correlation plot showing co-localization of cell types on the visium slides, plot for slide BCLL-10-T (top) and for all slides combined (bottom). (**H**) UMAP projection of the activity (AUCell score) of VDR and CREB3 TFs. (**I**) Barplot of SIX5 expression in different subpopulations from peripheral blood scRNA-seq data from Hao Yuhan *et al.*. (**J**) Boxplot of PC phenotypic markers, SIX5 and SIX5 predicted target in different populations from bone marrow scRNA-seq data from Hay Stuart B *et al.*. Link http://www.altanalyze.org/ICGS/HCA/Viewer.php

**Figure S8. Epithelial cells in the human tonsils. *Related to Figure 6.*** (**A**) UMAP projection of epithelial tonsillar cells colored by scRNA-seq clusters. (**B**) Dotplot showing the average expression of the top markers that described the main clusters identified in the epithelial compartment. Dot size reflects the percentage of cells in a cluster expressing each gene and the color the average expression level. **(C)** Cell type proportions of populations of interest within each spot on slide BCLL-2-T along with an H&E image of the tissue slice. Cell type deconvolution was carried out with SPOTlight. We deconvoluted the spots using epithelial cell subpopulation annotations along with the general annotation of other cell types ensuring we captured the biological signal of all our cell types. (**D**) UMAP projection of follicular dendritic cells (FDC) colored by scRNA-seq clusters. (**E**) Dotplot showing the average expression of the top markers that described the main clusters identified. **(F)** MAGIC-normalized gene expression of genes of interest on slide BCLL-10-T.

**Figure S9. Novel roles of slancyte subsets in human tonsils**. ***Related to Figure 6.*** (**A**) Dotplot showing the average expression of the top markers that described the main clusters identified in the DC (*left*) and aDC (*right*) compartment. Dot size reflects the percentage of cells in a cluster expressing each gene and the color the average expression level. **(B)** Violinplot showing seven signatures computed using UCell R package for aDC subsets. **(C)** Stacked barplot showing the proportion of cells *(y-axis*) detected in each donor (*x-axis*) for each subpopulation. **(D)** Dotplot showing the average expression of three markers (C1QA, MMP12, SELENOP) across all defined cell types and states in the tonsil atlas.

**Figure S10. Extended analysis of two mantle cell lymphoma patients. *Related to Figure 7.*** (**A**) UMAP projection of six genes encoded in chromosome Y for MCL1 (top) and MCL2 (bottom). **(B)** Distribution (left) and UMAP projection of a chrY expression signature derived from the six genes shown in (A). Vertical dashed line shows the cutoff used to classify cells in chrY+/-. **(C)** UMAP projection of the annotated clusters for MCL2. **(D)** UMAP projection of cells classified as chrY+/-, as defined in (B). **(E)** Dotplot showing the average expression of the top markers that described the main clusters identified in for MCL2. Dot size reflects the percentage of cells in a cluster expressing each gene and the color the average expression level. **(F)** Jitter plot of cell cycle scores (top: S.Score, bottom: G2M.Score) across all clusters annotated for MCL2. **(G)** Heatmap showing the pseudobulk gene expression profiles of top 100 upregulated genes in MCL for clusters in MCL2, NBC/MBC, GCBC, and PC. Rows are clustered with hierarchical clustering

**Figure S1**

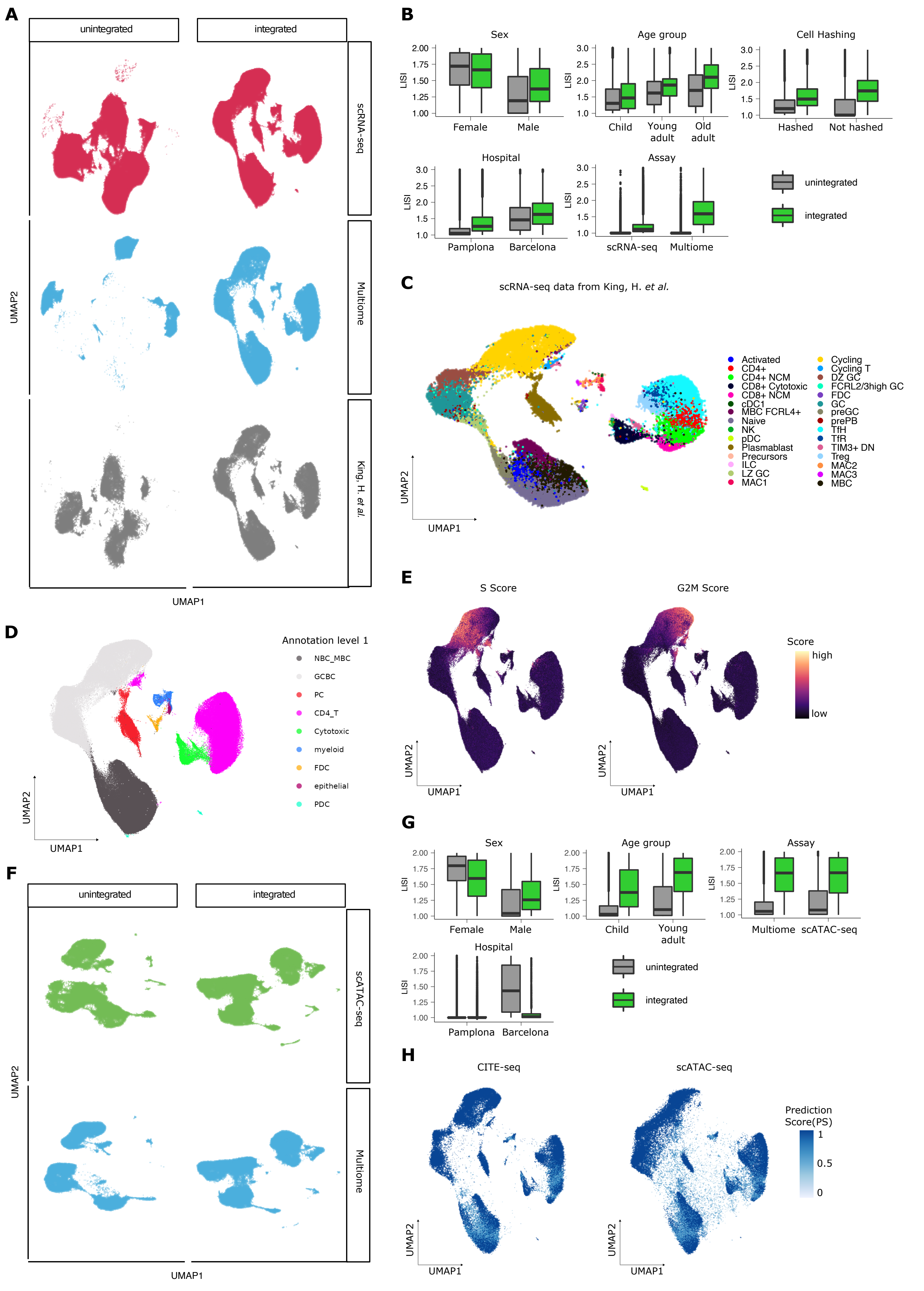
A
unintegrated
integrated
scRNA-seq
Multiome
King, H. et al.
UMAP2
UMAP1
B
Sex
LISI
Female
Male
Age group
Child
Young adult
Old adult
Cell Hashing
Hashed
Not hashed
Hospital
Pamplona
Barcelona
Assay
scRNA-seq
Multiome
unintegrated
integrated
C
scRNA-seq data from King, H. et al.
Activated
CD4+
CD4+ NCM
CD8+ Cytotoxic
CD8+ NCM
cDC1
MBC FCRL4+
Naive
NK
pDC
Plasmablast
Precursors
ILC
LZ GC
MAC1
Cycling
Cycling T
DZ GC
FCRL2/3high GC
FDC
GC
preGC
prePB
TfH
TfR
TIM3+ DN
Treg
MAC2
MAC3
MBC
UMAP2
UMAP1
D
Annotation level 1
NBC_MBC
GCBC
PC
CD4_T
Cytotoxic
myeloid
FDC
epithelial
PDC
UMAP2
UMAP1
E
S Score
G2M Score
Score
high
low
UMAP2
UMAP1
F
unintegrated
integrated
scATAC-seq
Multiome
UMAP2
UMAP1
G
Sex
LISI
Female
Male
Age group
Child
Young adult
Assay
Multiome
scATAC-seq
Hospital
Pamplona
Barcelona
unintegrated
integrated
H
CITE-seq
scATAC-seq
Prediction Score(PS)
1
0.5
0
UMAP2
UMAP1

### Figure S2

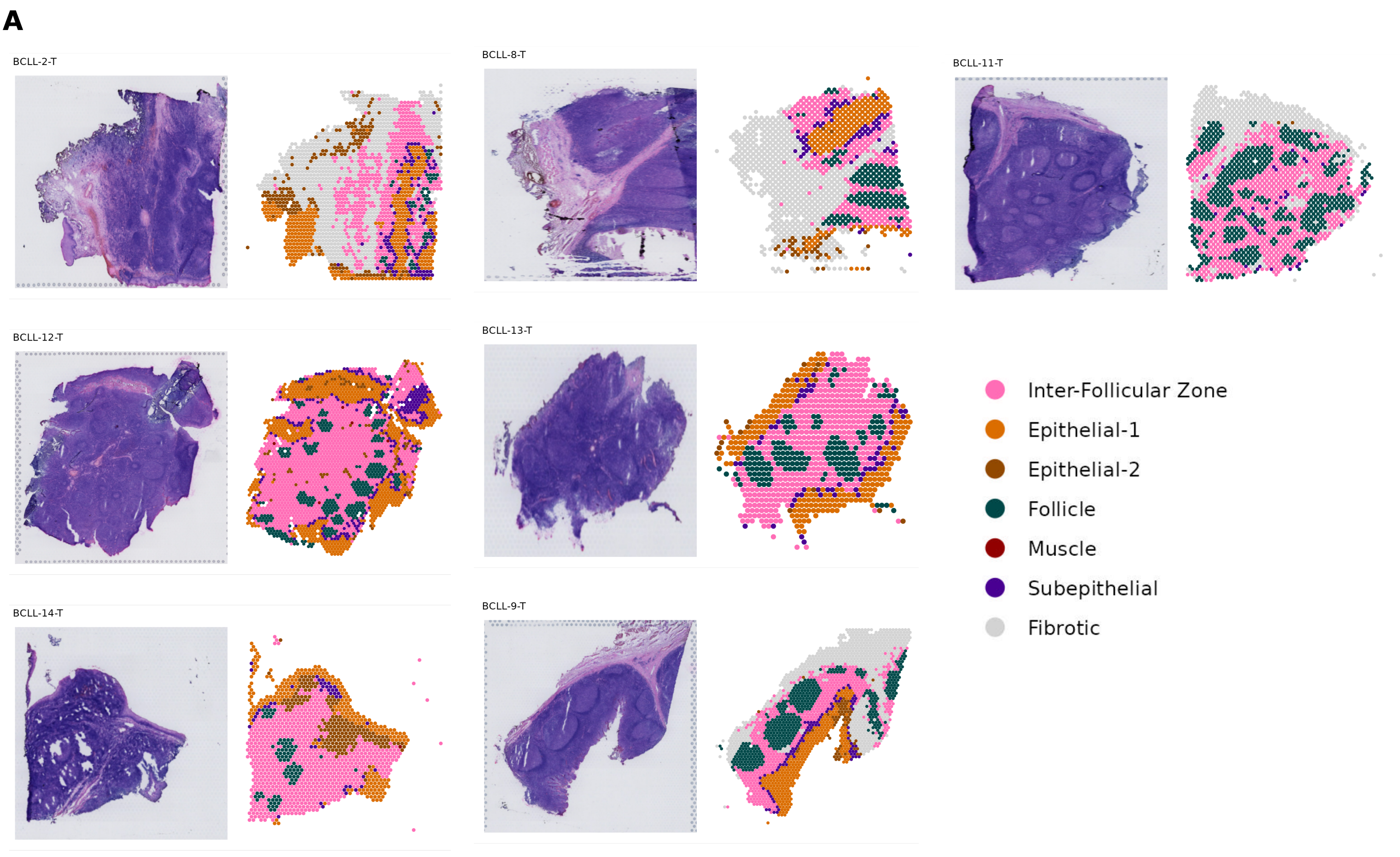
A
BCLL-2-T
BCLL-8-T
BCLL-11-T
BCLL-12-T
BCLL-13-T
BCLL-14-T
BCLL-9-T
Inter-Follicular Zone
Epithelial-1
Epithelial-2
Follicle
Muscle
Subepithelial
Fibrotic

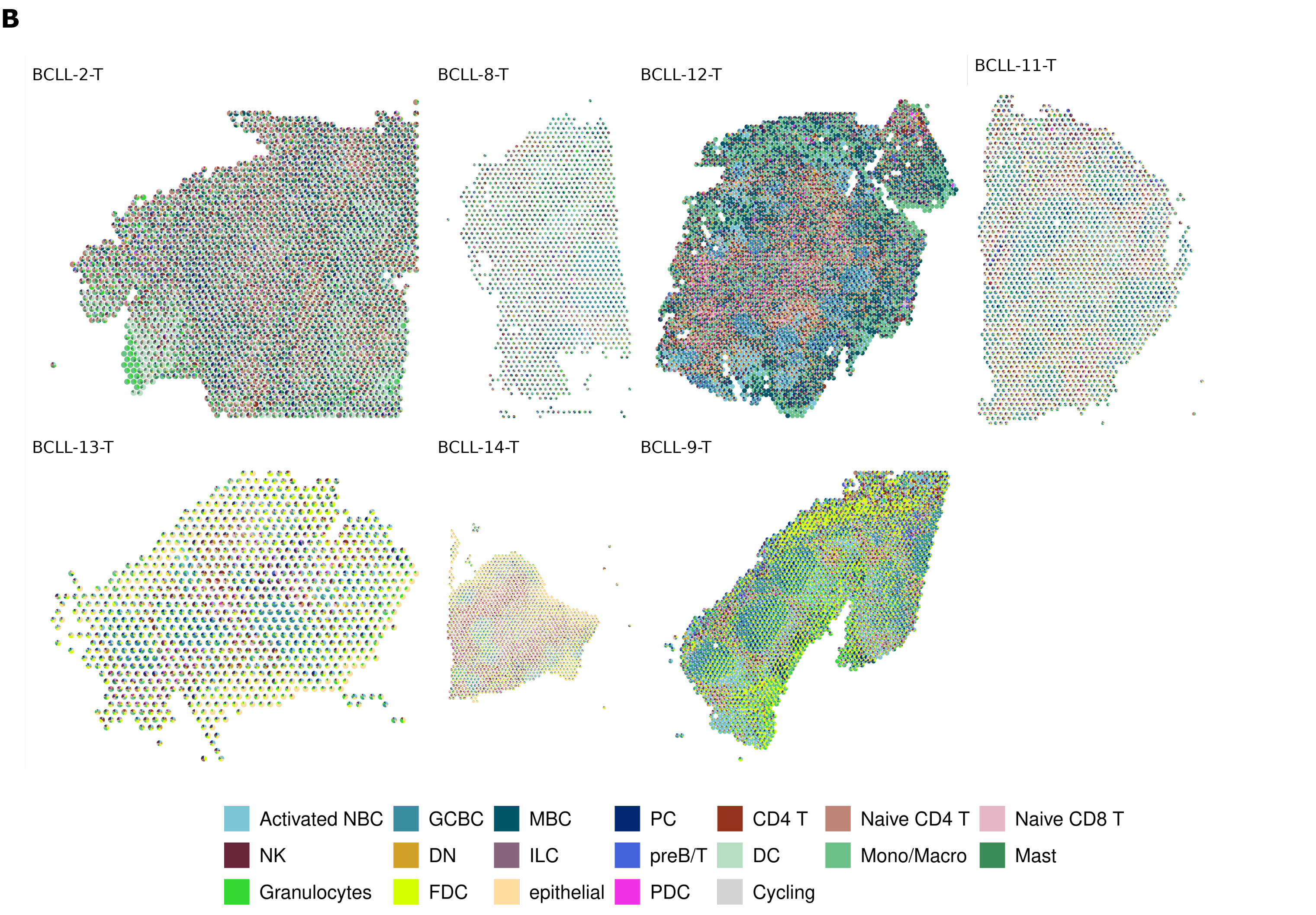
B
BCLL-2-T
BCLL-8-T
BCLL-12-T
BCLL-11-T
BCLL-13-T
BCLL-14-T
BCLL-9-T
Activated NBC
GCBC
MBC
PC
CD4 T
Naive CD4 T
Naive CD8 T
NK
DN
ILC
preB/T
DC
Mono/Macro
Mast
Granulocytes
FDC
epithelial
PDC
Cycling

**Figure S3**

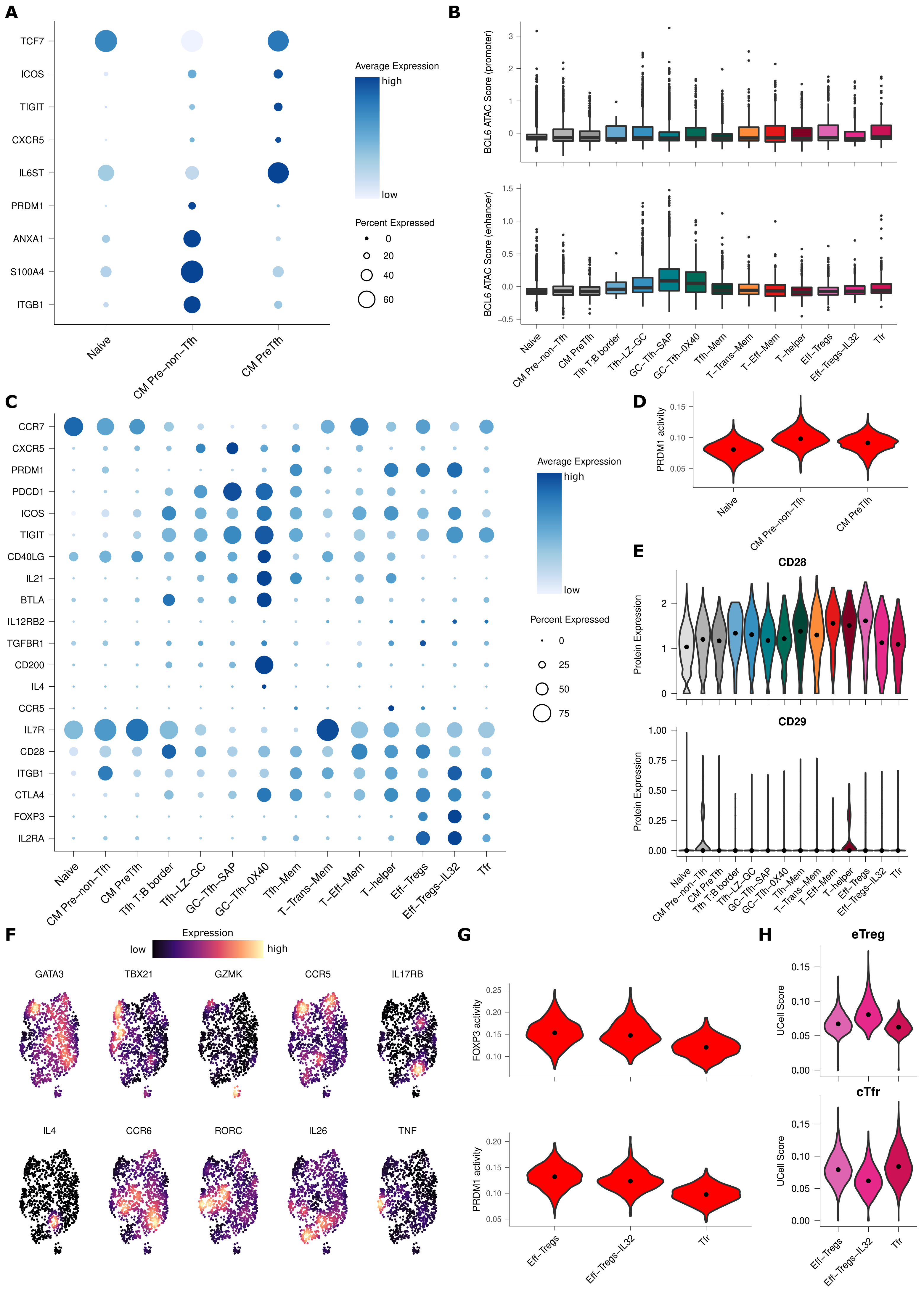

A
TCF7
ICOS
TIGIT
CXCR5
IL6ST
PRDM1
ANXA1
S100A4
ITGB1
Naive
CM Pre–non–Tfh
CM PreTfh
Average Expression
high
low
Percent Expressed
0
20
40
60
B
BCL6 ATAC Score (promoter)
BCL6 ATAC Score (enhancer)
Naive
CM Pre–non–Tfh
CM PreTfh
Tfh T:B border
Tfh–LZ–GC
GC–Tfh–SAP
GC–Tfh–0X40
Tfh–Mem
T–Trans–Mem
T–Eff–Mem
T–helper
Eff–Tregs
Eff–Tregs–IL32
Tfr
C
CCR7
CXCR5
PRDM1
PDCD1
ICOS
TIGIT
CD40LG
IL21
BTLA
IL12RB2
TGFBR1
CD200
IL4
CCR5
IL7R
CD28
ITGB1
CTLA4
FOXP3
IL2RA
Average Expression
high
low
Percent Expressed
0
25
50
75
D
PRDM1 activity
E
CD28
CD29
Protein Expression
F
Expression
low
high
GATA3
TBX21
GZMK
CCR5
IL17RB
IL4
CCR6
RORC
IL26
TNF
G
FOXP3 activity
PRDM1 activity
Eff–Tregs
Eff–Tregs–IL32
Tfr
H
eTreg
cTfr
UCell Score

### Figure S4

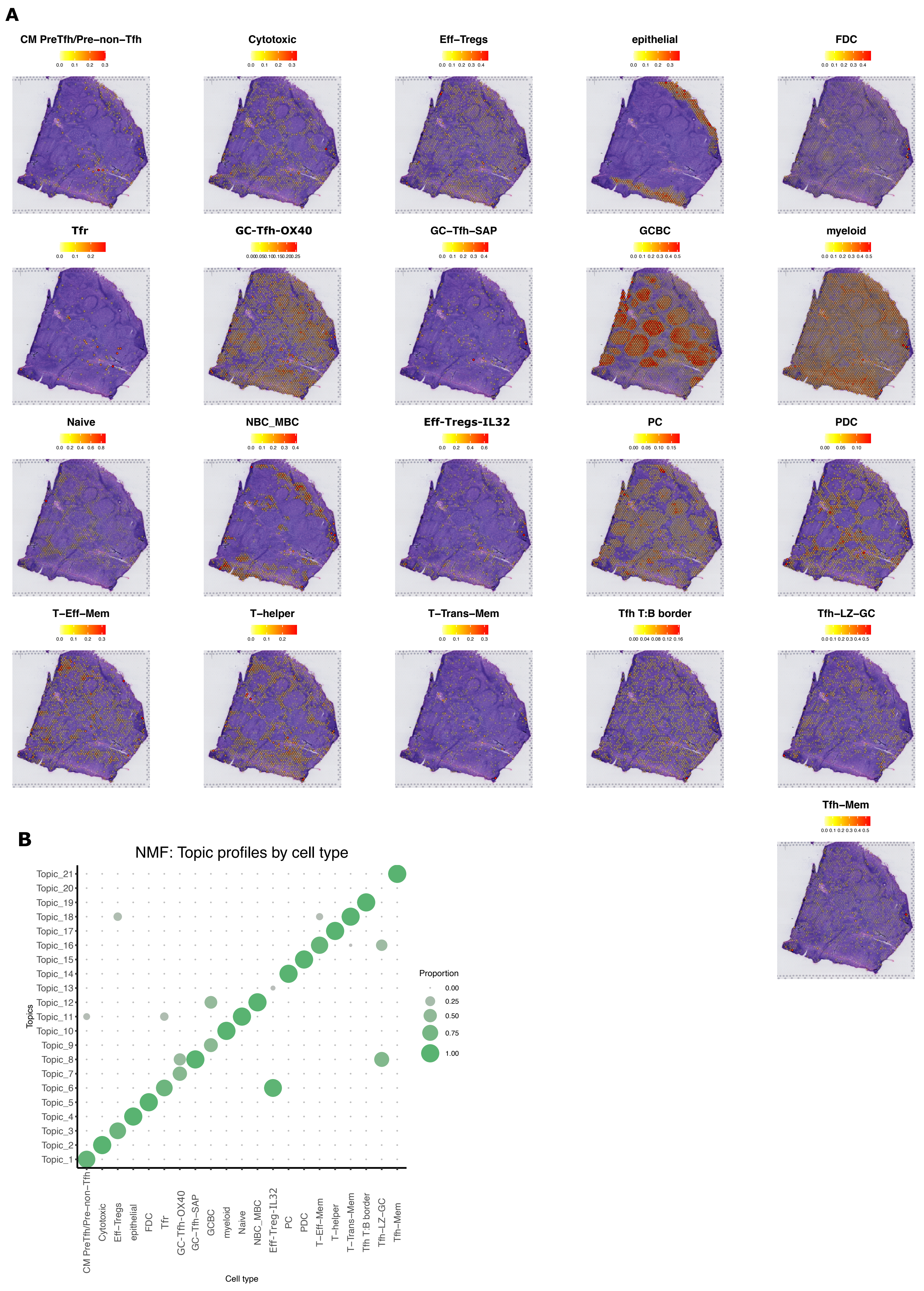
A
CM PreTfh/Pre–non–Tfh
Cytotoxic
Eff–Tregs
epithelial
FDC
Tfr
GC-Tfh-OX40
GC–Tfh–SAP
GCBC
myeloid
Naive
NBC_MBC
Eff-Tregs-IL32
PC
PDC
T–Eff–Mem
T–helper
T–Trans–Mem
Tfh T:B border
Tfh–LZ–GC
Tfh–Mem
B
NMF: Topic profiles by cell type
Topics
Topic_21
Topic_20
Topic_19
Topic_18
Topic_17
Topic_16
Topic_15
Topic_14
Topic_13
Topic_12
Topic_11
Topic_10
Topic_9
Topic_8
Topic_7
Topic_6
Topic_5
Topic_4
Topic_3
Topic_2
Topic_1
Proportion
0.00
0.25
0.50
0.75
1.00
CM PreTfh/Pre–non–Tfh
Cytotoxic
Eff–Tregs
epithelial
FDC
Tfr
GC–Tfh–OX40
GC–Tfh–SAP
GCBC
myeloid
Naive
NBC_MBC
Eff–Treg–IL32
PC
PDC
T–Eff–Mem
T–helper
T–Trans–Mem
Tfh T:B border
Tfh–LZ–GC
Tfh–Mem
Cell type

**Figure S5**

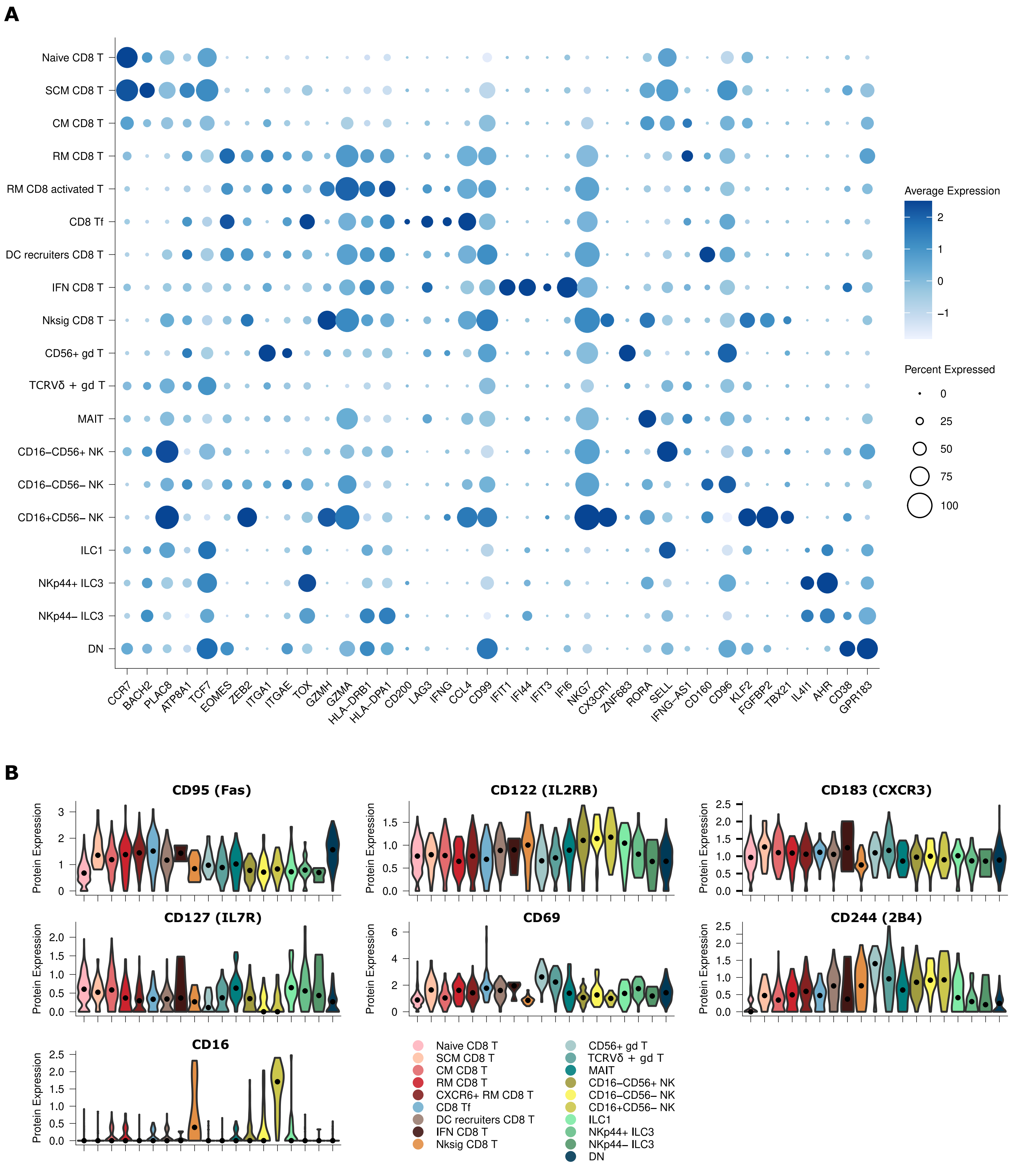
A
Naive CD8 T
SCM CD8 T
CM CD8 T
RM CD8 T
RM CD8 activated T
CD8 Tf
DC recruiters CD8 T
IFN CD8 T
Nksig CD8 T
CD56+ gd T
TCRVδ + gd T
MAIT
CD16−CD56+ NK
CD16−CD56− NK
CD16+CD56− NK
ILC1
NKp44+ ILC3
NKp44− ILC3
DN
CCR7 BACH2 PLAC8 ATP8A1 TCF7 EOMES ZEB2 ITGA1 ITGAE TOX GZMH GZMA HLA−DRB1 HLA−DPA1 CD200 LAG3 IFNG CCL4 CD99 IFIT1 IFI44 IFIT3 IFI6 NKG7 CX3CR1 ZNF683 RORA SELL IFNG−AS1 CD160 CD96 KLF2 FGFBP2 TBX21 IL4I1 AHR CD38 GPR183
Average Expression
2
1
0
−1
Percent Expressed
0
25
50
75
100
B
CD95 (Fas)
CD122 (IL2RB)
CD183 (CXCR3)
CD127 (IL7R)
CD69
CD244 (2B4)
CD16
Protein Expression
Naive CD8 T
SCM CD8 T
CM CD8 T
RM CD8 T
CXCR6+ RM CD8 T
CD8 Tf
DC recruiters CD8 T
IFN CD8 T
Nksig CD8 T
CD56+ gd T
TCRVδ + gd T
MAIT
CD16−CD56+ NK
CD16−CD56− NK
CD16+CD56− NK
ILC1
NKp44+ ILC3
NKp44− ILC3
DN

### Figure S6

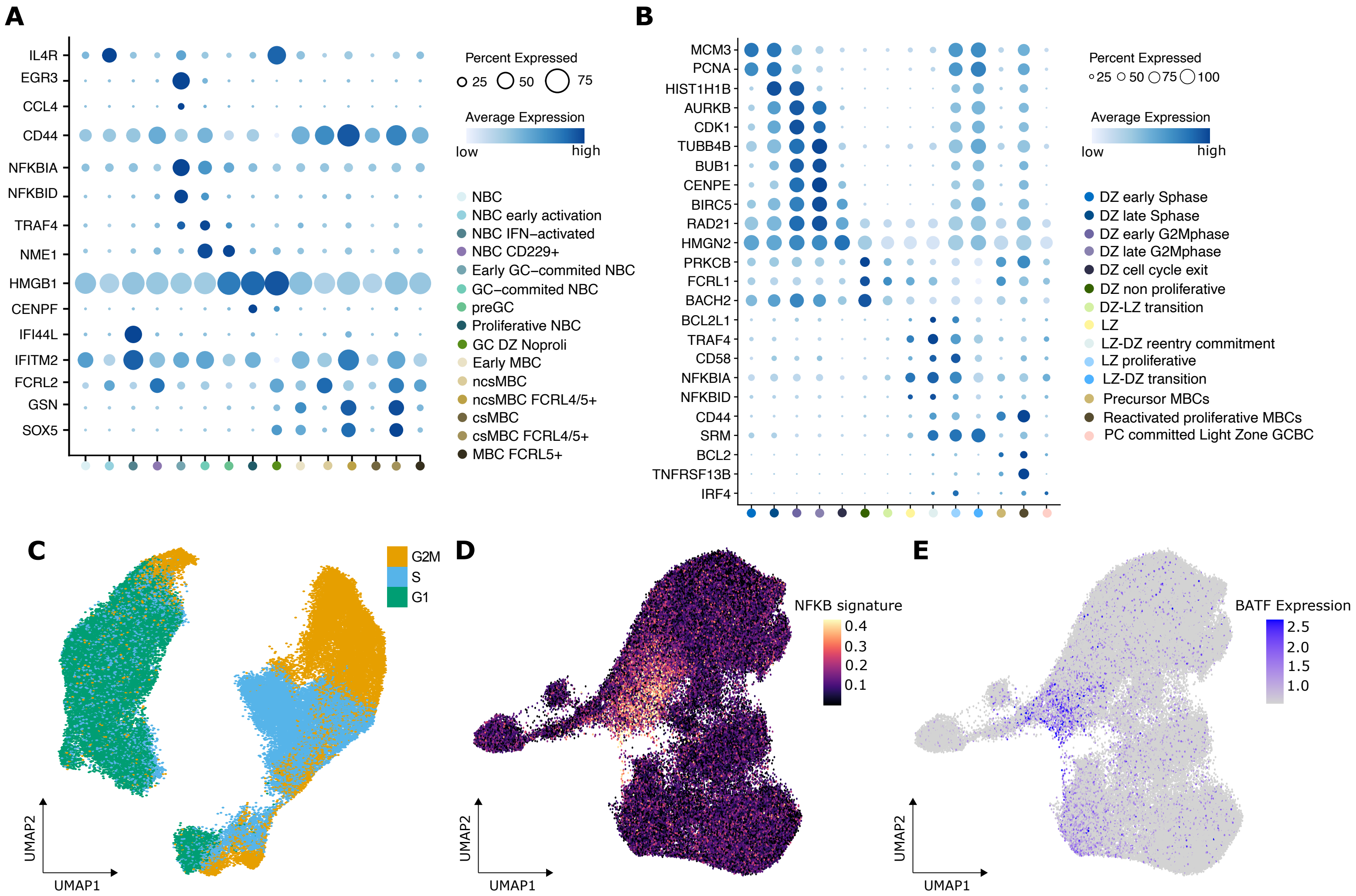
A
IL4R
EGR3
CCL4
CD44
NFKBIA
NFKBID
TRAF4
NME1
HMGB1
CENPF
IFI44L
IFITM2
FCRL2
GSN
SOX5
Percent Expressed
25
50
75
Average Expression
low
high
NBC
NBC early activation
NBC IFN–activated
NBC CD229+
Early GC–commited NBC
GC–commited NBC
preGC
Proliferative NBC
GC DZ Noproli
Early MBC
ncsMBC
ncsMBC FCRL4/5+
csMBC
csMBC FCRL4/5+
MBC FCRL5+
B
MCM3
PCNA
HIST1H1B
AURKB
CDK1
TUBB4B
BUB1
CENPE
BIRC5
RAD21
HMGN2
PRKCB
FCRL1
BACH2
BCL2L1
TRAF4
CD58
NFKBIA
NFKBID
CD44
SRM
BCL2
TNFRSF13B
IRF4
Percent Expressed
25
50
75
100
Average Expression
low
high
DZ early Sphase
DZ late Sphase
DZ early G2Mphase
DZ late G2Mphase
DZ cell cycle exit
DZ non proliferative
DZ-LZ transition
LZ
LZ-DZ reentry commitment
LZ proliferative
LZ-DZ transition
Precursor MBCs
Reactivated proliferative MBCs
PC committed Light Zone GCBC
C
G2M
S
G1
UMAP2
UMAP1
D
NFKB signature
0.4
0.3
0.2
0.1
UMAP2
UMAP1
E
BATF Expression
2.5
2.0
1.5
1.0
UMAP2
UMAP1

#### Figure S7

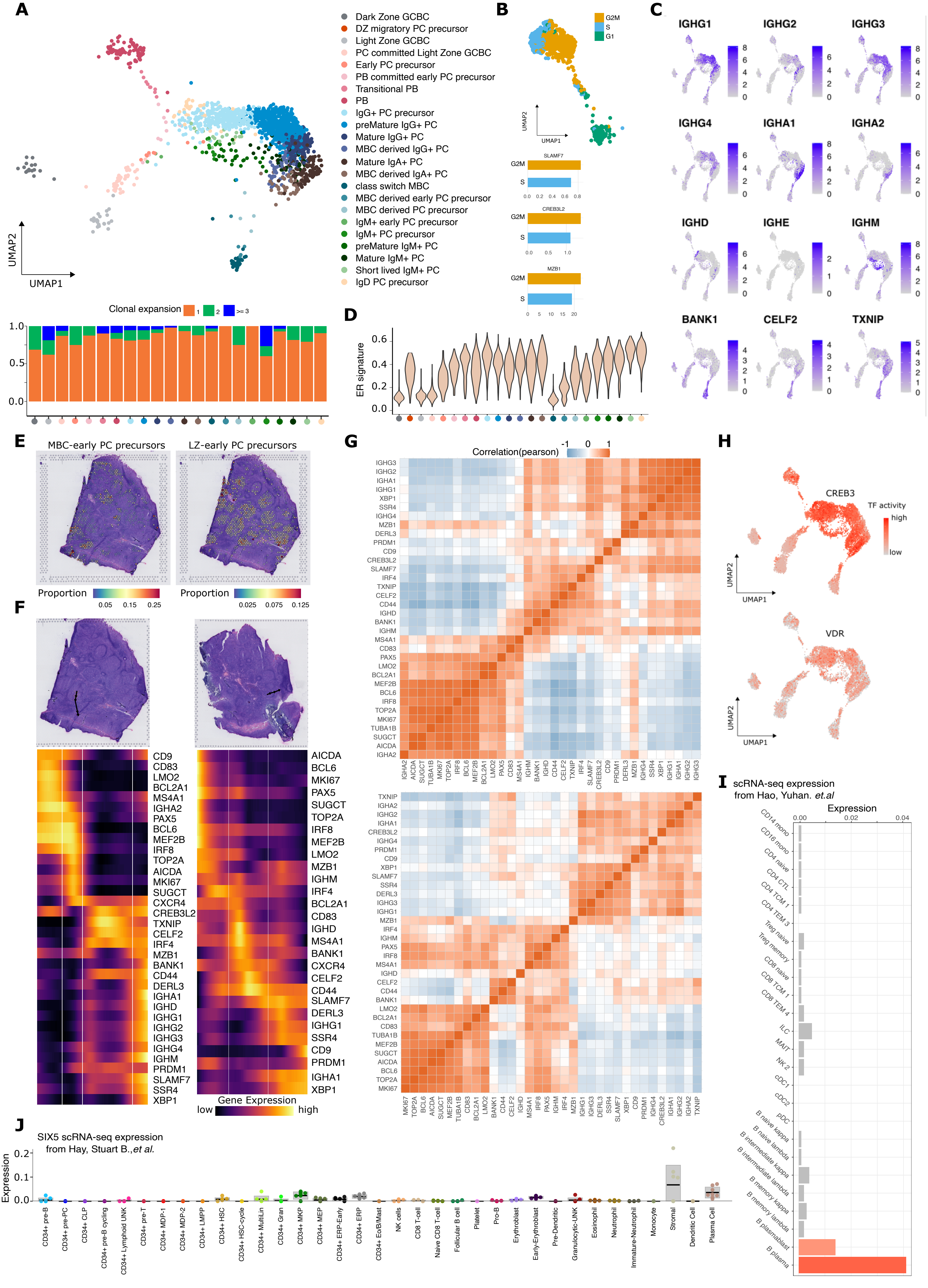
A
Dark Zone GCBC
DZ migratory PC precursor
Light Zone GCBC
PC committed Light Zone GCBC
Early PC precursor
PB committed early PC precursor
Transitional PB
PB
IgG+ PC precursor
preMature IgG+ PC
Mature IgG+ PC
MBC derived IgG+ PC
Mature IgA+ PC
MBC derived IgA+ PC
class switch MBC
MBC derived early PC precursor
MBC derived PC precursor
IgM+ early PC precursor
IgM+ PC precursor
preMature IgM+ PC
Mature IgM+ PC
Short lived IgM+ PC
IgD PC precursor
UMAP1
UMAP2
Clonal expansion 1 2 >=3
B
G2M
S
G1
SLAMF7
CREB3L2
MZB1
C
IGHG1
IGHG2
IGHG3
IGHG4
IGHA1
IGHA2
IGHD
IGHE
IGHM
BANK1
CELF2
TXNIP
D
ER signature
E
MBC-early PC precursors
LZ-early PC precursors
Proportion
F
Gene Expression
low
high
G
Correlation(pearson)
H
CREB3
VDR
TF activity
high
low
I
scRNA-seq expression from Hao, Yuhan. et.al
Expression
CD14 mono
CD16 mono
CD4 naive
CD4 CTL
CD4 TCM 1
CD4 TEM 3
Treg naive
Treg memory
CD8 naive
CD8 TCM 1
CD8 TEM 4
ILC
MAIT
NK 2
cDC1
cDC2
pDC
B naive kappa
B naive lambda
B intermediate kappa
B intermediate lambda
B memory kappa
B memory lambda
B plasmablast
B plasma
J
SIX5 scRNA-seq expression from Hay, Stuart B.,et al.
Expression
CD34+ pre-B
CD34+ pre-PC
CD34+ CLP
CD34+ pre-B cycling
CD34+ Lymphoid UNK
CD34+ pre-T
CD34+ MDP-1
CD34+ MDP-2
CD34+ LMPP
CD34+ HSC
CD34+ HSC-cycle
CD34+ MultiLin
CD34+ Gran
CD34+ MKP
CD34+ MEP
CD34+ ERP-Early
CD34+ ERP
CD34+ Eo/B/Mast
NK cells
CD8 T-cell
Naive CD8 T-cell
Follicular B cell
Platelet
Pro-B
Erythroblast
Early-Erythroblast
Pre-Dendritic
Granulocytic-UNK
Eosinophil
Neutrophil
Immature-Neutrophil
Monocyte
Stromal
Dendritic Cell
Plasma Cell

### Figure S8

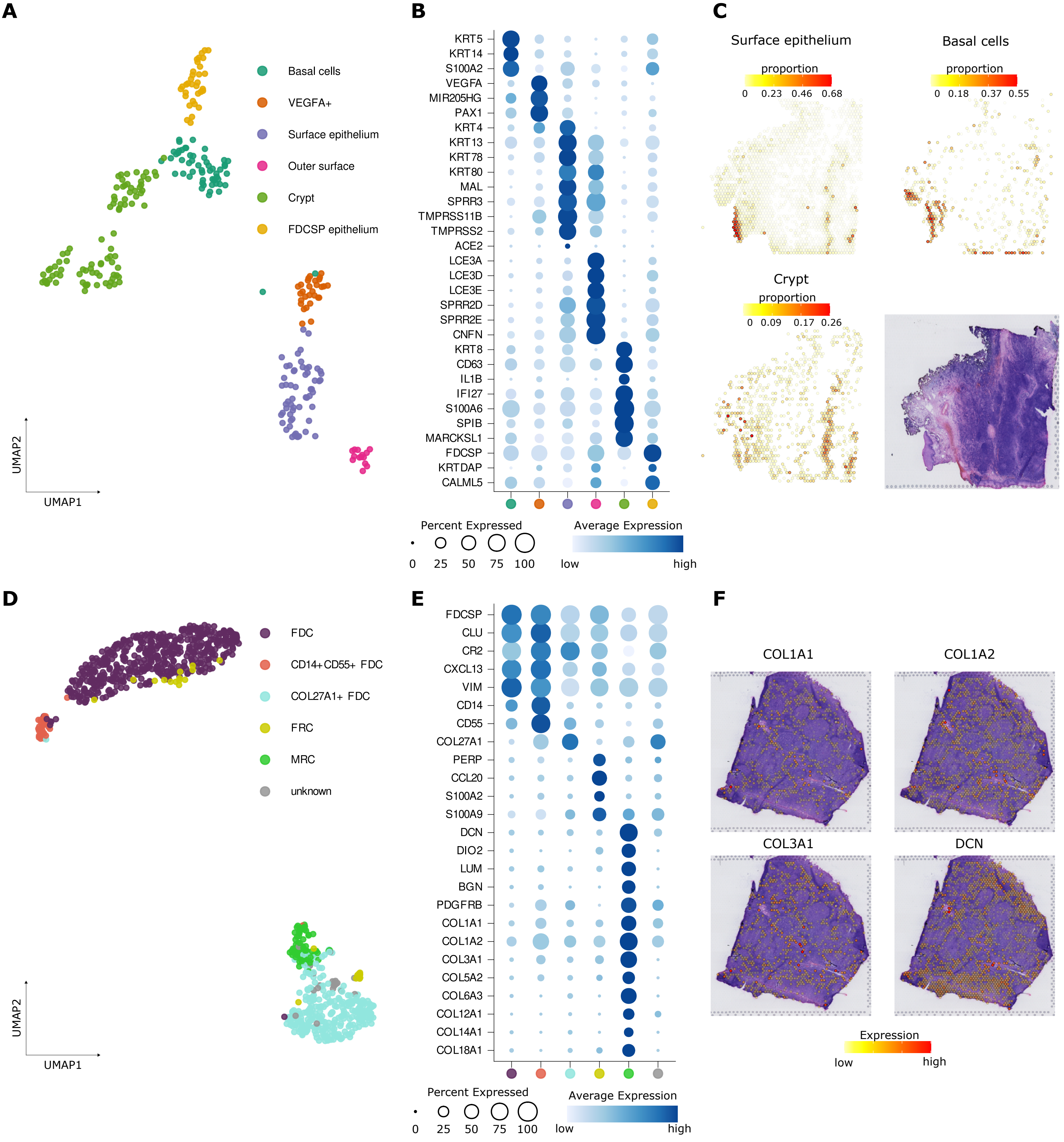

A
Basal cells
VEGFA+
Surface epithelium
Outer surface
Crypt
FDCSP epithelium
UMAP2
UMAP1
B
KRT5
KRT14
S100A2
VEGFA
MIR205HG
PAX1
KRT4
KRT13
KRT78
KRT80
MAL
SPRR3
TMPRSS11B
TMPRSS2
ACE2
LCE3A
LCE3D
LCE3E
SPRR2D
SPRR2E
CNFN
KRT8
CD63
IL1B
IFI27
S100A6
SPIB
MARCKSL1
FDCSP
KRTDAP
CALML5
Percent Expressed
0 25 50 75 100
Average Expression
low high
C
Surface epithelium
proportion
0 0.23 0.46 0.68
Basal cells
proportion
0 0.18 0.37 0.55
Crypt
proportion
0 0.09 0.17 0.26
D
FDC
CD14+CD55+ FDC
COL27A1+ FDC
FRC
MRC
unknown
UMAP2
UMAP1
E
FDCSP
CLU
CR2
CXCL13
VIM
CD14
CD55
COL27A1
PERP
CCL20
S100A2
S100A9
DCN
DIO2
LUM
BGN
PDGFRB
COL1A1
COL1A2
COL3A1
COL5A2
COL6A3
COL12A1
COL14A1
COL18A1
Percent Expressed
0 25 50 75 100
Average Expression
low high
F
COL1A1
COL1A2
COL3A1
DCN
Expression
low high

### Figure S9

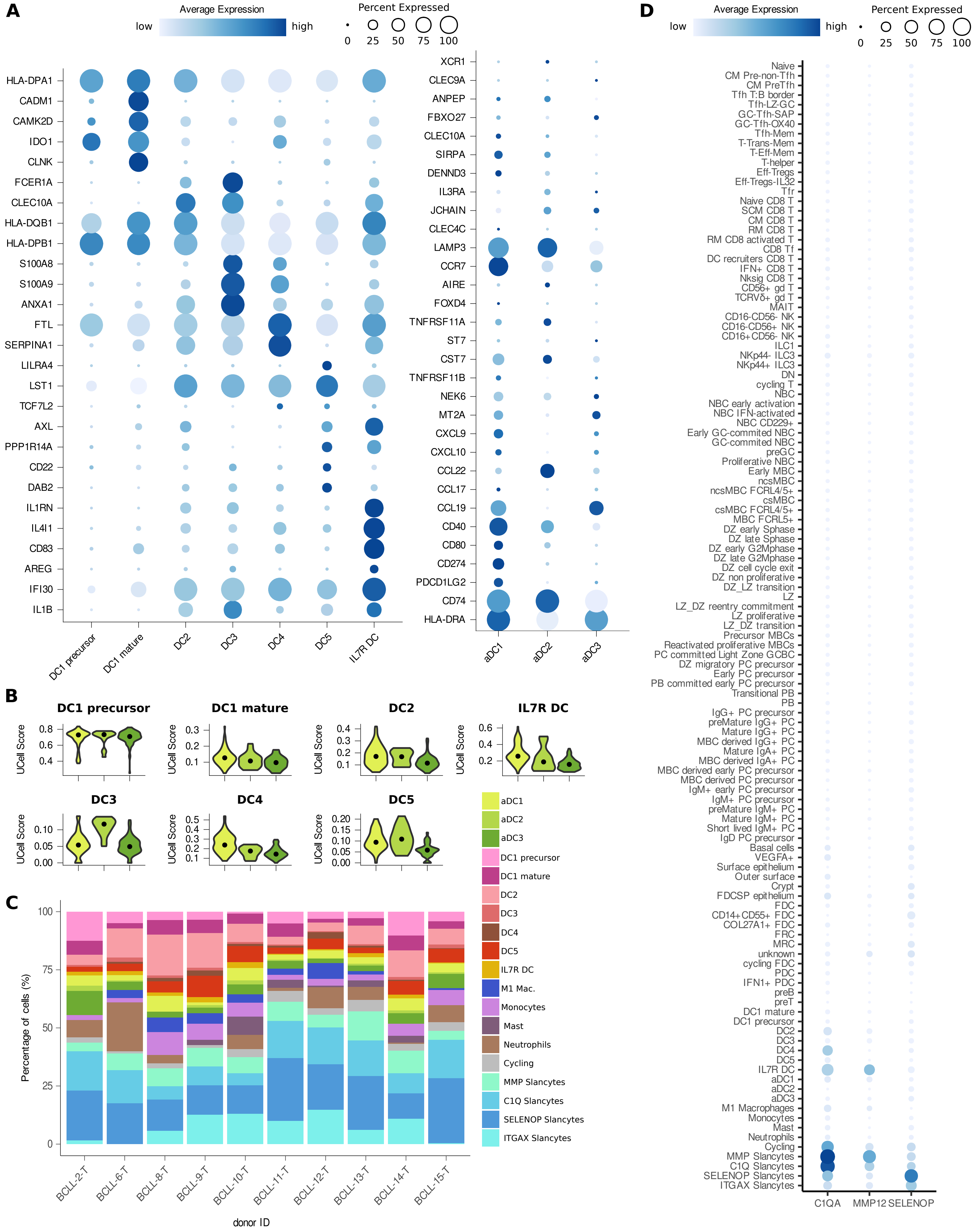
A
Average Expression
low
high
Percent Expressed
0
25
50
75
100
HLA-DPA1
CADM1
CAMK2D
IDO1
CLNK
FCER1A
CLEC10A
HLA-DQB1
HLA-DPB1
S100A8
S100A9
ANXA1
FTL
SERPINA1
LILRA4
LST1
TCF7L2
AXL
PPP1R14A
CD22
DAB2
IL1RN
IL4I1
CD83
AREG
IFI30
IL1B
DC1 precursor
DC1 mature
DC2
DC3
DC4
DC5
IL7R DC
XCR1
CLEC9A
ANPEP
FBXO27
CLEC10A
SIRPA
DENND3
IL3RA
JCHAIN
CLEC4C
LAMP3
CCR7
AIRE
FOXD4
TNFRSF11A
ST7
CST7
TNFRSF11B
NEK6
MT2A
CXCL9
CXCL10
CCL22
CCL17
CCL19
CD40
CD80
CD274
PDCD1LG2
CD74
HLA-DRA
aDC1
aDC2
aDC3
B
DC1 precursor
DC1 mature
DC2
IL7R DC
DC3
DC4
DC5
UCell Score
aDC1
aDC2
aDC3
DC1 precursor
DC1 mature
DC2
DC3
DC4
DC5
IL7R DC
M1 Mac.
Monocytes
Mast
Neutrophils
Cycling
MMP Slancytes
C1Q Slancytes
SELENOP Slancytes
ITGAX Slancytes
C
Percentage of cells (%)
0
25
50
75
100
BCLL-2-T
BCLL-6-T
BCLL-8-T
BCLL-9-T
BCLL-10-T
BCLL-11-T
BCLL-12-T
BCLL-13-T
BCLL-14-T
BCLL-15-T
donor ID
D
Average Expression
low
high
Percent Expressed
0
25
50
75
100
Naive
CM Pre-non-Tfh
CM PreTfh
Tfh T:B border
Tfh-LZ-GC
GC-Tfh-SAP
GC-Tfh-OX40
Tfh-Mem
T-Trans-Mem
T-Eff-Mem
T-helper
Eff-Tregs
Eff-Tregs-IL32
Tfr
Naive CD8 T
SCM CD8 T
CM CD8 T
RM CD8 T
RM CD8 activated T
CD8 Tf
DC recruiters CD8 T
IFN+ CD8 T
Nksig CD8 T
CD56+ gd T
TCRVδ+ gd T
MAIT
CD16-CD56- NK
CD16-CD56+ NK
CD16+CD56- NK
ILC1
NKp44- ILC3
NKp44+ ILC3
DN
cycling T
NBC
NBC early activation
NBC IFN-activated
NBC CD229+
Early GC-commited NBC
GC-commited NBC
preGC
Proliferative NBC
Early MBC
ncsMBC
ncsMBC FCRL4/5+
csMBC
csMBC FCRL4/5+
MBC FCRL5+
DZ early Sphase
DZ late Sphase
DZ early G2Mphase
DZ late G2Mphase
DZ cell cycle exit
DZ non proliferative
DZ_LZ transition
LZ
LZ_DZ reentry commitment
LZ proliferative
LZ_DZ transition
Precursor MBCs
Reactivated proliferative MBCs
PC committed Light Zone GCBC
DZ migratory PC precursor
Early PC precursor
PB committed early PC precursor
Transitional PB
PB
IgG+ PC precursor
preMature IgG+ PC
Mature IgG+ PC
MBC derived IgG+ PC
Mature IgA+ PC
MBC derived IgA+ PC
MBC derived early PC precursor
MBC derived PC precursor
IgM+ early PC precursor
IgM+ PC precursor
preMature IgM+ PC
Mature IgM+ PC
Short lived IgM+ PC
IgD PC precursor
Basal cells
VEGFA+
Surface epithelium
Outer surface
Crypt
FDCSP epithelium
FDC
CD14+CD55+ FDC
COL27A1+ FDC
FRC
MRC
unknown
cycling FDC
PDC
IFN1+ PDC
preB
preT
DC1 mature
DC1 precursor
DC2
DC3
DC4
DC5
IL7R DC
aDC1
aDC2
aDC3
M1 Macrophages
Monocytes
Mast
Neutrophils
Cycling
MMP Slancytes
C1Q Slancytes
SELENOP Slancytes
ITGAX Slancytes
C1QA
MMP12
SELENOP

### Figure S10

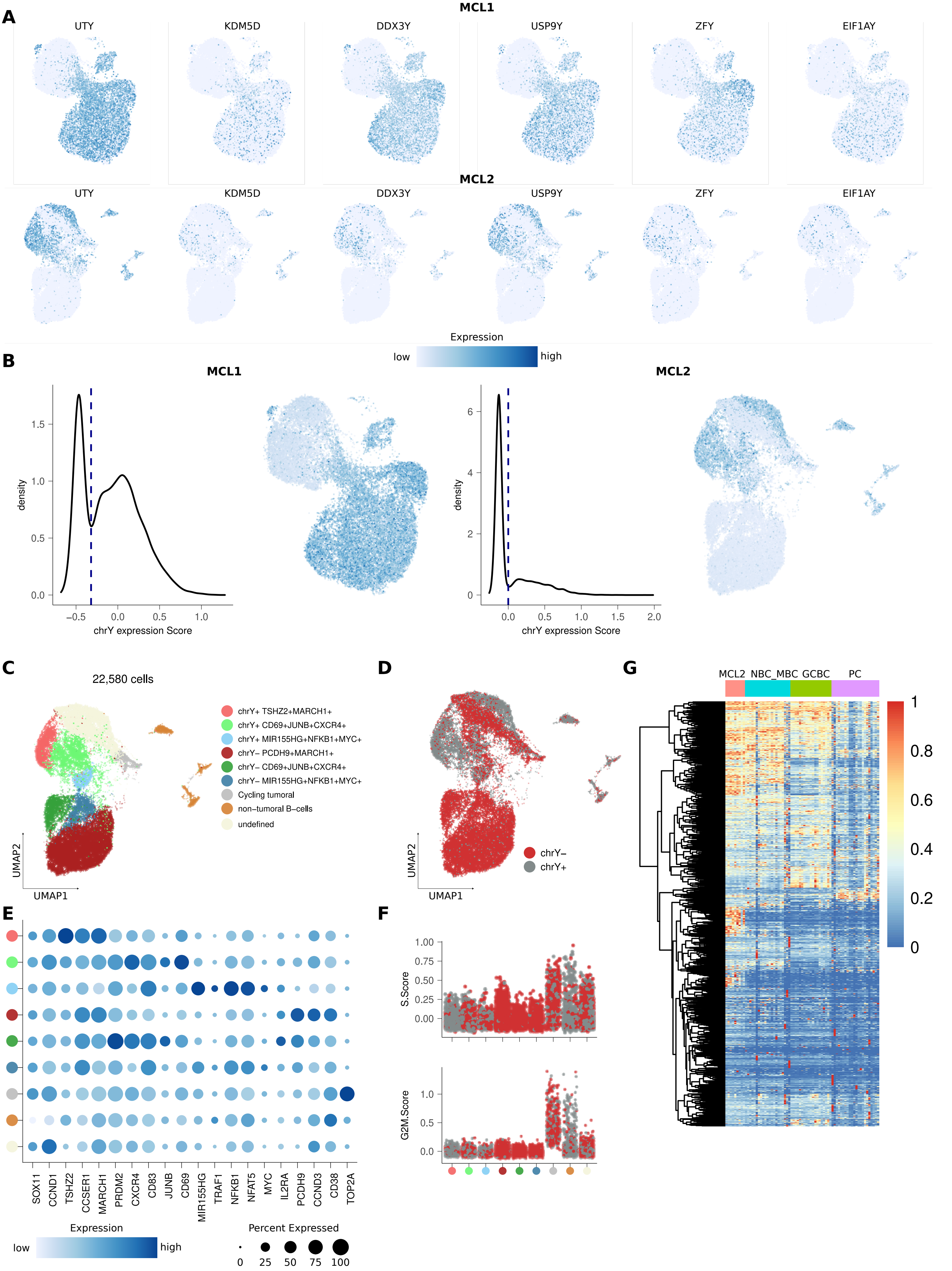

A
MCL1
UTY
KDM5D
DDX3Y
USP9Y
ZFY
EIF1AY
MCL2
Expression
low
high
B
MCL1
MCL2
density
chrY expression Score
C
22,580 cells
chrY+ TSHZ2+MARCH1+
chrY+ CD69+JUNB+CXCR4+
chrY+ MIR155HG+NFKB1+MYC+
chrY− PCDH9+MARCH1+
chrY− CD69+JUNB+CXCR4+
chrY− MIR155HG+NFKB1+MYC+
Cycling tumoral
non−tumoral B−cells
undefined
UMAP2
UMAP1
D
chrY−
chrY+
G
MCL2 NBC_MBC GCBC PC
E
SOX11
CCND1
TSHZ2
CCSER1
MARCH1
PRDM2
CXCR4
CD83
JUNB
CD69
MIR155HG
TRAF1
NFKB1
NFAT5
MYC
IL2RA
PCDH9
CCND3
CD38
TOP2A
Percent Expressed
F
S.Score
G2M.Score
