## Supplementary material for "An Atlas of Cells in the Human Tonsil": Methods

STAR Methods: An Atlas of Cells in the Human Tonsil

### KEY RESOURCES TABLE

#### Software and Algorithms

| Cellranger-atac v1.2 | CellRanger ATAC (10X Genomics) | https://support.10xgenomics.com/single-cell-atac/software/overview/welcome |
| --- | --- | --- |
| Cellranger v4.0.0 | CellRanger (10X Genomics) | https://support.10xgenomics.com/single-cell-gene-expression/software/overview/welcome |
| Cellranger-arc v1.0 | CellRanger ARC (10X Genomics) | https://support.10xgenomics.com/single-cell-multiome-atac-gex/software/overview/welcome |
| Cellranger-multi v6.0.1 | CellRanger multi (10X Genomics) | https://support.10xgenomics.com/single-cell-vdj/software/pipelines/latest/using/multi |
| Spaceranger 1.1.0 | 10X Genomics | https://support.10xgenomics.com/spatial-gene-expression/software/overview/welcome |
| Seurat | (Hao et al., 2021) | https://satijalab.org/seurat/ |
| Signac | (Stuart et al., 2021) | https://satijalab.org/signac/ |
| Harmony | (Korsunsky et al., 2019) | https://github.com/immunogenomics/harmony |
| clusterProfiler | (Wu et al., 2021) | https://yulab-smu.top/biomedical-knowledge-mining-book/ |
| UCell | (Andreatta and Carmona, 2021) | https://bioconductor.org/packages/release/bioc/html/UCell.html |
| MACS2 | (Zhang et al., 2008) | https://github.com/macs3-project/MACS |
| ChromVar | (Schep et al., 2017) | http://www.bioconductor.org/packages/release/bioc/html/chromVAR.html |
| Cicero | (Pliner et al., 2018) | [http://cole-trapnell-lab.github.io/cicero-release](http://cole-trapnell-lab.github.io/cicero-release/) |
| pySCENIC v0.10.3 | (Van de Sande et al., 2020) | https://pyscenic.readthedocs.io/en/latest/ |
| Vireo | (Huang et al., 2019) | https://github.com/single-cell-genetics/vireo |
| cellSNP | (Huang and Huang, 2021) | https://github.com/single-cell-genetics/cellSNP |
| Scirpy | (Sturm et al., 2020) | https://github.com/scverse/scirpy |
| SPOTlight | (Elosua-Bayes et al., 2021) | https://github.com/MarcElosua/SPOTlight |
| MAGIC | (van Dijk et al., 2018) | https://github.com/KrishnaswamyLab/MAGIC |
| SPATA2 | (Kueckelhaus et al., 2020) | https://github.com/theMILOlab/SPATA2 |
| LISI | (Korsunsky et al., 2019) | https://github.com/immunogenomics/LISI |
| Scrublet | (Wolock et al., 2019) | https://github.com/swolock/scrublet |
| CellphoneDBv3 | (Garcia-Alonso et al., 2021) | https://github.com/ventolab/CellphoneDB |
| R | R core | [https://www.r-project.org](https://www.r-project.org/) |
| Python | Python Software Foundation | https://www.python.org |
| Shiny App | This paper | <https://singlecellgenomics-cnag-crg.shinyapps.io/Annotation/> |
| HCATonsilData | This paper | https://github.com/massonix/HCATonsilData |
| SLOcatoR | This paper | https://github.com/massonix/SLOcatoR/tree/main/R |
| iSee instance | This paper | http://shiny.imbei.uni-mainz.de:3838/iSEE_TonsilDataAtlas/ |

#### Others

| Scenic TF Database | (Van de Sande et al., 2020) | <https://github.com/aertslab/SCENICprotocol/blob/master/example/allTFs_hg38.txt> |
| --- | --- | --- |
| CisTarget databases  Hg38__refseq-r80__500bp_up_and_100bp_down_tss.mc9nr.feather  motifs-v9-nr.hgnc-m0.001-o0.0.tbl | (Herrmann et al., 2012; Imrichová et al., 2015) | <https://resources.aertslab.org/cistarget/> |
| JASPAR2020 | (Fornes et al., 2020; Tan and Lenhard, 2016) | http://bioconductor.org/packages/release/data/annotation/vignettes/JASPAR2020/inst/doc/JASPAR2020.html |
| chromVARmotifs | (Schep et al., 2017) | https://github.com/GreenleafLab/chromVARmotifs |

#### Deposited data

| Raw and analyzed data | This paper | EGA: EGAS00001006375 |
| --- | --- | --- |
| Outputs CellRanger | This paper | <https://doi.org/10.5281/zenodo.6678331> |
| Seurat objects | This paper | <https://doi.org/10.5281/zenodo.6340174> |
| Code and vignettes | This paper | https://github.com/Single-Cell-Genomics-Group-CNAG-CRG/TonsilAtlas |

### RESOURCE AVAILABILITY

#### Lead contact

Requests for further information or access to data should be directed to Holger Heyn.

#### Materials availability

This study did not generate new unique reagents.

#### Data and code availability

The data has been deposited in five levels of organization, from raw to processed data:

- Level 1: raw data. All fastq files for all data modalities have been deposited at the European Genome Archive (EGA) under accession id EGAS00001006375.
- Level 2: matrices. All data modalities correspond to different technologies from 10X Genomics. As such, they were mapped with different flavors of CellRanger (CR). The most important files in the “outs” folder of every CR run (including all matrices) have been deposited in Zenodo (<https://doi.org/10.5281/zenodo.6678331>).
- Level 3: Seurat Objects. All data was analyzed within the Seurat ecosystem (Hao et al., 2021). We have archived in Zenodo (<https://doi.org/10.5281/zenodo.6340174>) all Seurat Objects that contain the raw and processed counts, dimensionality reductions (PCA, Harmony, UMAP), and metadata needed to reproduce all figures from this manuscript.
- Level 4: to allow for programmatic and modular access to the whole tonsil atlas dataset, we developed HCATonsilData, available on GitHub: <https://github.com/massonix/HCATonsilData>. HCATonsilData provides a vignette which documents how to navigate and understand the data. In addition, we will periodically update the annotations as we refine it with suggestions from the community.
- Level 5: interactive mode. We provide different iSEE instances to browse the data interactively: <http://shiny.imbei.uni-mainz.de:3838/iSEE_TonsilDataAtlas/>.

All code related with this publication is available on GitHub:

- Scripts and notebooks to reproduce all analysis: <https://github.com/Single-Cell-Genomics-Group-CNAG-CRG/TonsilAtlas>. Most analysis notebooks have a companion html report that has all the plots that motivate the thresholds and parameters used in these analyses.
- SLOcatoR package: <https://github.com/massonix/SLOcatoR>.
- Shiny app used to annotate cells: <https://singlecellgenomics-cnag-crg.shinyapps.io/Annotation/>.
- Shiny app used to annotate histology slides: <https://github.com/Single-Cell-Genomics-Group-CNAG-CRG/shiny-pathology>.
- Code to generate iSEE instances: <https://github.com/iSEE/iSEE_instances/tree/master/iSEE_HCATonsilData>.

### EXPERIMENTAL MODEL AND SUBJECT DETAILS

#### Sample collection and processing

###### For the current study, ten tonsil samples were obtained from healthy donors from three different age groups, *i.e.*, children (age 3-5, 3 males and 3 females), 3 young male adults (age 26-35) and old male adult (age 65). Tonsils of children and two young adults cases undergoing tonsillectomies were obtained from the Clinica Universidad de Navarra (Pamplona, Spain) undergoing simultaneous adenoidectomy and tonsillectomy due to recurrent respiratory infections. All donors gave informed consent for their participation in this study, which was approved by the clinical research ethics committee of Clínica Universidad de Navarra. Additionally, one tonsil each of a young adult and an old adult undergoing tonsillectomies were obtained from Hospital Clinic of Barcelona (Spain), following the informed consent of both donors for their participation in this study. The Tonsil@tlas project was approved by the clinical research ethics committee of the Hospital Clinic of Barcelona (HCB/2018/0992). For the final part of the study, two cryopreserved tonsil samples of mantle cell lymphoma (MCL) patients were used. Informed consents were obtained according to the Institutional Review Board of the Hospital Clínic of Barcelona following the International Cancer Genome Consortium guidelines.

Tonsil tissue of healthy donors was split into three parts that were processed as follows: (1) first part was paraffin embedded to create FFPE blocks, according to standard pathology protocols; (2) second part was snap frozen to obtain OCT blocks, according to standard protocols, and (3) the remaining third part was processed to obtain a single-cell suspension, following the steps described below. Tonsils were first disaggregated by extensive manual mincing and filtered by 70 µm nylon strainer. In case non-disaggregated parts of tissue were still present, the samples were further dissociated by gentle MACS Dissociator (program tumor 04.01). The number and viability of cells was evaluated. All steps were performed at 4ºC or on ice. Cells were either processed directly for single-cell sequencing (scRNA-seq, scATAC-seq, CITE-seq, or Multiome) or cryopreserved for later use. MCL samples were obtained from cryopreserved dissociated cells from tonsils, from the ICGC case collection of Hospital Clinic, Barcelona. After thawing in culture medium supplemented with 20% FBS, the CD19 positive B cell fraction and CD19 negative non-B cell fraction were isolated by magnetic cell separation using CD19-MicroBeads MACS separation system protocol (Miltenyi Biotec, Auburn, CA). Separation steps were performed at 4ºC. Both fractions were directly processed for Multiome library preparation and sequenced separately.

### METHOD DETAILS

#### 3’ scRNA-seq and Cell hashing

###### Freshly isolated cells from tonsils were subjected to a Cell Hashing (Stoeckius et al., 2018) protocol before proceeding to scRNA-seq. Cell hashing was performed following manufacturer’s instructions (Cell hashing and Single Cell Proteogenomics Protocol Using TotalSeq™ Antibodies; BioLegend). Cells were counted with a TC20™ Automated Cell Counter (Bio-Rad Laboratories, S.A), and 50,000 unlabeled cells were saved in a separate tube before proceeding with the cell hashing protocol. Each sample was split into seven aliquots with equal numbers of cells. Briefly, each aliquot was resuspended in Cell Staining Buffer (BioLegend), incubated for 10 min at 4°C with Human TruStain FcX™ Fc Blocking reagent (BioLegend). To each aliquot, a specific TotalSeq-A antibody-oligo conjugate (Tables S17) was added and incubated on ice for 30 min. Cells were then washed three times with cold PBS+0.05% BSA (ThermoFisher) and centrifuged for 5 min at 500 rcf at 4°C. Finally, cells were resuspended in an appropriate volume of PBS+0.05% BSA to obtain a final cell concentration >1000 cells/µl, suitable for scRNA-seq. Assuming a 50% loss of cells in all tubes, an equal volume of hashed cell suspension from each of the seven aliquots was mixed and filtered with a 40 µm strainer. Cell concentration was verified with a TC20™ Automated Cell Counter (Bio-Rad Laboratories, S.A) upon cell staining with Trypan Blue.

###### Cells were partitioned into Gel Beads-in-emulsion (GEMs) by using the Chromium Controller system (10x Genomics). Each sample was loaded into two channels with a target recovery of 20,000 cells per channel, for a total final recovery of 40,000 cells per sample. To assess the potential effects of cell hashing on gene expression and cell type composition, we additionally added non-hashed cells as a control (TR=5,000 cells). cDNA sequencing libraries were prepared using the Next GEM Single Cell 3’ Reagent Kits v3.1 (10x Genomics, PN-1000121), with some adaptations for Cell hashing, as indicated in TotalSeq™-A antibodies and Cell Hashing with 10x Single Cell 3' Reagent Kit v3 3.1 protocol by BioLegend. Briefly, 1 µl of 0.2 µM hashtag oligonucleotides (HTO) primer (Integrated DNA Technologies, IDT) was added to the cDNA amplification reaction to amplify the HTO together with the full-length cDNAs. A SPRI selection clean up was done in order to separate mRNA-derived cDNA (>300 bp) from antibody-oligo-derived cDNA (<180 bp), as described in the above mentioned protocol form BioLegend. Gene Expression (GEX) libraries were prepared following 10X Genomics single-cell 3’ mRNA kit protocol, while HTO DNAs were indexed by PCR as follows. 5 µl of purified HTO DNA were mixed with 2.5 µl of 10 µM Illumina TruSeq D70X_s primer (IDT) carrying a different i7 index for each sample (Tables S18), 2.5 µl of SI primer from 10X single-cell 3’ mRNA kit, 50 µl of 2 X KAPA HiFi PCR Master Mix (KAPA Biosystem) and 40 µl of nuclease-free water. The reaction was carried out using the following thermal cycling conditions: 98°C for 2 min (initial denaturation), 12 cycles of 98°C for 20 seconds, 64°C for 30 seconds, 72°C for 20 seconds, and a final extension at 72°C for 5 min. The HTO libraries were purified by adding 1.2 X SPRI select reagent to the PCR reaction, incubating 5 min at room temperature (RT) and removing the supernatant after capturing the beads with a magnet. Samples were washed two times with 80% ethanol and elution was performed by adding 40.5 µl of nuclease-free water to the beads.

Size distribution and concentration of full-length cDNA and HTO libraries were verified on an Agilent Bioanalyzer High Sensitivity chip (Agilent Technologies). Finally, sequencing of HTO and GEX libraries was carried out on a NovaSeq 6000 sequencer (Illumina) using the following sequencing conditions: 28 bp (Read 1) + 8 bp (i7 index) + 0 bp (i5 index) + 89 bp (Read 2), to obtain approximately 2,000 and >20,000 paired-end reads per HTO and cell, respectively.

#### scATAC-seq

###### Cryopreserved samples were rapidly thawed in a 37ºC water bath and transferred to a 15 ml Falcon using a 1000 µl wide bore tip. Next, 1 ml of 37ºC pre-warmed Hibernate-A media supplemented with 10% FBS (Thermo Fisher Scientific) was added dropwise while gently swirling the sample. After 1 min RT incubation, 2 ml of pre-warmed media were added as mentioned before. Samples were again incubated at RT for 1 min and then additional media was added to bring the volume to 15 ml. Samples were centrifuged at 500 x g for 5 min at RT. Supernatant was removed and pellets resuspended in 10 ml of 1X PBS (Thermo Fisher Scientific) supplemented with 1% BSA. Samples were filtered with a 40 µm cell strainer to remove clumps. Cell number and viability were verified with a TC20™ Automated Cell Counter (Bio Rad). Dead cells were removed by FACS sorting DAPI negative cells using a FACSAria™ Fusion Flow Cytometer (BD Biosciences). In order to determine potential biases introduced by cell sorting, unsorted cells from 4 out of 10 samples were used for scATAC-seq analysis and compared with corresponding sorted samples.

###### Nuclei isolation was performed following the “Nuclei Isolation for Single Cell ATAC Sequencing” demonstrated protocol (10X Genomics; CG000169) starting from 1 million cells per sample and incubating on ice for 3 min for cell lysis. Based on the starting number of cells and assuming a 50% loss during the procedure, nuclei were resuspended into the appropriate volume of chilled Diluted Nuclei Buffer (10x Genomics) in order to achieve a nuclei concentration of 925-2300 nuclei/µl, suitable for a Target Nuclei Recovery of 5,000 nuclei per sample. The resulting nuclei concentration was determined by manual counting using a Neubauer chamber upon staining with Trypan Blue**.**

###### scATAC-seq libraries were prepared according to the Chromium Single Cell ATAC Reagent Kits v1.1 User Guide (10x Genomics; CG000209 Rev D). Transposed nuclei were partitioned into GEMs by using the Chromium Controller with Chip H for a target recovery of 5000 nuclei per sample. For samples used to assess potential artifacts due to FACS sorting, nuclei obtained from sorted and unsorted cells were loaded on separate channels of the same chip and parallelly processed for library preparation. After linear amplification, the resulting DNA was purified by sequential Dynabeads and SPRIselect reagent beads clean-ups. Libraries were indexed by PCR using the Single Index Kit N Set A (10x Genomics, PN-1000212) applying 10 cycles of amplification. Sequencing libraries were subjected to a final bead clean-up SPRIselect reagent and quantified on an Agilent Bioanalyzer High Sensitivity chip (Agilent Technologies). Finally, libraries were loaded on an Illumina NovaSeq 6000 with the following sequencing conditions: 50 bp (Read 1N) + 8 bp (i7 Index) + 16 bp (i5 Index) + 49 bp (Read 2N), aiming at a sequencing depth of >25,000 reads/nucleus.

#### Single cell RNA and chromatin accessibility profiling

###### Cryopreserved cells were thawed and cleaned from dead cells as described before (see scATAC-seq). Nuclei isolation was performed following the “Nuclei Isolation for Single Cell Multiome ATAC + Gene Expression Sequencing'' demonstrated protocol (10x Genomics; CG000365) starting from 0.5-1 M cells per sample and incubating on ice during 3 min for cell lysis. Nuclei were resuspended into the appropriate volume of chilled Diluted Nuclei Buffer (10X Genomics) to achieve a nuclei concentration of 925-2,300 nuclei/µl, suitable for a Target Nuclei Recovery of 7,000 per sample. The resulting nuclei concentration was determined by manual counting using a Neubauer chamber upon staining with Trypan Blue.

###### GEX and ATAC-seq libraries were prepared following the Chromium Next GEM Single Cell Multiome ATAC + Gene Expression User Guide (10x Genomics; CG000338). Transposed nuclei were partitioned into GEMs by using the Chromium Controller with Chip J aiming at a target recovery of 7,000 nuclei per sample. After GEMs incubation for mRNA reverse transcription and transposed DNA barcoding, the resulting cDNA and barcoded gDNA were purified and pre-amplified during 7 cycles, following the 10x Genomics protocol. After a clean-up, 35 µl of the pre-amplified cDNA were amplified with 7 additional PCR cycles. The resulting cDNA was quantified on an Agilent Bioanalyzer High Sensitivity chip (Agilent Technologies) and 100 ng were used for library preparation. GEX libraries were indexed with 13 cycles of amplification using the Dual Index Plate TT Set A (10x Genomics; PN-3000431). In parallel, 40 µl of the pre-amplified DNA were indexed with 7 cycles of amplification using the Sample Index N Set A (10X Genomics; PN 3000427). Size distribution and concentration of full-length GEX and ATAC-seq libraries were verified on an Agilent Bioanalyzer High Sensitivity chip (Agilent Technologies). Finally, sequencing of GEX libraries was carried out on a NovaSeq 6000 sequencer (Illumina) using the following sequencing conditions: 28 bp (Read 1) + 8 bp (i7 index) + 0 bp (i5 index) + 89 bp (Read 2), to obtain approximately >20,000 paired-end reads per cell. ATAC-seq libraries were also sequenced with a NovaSeq 6000 sequencer (Illumina) using the following conditions: 50 bp (Read 1N) + 8 bp (i7 Index) + 16 bp (i5 Index) + 49 bp (Read 2N), aiming at a sequencing depth of >25,000 reads/nucleus.

#### CITE-Seq

###### Cryopreserved cells were thawed and FACS sorted as previously described (see scATAC-seq and 3’ Single Cell RNA sequencing). For CITE-Seq experiments, samples were processed separately or processed in pools (subsequent demultiplexing by genotypes) before cell labeling with a custom panel of 192 oligo-barcoded antibodies (TotalSeq-C Custom Human Panel, Biolegend), following the same staining protocol of for Cell Hashing (see 3’ Single Cell RNA sequencing). Antibody details are included in Tables S19. Cells were loaded on the 10x Chromium Controller using the Next GEM Single Cell V(D)J Reagent Kits v1.1 with Feature Barcoding technology (10x Genomics, CG00208) according to manufacturer’s instructions. Each sample was loaded in duplicate for a total target recovery of 5,000 cells (20,000 for sample pools).

###### After GEM dissolution and Dynabeads purification, 15 PCR cycles were done using the SC5ʹ Feature cDNA Primers (PN-1000080) to amplify the DNA from cell surface protein Feature Barcode oligos together with the full-length cDNA. The two products were separated by size selection and used for generating the different types of libraries. To construct the GEX library, the amplified full-length cDNA was fragmented, end repaired, A-tailed, and sample indexed using the Chromium Single Cell 5’ Library Construction Kit (10x Genomics, 1000020). For the V(D)J library, human T and B cell V(D)J sequences were enriched from the amplified cDNA with the Chromium Single Cell V(D)J Enrichment Kits (PN-1000005 and PN-1000016 for T and B cells respectively) followed by fragmentation, end repairing, A-tailing and sample indexing. Finally, the CSP library was generated from the amplified DNA from cell surface protein Feature Barcode by one-step PCR amplification using the Chromium Single Cell 5' Feature Barcode Library Kit (PN-1000080). Quantification and fragment size distribution of cDNAs and final libraries were determined using the Agilent 2100 BioAnalyzer High Sensitivity DNA kit (Agilent Technologies). All constructs were sequenced together on a Novaseq 6000 (Illumina), targeting a median sequencing depth of 20,000 (GEX), 2000 (VDJ) and 8,000 (CSP) reads per cell.

#### Spatial Transcriptomics (Visium OCT)

###### Spatial visualization of gene expression within tonsil tissue was conducted using the Visium Spatial Gene Expression kit (10x Genomics) as per manufacturer's protocol. The OCT blocks were cut twice using a cryostat (Leica CM1950): a first time to assess RNA quality and assure a minimum RNA Integrity Number (RIN) number of 7 (RNA pico Chip) and a second time to mount a 10 μm section on the Visium slides. Slides were H&E stained before the sections were imaged using the NanoZoomer S60 (Hamamatsu) to assess tissue morphology and quality. The sections were then permeabilized for 6 min, according to the results of a corresponding Tissue Optimization experiment (10x Genomics, CG000238), and processed according to the Visium Spatial Gene Expression user guide (10x Genomics, CG000239). In short, tissue was lysed and reverse transcription was performed followed by second strand synthesis and cDNA denaturation. Spatially barcoded, full length cDNAs were amplified by PCR for 16 or 18 cycles, depending on the initial concentration previously determined by qPCR. Indexed sequencing libraries were generated via end repair, A-tailing, adaptor ligation and sample index PCR and analyzed using the Agilent 2100 BioAnalyzer. Libraries were sequenced on an Illumina NovaSeq 6000 with sequencing depth of ~100,000 reads per spot.

### QUANTIFICATION AND STATISTICAL ANALYSIS

#### scRNA-seq: Data alignment

We used cellranger count (v4.0.0, 10x Genomics) to align reads to the GRCh38 human genome, with the “chemistry” parameter set to “SC3Pv3”. As cell-hashed and non-cell-hashed libraries had different target recoveries, we set the “expect-cells” parameter to 20,000 and 5,000, respectively. For cell-hashed samples, the “libraries” and “feature-ref” parameters were specified as described in the “Feature Barcode Analysis” pipeline of Cell Ranger (<https://support.10xgenomics.com/single-cell-gene-expression/software/pipelines/latest/using/feature-bc-analysis>). HTO sequences for each library can be found at Tables S20.

#### scRNA-seq: Demultiplexing of HTO

We performed all downstream pre-processing with Seurat (v3.2.0 and v4.1.0). To normalize HTO counts, we applied a centered-log-ratio transformation across HTO, as implemented in the function “NormalizeData” (normalization.method = “CLR”, margin = 1). To assign an HTO to each cell, we used the “HTODemux” function (positive.quantile = 0.99). Briefly, this function performs k-medoid clustering (k = # HTO + 1) and uses the cluster with the lowest average to find the “negative” distribution for each HTO. Then, it fits a negative binomial distribution and uses the 0.99 quantile as threshold, which classifies cells as positive or negative for each HTO. We excluded cell barcodes not assigned to any HTO (“Negative”), as they had a lower library size and low number of detected genes. On the other hand, we kept cell barcodes assigned to two or more HTO to increase the statistical power and robustness of our doublet detection strategy (see below). To compare hashing efficiency across libraries, we computed a signal-to-noise ratio (SNR) for each cell as follows:

SNR =$\frac{CLR-normalized counts (HTO1) + 0.1}{CLR-normalized counts (HTO2) + 0.1}$

Where HTO1 and HTO2 are the HTO with the first and second largest counts for that cell, respectively.

#### scRNA-seq: Filtering and data normalization

We noticed that the library size distribution (total unique molecular identifiers; UMI) was higher in cell-hashed libraries than in non-cell-hashed samples. Following the current best practices (Luecken and Theis, 2019), we determined the quality control (QC) thresholds for cell-hashed and non-cell-hashed libraries separately. Not to bias cell type composition, we decided to be as permissive as possible and applied more stringent thresholds at the cluster level. For non-cell-hashed libraries, we excluded cell barcodes with <1,000 UMI, <250 detected genes or a mitochondrial expression >20% (potential lysed cells or empty droplets). For cell-hashed libraries, we filtered out cell barcodes with <1,000 UMI, <400 detected genes or a mitochondrial expression >20%. In addition, we excluded genes detected in <=5 cells. A full discussion on why we chose these thresholds can be found at the associated reports available on GitHub. To adjust for differences in total UMI across cells, we used the function NormalizeData (normalization.method = "LogNormalize", scale.factor = 1e4). This function divides the raw gene counts for each cell by the total counts of that cell and multiplies it by the scale factor (10,000), which is then log-normalized as log(1+x).

#### scRNA-seq: Feature selection, dimensionality reduction and batch effect correction

Before clustering cells to define tonsillar cell types and states, we performed three important steps:

(1) calculate the proportion of doublet nearest neighbor for each cell (pDNN, described below);

(2) merge and integrate our dataset with the Seurat object from King et al.(King et al., 2021) , and

(3) merge and integrate the resulting Seurat object with the RNA slot of our Multiome experiments.

The latter two allowed us to reach a robust consensus annotation across studies, to include an external control to ensure we preserved biological variability, and to connect chromatin accessibility with gene expression. Because we included new sets of cells in these steps, we reasoned that the set of highly variable genes (HVG) and axis of variability would change. Thus, for each step we executed the following steps:

1. We used the function FindVariableFeatures of Seurat (with default parameters) to extract the top 3,000 HVG for steps 1 (doublet detection) and 2 (integration with King *et al.*(King et al., 2021). For step 3, we noticed that the three expression matrices (scRNA-seq, King et al., Multiome) consisted of a different set of genes, consistent with the poor mixability between single-cell and single-nuclei RNA-seq techniques (Mereu et al., 2020). To homogenize it, we found the top 5,000 dataset-specific HVG and took the intersection (1,740 genes), which we used as input for the next two functions.
2. We performed a z-score transformation (Seurat: ScaleData) on the normalized-values, followed by principal component analysis (Seurat: RunPCA).
3. To correct for batch effects, we used Harmony v1.0 (Korsunsky et al., 2019), as a recent benchmarking effort reported that it scales well to hundreds of thousands of cells and it is amongst the best integration tools (Tran et al., 2020). We used the function RunHarmony, with the top 30 principal components (PC) as input. We considered cells coming from different GEM wells (see above) as different batches (specified in the group.by.vars parameter).
4. We then assessed the success of the data integration qualitatively with UMAP (Seurat: RunUMAP, first 30 PC) and quantitatively with the Local Inverse Simpson’s Index (lisi v1.0: compute_lisi). This LISI score ranges from 1 to N (number of batches), and quantifies the diversity of batch labels on the neighborhood of each cell. Thus, the larger the LISI, the better the batch integration. We applied both approaches to measure the effect of six potential confounders before and after integration: library, sex, age group, hashing status, sampling center and assay (3’, 5’ or Multiome). To ensure we preserved biological heterogeneity, we plotted the cell type labels provided by King et al. on the aforementioned UMAP.

#### scRNA-seq: Doublet detection and removal

Although we considered cell hashing as our gold-standard method for doublet detection, it presents some limitations. First, cell hashing cannot detect intra-index doublets (doublets with the same hashtag/index). Second, hashing efficiency differencing across libraries can introduce detection variability (measured by signal-to-noise ratio). Finally, non-hashed libraries will have a doublet rate at approximately 4% per sample. To mitigate these issues, we ran Scrublet v0.2.1 (Wolock et al., 2019), which simulates and predicts doublets computationally. Following the best practices, we executed scrublet for each library separately. Since we expect a doublet rate of 4% for a target recovery (TR) of 5,000 cells, we set the “expected_doublet_rate” of the “Scrublet” function to 0.04 and 0.16 for non-hashed (TR=5,000) and hashed libraries (TR=20,000), respectively. For the scrub_doublets function, we set the parameters min_counts to 2, min_cells to 3, min_gene_variability_pctl to 75 and n_prin_comps to 50. The resulting doublet scores (which range from 0 to 1) and predictions (True or False) were added to the metadata of the Seurat object.

Notably, both cell hashing and scrublet were run for each library independently. Thus, we aimed to combine both approaches in a single metric that can consider all cells in the dataset, hence increasing the statistical power to detect doublets. To this end, we computed the pDNN, a metric inspired by the proportion of artificial nearest neighbors (pANN) introduced by DoubletFinder (McGinnis et al., 2019). Briefly, we used the top 30 harmony-corrected PCs to find the 75-nearest neighbors for each cell (Seurat: FindNeighbors). Then, we calculated a pDNN for each cell by dividing the number of nearest neighbors labeled as doublets by the neighborhood size (75). We followed this approach for three doublet annotations: cell hashing, scrublet and their union. Finally, we observed that the regions of the UMAP with the highest pDNN values corresponded to cell neighborhoods that expressed two or more markers of different lineages (CD3D and CD79B, for example); which validated its use. Overall, we excluded 80,577 doublets labeled by cell hashing; and flagged the ones detected by scrublet. In addition, we kept the pDNN scores in the metadata to filter out clusters of doublets downstream.

#### scRNA-seq: Clustering and annotation

To cluster cells into cell types and states, we followed a top-down, recursive approach, organized from general to specific. This approach is inspired by the mouse brain atlas (Zeisel et al., 2018). At each level, we performed Louvain clustering by first calculating a K-nearest neighbors graph (Seurat: FindNeighbors, reduction = “harmony”, top 30 PC), and then determining the exact clusters (Seurat: FindClusters). The “resolution” parameter of FindClusters is what drives the number of clusters, and was adjusted differently at each level. At each level, we subsetted one or more clusters, thus increasing our ability to detect finer-grained heterogeneity. Therefore, at each level we have rerun steps 1-3 described above to find HVG, perform PCA and correct for batch effects. Every level was an opportunity to fetch and discard clusters of poor-quality cells and doublets using the different sources of evidence we gathered in the analysis explained above. Below a more detailed explanation of each cluster level:

- Level 1: we reasoned that, if clusters represent stable categories, we should be able to classify unseen transcriptomes to those categories with high accuracy. Thus, we fitted a random forest classifier (randomForest package) using the top 30 harmony-corrected PC as features to predict clusters derived from varying clustering resolutions. We plotted the resulting out-of-bag accuracies as a function of the clustering resolution, and determined an “elbow” in the plot to find the optimal resolution (0.25), which resulted in 12 clusters. We performed a “one-vs-all” differential expression analysis to find markers specific to each cluster (Seurat: FindMarkers, test.use = “wilcox”). After interpreting the top markers per cluster, we split and merged them into a biological sound manner. For example, the epithelial cells clustered together with the myeloid; and the precursor T and B cells clustered with the PDC. Moreover, we removed one cluster that showed hallmarks of poor-quality cells. Overall, we identified nine major cell compartments, which correspond to the categories in (Figure S1D).
- Level 2: we split the main Seurat object into nine (one per major compartment), and rerun the data integration pipeline. Moving forward, we considered “assay” (scRNA-seq or Multiome) as the batch variable to correct for, as it was the main driver of variance. In this step, we had high statistical power to remove clusters of poor-quality cells and doublets to clean the dataset.
- Level 3: for cell compartments with fewer cells (myeloid, FDC, PDC and epithelial), we excluded both cells from King *et al.* (King et al., 2021) dataset and from Multiome; because we reasoned that our assay-specific feature selection and integration strategy would overcorrect biological heterogeneity in these underrepresented compartments. In addition, since epithelial cells were composed of fewer than 1,000 cells, we used only the top 20 PCs and reduced the neighborhood size (k) from 20 to 10.
- Levels 4-5: we leverage the annotation from King et al. (King et al., 2021) as a starting annotation, and hereafter focused solely on our own data. Because biology-driven clustering led to more meaningful cluster labels, we discarded the random forest strategy. We found markers for each cluster with FindMarkers, as explained above. We developed a Shiny app that allowed annotation experts to explore the expression of marker genes (<https://singlecellgenomics-cnag-crg.shinyapps.io/Annotation/>) and to determine a biologically sound clustering. Further, we used the FindSubCluster function from Seurat to stratify heterogeneous clusters. Of note, we moved naive CD8 T cells from the CD4 T cell compartment to Cytotoxic cells.

#### scRNA-seq: Gene signature scoring

To collapse the expression of a set of genes into a per cell gene signature, we used the AddModuleScore function from Seurat. Later, we additionally used the UCell package (Andreatta and Carmona, 2021), which was shown to outperform AddModuleScore. Finally, we used the CellCycleScoring function and the predefined list of cell cycle markers from Seurat to calculate a per cell S.Score and G2M Score, as well as to classify cells into cell cycle phases. For the obtention of endoplasmic reticulum (ER) signature in PC we first performed a differential expression analysis (DEA) between the LZ-GCBC and short-lived IgM+ PC and we then used the Database for Annotation, Visualization and Integrated Discovery (DAVID) (Huang et al., 2009; Sherman et al., 2022) to perform a KEGG pathway enrichment analysis using the upregulated genes.

#### scRNA-seq: Gene Set Enrichment Analysis

To find specific functions associated with each slancyte subset, we conducted a one-vs-all differential expression analysis for each slancyte subset (Seurat: FindMarkers, only.pos = FALSE, logfc.threshold = 0). We arranged the resulting gene list by decreasing log2 fold-change. We performed a gene set enrichment analysis (GSEA) for each gene list using the function gseGO (ont = "BP", OrgDb = org.Hs.eg.db, keyType = "SYMBOL", minGSSize = 10, maxGSSize = 250) from clusterProfiler v4.3.4 (Wu et al., 2021). We filtered out gene ontology (GO) terms with an adjusted p-value > 0.05, and arranged them by decreasing normalized enrichment score (NES). Finally, we plotted the running enrichment scores with the gseaplot function for selected GO terms.

#### scRNA-seq: Validation with external datasets

To validate the upregulation of SIX5 in plasma cells using external datasets, we queried the human cell atlas Bone Marrow Viewer, available at <http://www.altanalyze.org/ICGS/HCA/splash.php> (Hay et al., 2018) and the “Blood (PBMC) Hao” single cell RNA-seq track (Hao et al., 2021) at the UCSC genome browser (<https://genome-euro.ucsc.edu/cgi-bin/hgGateway>). In addition, we downloaded ChIP-seq data of the histone mark H3K27ac, which was generated as described in (<http://www.blueprint-epigenome.eu/index.cfm?p=7BF8A4B6-F4FE-861A-2AD57A08D63D0B58>) (Beekman et al., 2018; Ordoñez et al., 2020). The normalized signal from this histone mark was captured for SIX5 and its target genes, except for TSC22D3 gene (located at chromosome X), in NBCT, tonsillar NBC (n=3); NBCB, NBC from peripheral blood (n=3); GCBC (n=3), csMBC, class-switch MBC (n=2); ncsMBC, non-class switch MBC (n=1); PC (n=3); MM (n=4). To identify slancyte subpopulation, we downloaded the differentially expressed genes between tonsillar slan+ cells, CD11b+CD14+-macrophages, and CD1c+DCs/cDC2 from a recent study (Bianchetto-Aguilera et al., 2020).

#### Gene regulatory network inference

To infer transcription factor (TF) activity, we used pySCENIC v0.10.3 (Van de Sande et al., 2020) on the scRNA-seq raw matrices from T cells and B cells separately. Briefly, pySCENIC infers co-expression modules (known as regulons), composed of a given TF and its putative target genes, and measures their activity in each individual cell. GRNBoost2 algorithm from the Arboreto package (Moerman et al., 2019) was used to infer the co-expressed modules from a predefined curated list of 1,797 human TFs, provided in the pySCENIC repository (https://github.com/aertslab/pySCENIC). Next, cisTarget (Herrmann et al., 2012) was applied to refine regulons by pruning indirect targets based on cis-regulatory motifs footprints using human motifs v9 and hg38 (500bp upstream of TSS, and 100 bp downstream) from cisTarget databases (https://resources.aertslab.org/cistarget/). This process gave a total of 189 and 214 regulons for T cells and B cells respectively. Finally, the regulon activity per cell was quantified using an enrichment score for the targets of each regulon (AUCell). Additionally, regulon specificity score (RSS) for the B cell lineage was computed using the Jensen-Shannon Divergence (Suo et al., 2018).

#### Cell-to-cell interactions

To analyze potential cell-to-cell interactions, we used CellPhoneDBv3 (Garcia-Alonso et al., 2021) that allowed us to identify significant ligand-receptor pairs between myeloid and T cell populations. First, we subset the scRNA-seq dataset to all myeloid populations and naive CD4 T, naive CD8 T, Eff-Treg, Eff-Treg-IL32 and Tfr cell types. We then performed differential gene expression analysis of each cell type versus the rest of the subset populations using a two-sided Wilcoxon rank-sum test in Seurat v4.0.0 (FindMarkers) (Hao et al., 2021). All genes expressed in >10% of the cells of the target population and a Bonferroni-adjusted p-value < 0.05 were selected for CellPhoneDB analysis. CellPhoneDB method degs_analysis selected those interactions where all genes were expressed in >10% of the cells and at least one of the genes was a differentially expressed gene. The output file was manually curated to select those interactions involving slancytes that best featured their potential function in T cell modulation.

#### scATAC-seq: Data alignment

We used Cell Ranger ATAC v1.2.0 to map the fastq files using GRCh38-1.2.0 as the human reference genome. Specifically, we run cellranger-atac count command on each individual library to perform read filtering and alignment, barcode correction and counting, and peak calling. Then, to pool the samples together, we used cellranger-atac aggr command with a new peak calling round. To maximize the sensitivity of the input libraries, we set the normalization model to “None”.

#### scATAC-seq: Data quality control

We performed all downstream analysis with Seurat v3.9.9 and its extension package Signac v1.1.0 (Stuart et al., 2021). The total number of non-filtered aggregated cells were 64,162 with 5,724 median fragments per cell, 71,3% fraction of fragments overlapping any targeted region and a 52.7% fraction of transposition events in peaks in cell barcodes. We determined the QC thresholds for each library individually by applying non-restrictive thresholds. At this point, we noticed that the fraction of fragments falling within the peaks was significantly higher in FACS-processed libraries than in non-FACS libraries. To remove low-quality cells from the aggregated libraries, we excluded cells that presented: (1) a total number of fragments in peaks fewer than 700 or greater than 30,000; (2) had a fraction of fragment in peaks fewer than 15; (3) a transcriptional start site enrichment score fewer than 2; (4) a ratio of reads assigned to blacklist regions greater than 0.03; (5) and cells with peak counts fewer than 500 or higher than 100,000; (6) peaks detected in 5 o fewer cells. After this filtering step, we end up with 58,049 high-quality cells. A full discussion on the reasoning behind these thresholds can be found at the associated reports available on GitHub in the following link: https://github.com/Single-Cell-Genomics-Group-CNAG-CRG/TonsilAtlas/tree/main/scATAC-seq/2-QC

#### scATAC-seq: Data normalization and Integration

We merged and integrated the scATAC-seq dataset with the ATAC slot extracted from the Multiome experiments (described below). This approach allowed us to (1) increase the number of scATAC-seq cells, (2) reach a consensus chromatin profile not biased by the techniques, (3) create a direct link between ATAC peaks and gene expression to avoid inferring the activity of each genes from chromatin accessibility, and (4) transfer both the cluster labels and UMAP coordinates defined with gene expression to the scATAC-seq dataset (see below). When merging multiple single-cell chromatin datasets, it is essential to note that the peak calling was done in each dataset independently. For this reason, we first created a consensus set of peaks across all datasets using the *UnifiedPeaks* function (mode = "reduce"). We then filtered out the peaks based on length; peaks with a width greater than 10,000 bp or less than 20 bp were removed. We quantified the consensus peaks in each dataset using the *FeatureMatrix* function, and then used these unified matrices to create new Seurat objects that were finally merged using the *merge* function. This resulted in 101,279 cells and 166,156 features. For each dataset individually and then for the merged dataset (scATAC-seq and Multiome), we ran the following pipeline:

1. We applied the term frequency-inverse document frequency (TF-IDF) normalization (Signac: *RunTFIDF*, method =1, with the default parameters). TF-IDF corrects for differences in library size across cells, and penalizes peaks that are homogeneously open or closed across all cells.
2. We set the *FindTopFeatures* function to q0 to consider all the features for the dimensional reduction step. To obtain a reduced dimension representation of the dataset, we ran the singular value decomposition (SVD) on the TF-IDF data normalized. Because the variance captured by the first LSI component was explained by library size, we decided to exclude it for downstream analysis.
3. We used Harmony v1.0 (Korsunsky et al., 2019), to correct for batch effects because it is amongst the best-performing integration tools for scATAC-seq data (Luecken et al., 2022). Specifically, we executed the *RunHarmony* function with default parameters selecting from the second to the n first LSI components (n is variable depending on the dataset analyzed) and group.by.var equal to "gem_id" (GEM well) or "assay".
4. To assess Harmony’s performance, we used the LISI score (Korsunsky et al., 2019) to verify the quality of the data integration across 5 main categorical confounders: sex, age group, sampling center, library and assay or tecnhique applied in the dataset.

#### scATAC-seq: Doublet detection

To predict the potential doublets from single-cell ATAC-seq data, we accumulated different sources of information:

1. We used a modified version of Scrublet v0.2.1 (Wolock et al., 2019) to compute the cell doublet score per library following these parameters: log_transform=True, min_counts=2, min_cells=3, min_gene_variability_pctl=70, n_prin_comps=50. Note that for the BCLL-14-T and BCLL-15-T samples, the expected doublet rate was set to 0.056 (TR=7,000 nuclei) compared to the rest that was set at 0.04 (TR=5,000 nuclei). We flagged as True the predicted doublets defined by the automatic threshold provided by Scrublet and this information was added to the Seurat object metadata.
2. To discard doublets, nuclei clumps and other artifacts, we removed cells with extremely high numbers of fragments in peaks.

#### Multiome: Data alignment

We used Cell Ranger v1.0 to map the fastq files to the GRCh38-2020-A as a human reference genome. Specifically, we run cellranger-arc count on each individual library to perform read filtering and alignment, barcode correction and counting, peak calling and counting of both ATAC and GEX molecules. A detailed description of all the quality control parameters evaluated can be found at the associated reports available on GitHub.

#### Multiome: Data quality control

The downstream analysis was done in R applying Seurat v3.9.9 and its extension package Signac v1.1.0. The total number of non-filtered cells was 77,006. To remove low-quality cells from each library, we applied the following filters: (i) For scRNA-seq, we excluded cell barcodes with fewer than 550 UMI, fewer than 250 detected genes or a mitochondrial expression higher than 20%, (ii) For scATAC-seq, we excluded cells that presented a total number of transposition events less than 500 or greater than 100,000, that had a transcriptional start site enrichment score less than 2 and a nucleosome signal less 2, which resulted in 69,118 filtered cells. Note that the BCLL-2 sample was excluded from the scATAC-seq analysis because it presented a low overall quality that could lead to a misinterpretation of the data. After peak calling individually for each library, we merged the libraries using the *UnifyPeaks* function (with default parameters), which aligns the ranges of peaks and merges the overlapping ones to produce a simplified set of intersecting peaks. We filtered out peaks with a width size less than 20 bp or greater than 10,000 bp. We quantified the accessibility counts in the new set of peaks using the *FeatureMatrix* function. Since the peaks were common across libraries, we merged all cells from all libraries into a single accessibility matrix.

#### Multiome: Doublet detection

To predict the potential doublets from single-cell ATAC-seq data, we accumulated different sources of information:

1. We used a modified version of Scrublet v0.2.1 (Wolock et al., 2019) to compute the cell doublet score per library following these parameters: log_transform=True, min_counts=2, min_cells=3, min_gene_variability_pctl=70, n_prin_comps=50. Note that for the BCLL-14-T and BCLL-15-T samples, the expected doublet rate was set to 0.056 (TR=7,000 nuclei) . We flagged as True the predicted doublets defined by the automatic threshold provided by Scrublet and this information was added to the Seurat object metadata.
2. To discard doublets, nuclei clumps and other artifacts, we removed cells with extremely high numbers of fragments in peaks.

#### Multiome: Data normalization and Integration

The data was treated as individual scRNA-seq and scATAC-seq objects and we repeated the standard downstream analysis explained above including data normalization, variable gene detection, data scaling, dimensionality reduction analysis, batch correction with Harmony and UMAP representation. To identify the nearest neighbors for each cell based on the weighted combination of the scRNA-seq and scATAC-seq modalities, we applied *FindMultiModalNeighbors* function to construct a weighted nearest neighbor (WNN) graph. The harmony integration of each modality was used as a dimensionality representation of each object using the first 30 PCs for scRNA-seq and the first 40 LSI components for scATAC-seq.

#### Alignment of scATAC-seq with Multiome datasets.

To help the interpretation of the scATAC-seq integrated dataset, we classify the Multiome ATAC-seq cells using the annotation previously defined by the scRNA-seq from the same experiment, since the cells share the same cellular barcode. To extend the annotation to the rest of the scATAC-seq cells, we applied a k-nearest neighbour (KNN) algorithm to classify those cells to a given cell type category with the help of our Multiome training set. Note that KNN works on a basic assumption that data points of similar categories are closer to each other. To cross-validate the number of nearest neighbours to consider (the K parameter), we split our training set in two parts: a train.loan, that corresponds to the random selection of the 70% of the training set and the test.loan, that is the remaining 30% of the data set. The first one was used to train the system while the second was used to evaluate the learned system. We built the machine learning model using the optimal k. Note that the probability of the prediction was lower in the transitioning cells and in not-defined clusters.

#### Peak calling based on annotation levels

To identify more precise and specific peaks on the annotated cell types, we decided to do multiple rounds of peak calling using MACS2 v2.2.7.1 (Zhang et al., 2008) as we increase the level of the clustering resolution. Specifically, we used the *CallPeaks* function provided by Signac setting the group.by parameter by the corresponding annotation level, removing peaks on non-standard chromosomes and on genomic blacklist regions. We quantified the new peak counts in the specific dataset by generating a consensus peak set and repeated the standard downstream analysis explained above; including data normalization, dimensionality reduction analysis, batch correction with Harmony and UMAP representation.

#### scATAC-seq specific chromatin features

To find differentially accessible features between the clusters defined at level 1, we performed a differentially accessibility (DA) test between all of them. We use a logistic regression (Ntranos et al., 2019) adding the nCounts_peaks as a latent variable to mitigate the effect of sequencing depth (Signac: FindAllMarkers). Next, we filtered out the DA peaks with a Bonferroni-adjusted p-value less than 0.5 and selected the top 2,000 DA peaks to create the list of features needed to calculate the chromatin signature. To do that, we compute the ChromatinVar deviation for each cell type applying the *ChromatinModule* function. A more restrictive analysis was done to select specific DARs in the context of GCBC cells. In this case, a score was computed to identify regions with high accessibility of one cell type in relation to the others. Specifically, for each region, we computed the ratio between the maximum accessibility value across cell types to the sum of all accessibility values for that region. Then, the top scoring DARs that belonged to clusters of interest were selected.

#### Motif analysis

To find the consensus binding motifs in the DNA sequences, we used the chromVar v1.1.0 R package (Schep et al., 2017), which calculates for each motif annotation and each cell, a bias-corrected “deviation” in accessibility from an expected value based on the average of all the cells. This allowed us to visualize motif activities per cell in each of the clusters. Note that the motif annotation was performed using two databases to compare the robustness of the results. Specifically, we downloaded 746 transcription factor motifs from the JASPAR vertebrates core (using JASPAR2020 v0.99.10 R package) (Fornes et al., 2020; Tan and Lenhard, 2016), and 1,764 from CisBP database using human_pwms_v1, the curated collection of human motifs (as included in the R package chromVARmotifs v0.2.0 (Schep et al., 2017). Based on the underlying question, we performed two different type of analysis: (1) identification of overrepresented motifs in a set on genomic regions using *FindMotifs* function provided by Signac or (2) differential testing on the chromVAR z-score using the *FindMarkers* function between the clusters to compare.

#### Estimating co-accessible sites

To enhance the interpretation of the scATAC-seq data, we decided to use the Cicero v1.3.4 R package (Pliner et al., 2018) that allows: (1) estimating the co-accessible sites in the genome, and (2) predicting potential cis-interacions between proximal/distal regulatory elements and their putative target genes. Specifically, we first converted the CD4 T Seurat object to CellDataSet format and to the Cicero object, using the Signac-provided functions: *as.cell_data_set and make_cicero_cds* respectively. Note that for the second function, we specify the coordinates in low-dimensional Harmony space. Then, we executed the wrapper function called *run_cicero* (with the default parameters) to get the pairwise co-accessibility scores for all the peaks identified in CD4 T cells.

#### CITE-seq: Data alignment

We used Cell Ranger v6.0.1 multi to align simultaneously 5’ scRNA-seq, antibody profiles and TCR/BCR-seq, enabling consistent cell calling between the library types. Specifically, the Cell Ranger uses the fastq files from all four modalities and performs alignment to the GRCh38-2020-A, filtration, feature barcode and UMI counting for both genes and antibody tags; along with the VDJ sequence assembly and clonotype counting (GRCh38-alts-ensembl-5.0.0 as the human genome reference).

#### CITE-seq: Genotype demultiplexing

For BCLLATLAS_38, we pooled donors into a single experiment. Genotypes were subsequently demultiplexed using Vireo v0.5.0 (Huang et al., 2019) based on individual genotypes inferred from scRNA-seq read information. First, we used cellSNP (Huang and Huang, 2021) to pileup the mapped reads at each single nucleotide variant (SNP), filtering variants with less than 20 UMIs and a minor allele frequency of less than 10% in the compiled list of 7.4 million common variants (AF>5%) present in 1000 Genome Project and gnomAD. Then, the pileup allelic profile of each cell barcode was used for donor deconvolution and doublet detection using Vireo. Out of 6,679 multiplexed cells, 380 were detected as doublets and 301 were unassigned cells.

#### CITE-seq: Quality control

We performed the downstream analysis using Seurat v4.0. Specifically, the quality control was performed in two main stages. Firstly, cells assigned as doublets by Scrublet (Wolock et al., 2019) and genotype doublets and unassigned donor cells by Vireo were eliminated. Following the current best practices (Luecken and Theis, 2019), we performed QC on each CITE-seq experiment separately. Cells outside of the threshold range of mitochondrial content, UMI counts and feature count set per subproject were filtered. After this first filtering step, we obtained a total of 42,929 cells, of which 12,867 had BCR (B cell repertoire) and 7,795 TCR (T cell repertoire) information. The QC plots and exact filtering thresholds can be found at the GitHub repository (see code availability section). Next, a top-down approach was used for a second quality control. We zoomed into T and B cell clusters to refine the quality based on marker expression. Specifically, we removed B cells that exhibited a high expression of T cell marker genes (such as CD3, CD4 and CD8) and T cells with high expression of B cell markers (such as CD19, CD5, CD27) as well as cells with dual repertoire (both TCR and BCR) as potential doublets. 40,396 high-quality cells entered the subsequent analysis.

#### CITE-seq: Data normalization and Integration

We performed data normalization and integrations as follows:

- Normalize and correct for batch effects: the gene expression matrix was normalized as described before for scRNA-seq. For ADT data, we applied a centered log ratio (CLR) across cells (Seurat: NormalizeData, margin=2) and corrected for batch effects using Harmony (Korsunsky et al., 2019).
- Weighted-nearest neighbor (WNN) graph-based integration: The normalized and homogenized matrices were used for the construction of WNN graphs based on cell-specific data modality weights. This enables dimensionality reduction based on the weight of both modalities (Seurat: FindMultiModalNeighbors). We used the first 30 and 20 harmony-corrected principal components for RNA and ADT, respectively.
- Label Transfer: SLOcator was used to transfer labels and coordinates defined with scRNA-seq to CITE-seq (see below).

#### CITE-seq: Repertoire analysis

We used Scirpy v0.7.0 (Sturm et al., 2020) to analyze TCR and BCR repertoires. Each repertoire sequence is formed by V, D and J gene recombination. Structurally, each of the repertoire representations consists of the framework region (FWR) and complementarity determining regions (CDR), which primarily interacts with the epitope. Clonotypes can be defined by using different algorithms (such as identity, blosum matrix based similarity, hamming distance and levenshtein distance) and sequence levels (nucleotide and amino acid). Here, we defined clonotypes based on the CDR3 nucleotide sequence identity and V gene usage (Scirpy: define_clonotypes, default parameters) within samples. For TCR and BCR analysis, we defined clonotypes expanded if three or more cells showed the same sequence (Scirpy: clonal_expansion).

#### ST: Data processing

We used spaceranger count v1.1.0 to align the fastq files to the GRCh38-2020-A as a human reference genome. For each tissue slice we ran spaceranger count with its specific Visium slide ID, its capture area and the specific image in .jpeg format, these can be found in the project's github (see code availability section). Visium slide-specific spot layouts were used for each slide to determine the spatial coordinates of each spot.

#### ST: Quality control

Downstream analysis was done in R 4.0.1 and Seurat v4.1.0. We first assessed the distribution of library size, detected genes, and mitochondrial and ribosomal percentage across the slide to assess for overpermeabilization and subsequent lateral diffusion of reads. We noticed that the distribution of the library size highly correlated with histological features. Therefore, we decided to keep all spots overlaying the tissue. In the slide from sample BCLL-8-T there was a region that had been folded onto itself. Spots overlapping this folded region were removed since they showed lower library size and number of detected genes as well as a transcriptomic profile that reflected a mixture of regions. We determined mitochondrial and ribosomal percentage by dividing the number of UMIs assigned to genes starting with MT- or RPL|RPS respectively over the total library size for each spot. A detailed description of all the quality control parameters evaluated and environment used can be found at the associated reports available on GitHub (see code availability section).

#### ST: Data normalization

To adjust for differences in total UMI across spots, we used the function *NormalizeData* (normalization.method = "LogNormalize", scale.factor = 1e4). This function divides the raw gene counts for each cell by the total counts of that cell and multiplies it by the scale factor (10,000), which is then log-normalized as log(1+x).

#### ST: Feature selection, dimensionality reduction and batch effect correction

We first used the function *FindVariableFeatures* of Seurat (with default parameters) to extract the top 3,000 HVG. We compared this gene set with the ones obtained from algorithms aiming to detect spatially variable genes such as *FindSpatiallyVariableFeatures*, *spatialDE* and *SPARK* and found considerable overlap. We then proceeded to the downstream analysis with the HVG obtained with *FindVariableFeatures* due to the lower computational time required. Next, we performed a z-score transformation (as implemented in the function *ScaleData*) on the normalized-values. We then carried out PCA dimensionality reduction with the function *RunPCA*. At this point batch effects were observed in the PCA space, therefore, we corrected for batch effects using Harmony v1.0 (Korsunsky et al., 2019), as a recent benchmarking effort reported it to be amongst the best three performing integration tools (Korsunsky et al., 2019). We used the function *RunHarmony* with the top 20 principal components (PC) as input, the top 20 was decided after looking at the PCA elbow plot. We considered each tissue slice as different batches as each one was processed in a different capture area. Full analysis of the integration and batch correction can be found in the GitHub repository.

#### ST: Tissue region clustering and annotation

To annotate our tissue slices we followed two approaches. In the first approach, we carried out an unbiased data-driven approach in which we aimed to cluster the spots and annotate them using differentially expressed genes. In the second approach, expert pathologists manually annotated each tissue slide. Spot clustering was performed by using the functions *FindNeighbors*, which computes a shared nearest neighbor graph on the harmony integrated embedding; first 20 Harmony components were used, for all the spots. We then identified clusters of spots by using shared nearest neighbor (SNN) modularity optimization based Louvain clustering algorithm using *FindClusters* function. We computed the clustering with varying degrees of resolution to assess which one fit best our datasets. Annotation of tissue regions was carried out using resolution 0.3. We used the function *FindAllMarkers* to identify differentially expressed genes between clusters using the Wilcoxon rank sum test on the log-normalized gene expression. Prior knowledge marker genes were used to determine the identity of each cluster. Manual annotation by pathologists was carried out using a custom built shinyapp (<https://singlecellgenomics-cnag-crg.shinyapps.io/Annotation/>) that allowed to select spots on the tissue and download the spot barcode for each selection.

#### ST: Cell type deconvolution

Integration of spatial transcriptomics with the reference scRNA-seq to obtain cell type deconvolution was performed using SPOTlight v0.1.7 (Elosua-Bayes et al., 2021). Different SPOTlight runs were carried out for the different populations of interest as described below.

To deconvolute major cell types we used the annotation column *annotation_figure_1*. From this annotation we consolidated all the cycling cell types into one, *Cycling*, so the cycling signature didn’t drive the deconvolution of individual cell types. We then computed differential expressed genes between all populations Seurat’s function *FindAllMarkers* with the default Wilcoxon rank sum test and up to 500 cells per cell type. Next, we randomly sampled up to 50 cells per cell type from as few batches as possible in order to reduce the batch effect. Lastly, we ran deconvolution using the previously selected cells and the gene set resulting from union between the 3,000 most highly variable genes and the differentially expressed genes. Cell types predicted to contribute <3% of a spot were considered to be 0.

For CD4 T cell specific deconvolution, we used a combination of the annotation column *annotation_20220215* and *annotation_level_1*. This combination allowed us to have finer grained annotation for the CD4 T subpopulations, while maintaining the level-1 annotation for the other major cell types. This ensured that we captured the signal provided by the main cell types, while taking into account CD4 T heterogeneity. From the finer grained annotation, we consolidated CM Pre-non-Tfh, CM PreTfh into one cell type and labeled them as CM PreTfh/Pre-non-Tfh, since they presented very similar phenotypes. We excluded preBC, preTC as they represent very few cells overall and introduced undesired noise to the model. We also excluded those cells annotated as CD4 T cells in level-1 but not annotated as a CD4 subpopulation at the more granular level. We then carried out two rounds of differential expression. The first was carried out among all the cell types using the annotation specified in *annotation_level_1*. This allowed us to capture differentially expressed genes between major populations. As before, we used Seurat’s function *FindAllMarkers* with the default Wilcoxon rank sum test and up to 200 cells per cell type. The second round of differential expression was computed only between the CD4 T subtypes to capture genes more subtly differentially expressed between them. Both lists of differentially expressed genes were filtered by logFC, pct.1 and p value to keep only relevant genes. We then randomly sampled up to 100 cells per cell type from as few batches as possible to reduce the batch effect. Lastly we ran deconvolution using the previously selected cells and the gene set resulting from union between the 3,000 most highly variable genes and the differentially expressed genes. Cell types predicted to contribute <3% of a spot were considered to be 0 (see *spatial_transcriptomics/CD4-Analysis/CD4-deconvolution.Rmd*). Lastly, for epithelial cells we followed the same approach to the one described above for CD4 T cells (see *spatial_transcriptomics/epithelium_integration/epithelium-deconvolution.Rmd*).

#### ST: Gene expression denoising

Due to the sparsity nature of the data, we denoised the expression of genes of interest using MAGIC, Rmagic v2.0.3 package (van Dijk et al., 2018) to gain a better understanding of the spatial distribution of their expression. We performed MAGIC denoising for gene sets of interest related to CD4 T cells, plasma cells, myeloid cells and Follicular Dendritic cells. MAGIC was run for each slice independently to avoid contaminating expression signal between them. The *knn* parameter was set to a conservative 2 to avoid over-diffusion, furthermore *t* was set to “auto” to determine the extent of diffusion according to the Procrustes disparity. The remaining parameters were kept with their default setting.

#### ST: Spatial trajectory analysis

For Plasma cells, we carried out spatial trajectory analysis using SPATA2 v0.1.0 (Kueckelhaus et al., 2020) to visualize the differentiation trajectory from the germinal center light zone to dark zone to Plasma cell zone. We used the *createTrajectories* function to manually define spatial trajectories across germinal centers to plasma cell rich zones to recapitulate their migration pattern. This enabled us to visualize smoothed gene expression throughout the trajectory using *plotTrajectoryHeatmap* function with smooth_span equal to 0.5.

#### ST: Gene signatures

Gene signatures for Plasma Cells were computed using the package UCell v1.2.0 (Andreatta and Carmona, 2021). To extract relevant marker genes from each cell type, we ran Seurat’s FindAllMarkers on subsetted data containing only Plasma Cells to capture differentially expressed genes between them. For each cell type, we removed ribosomal and mitochondrial genes and filtered out those genes expressed in >25% of *other* cells (keeping those with pct.2 < 0.25). We then ranked them in decreasing order by their avg_log2FC and selected the top 25 for each cell type. Lastly, we ran *AddModuleScore_UCell* to compute each module’s score for each spot on the visium slides.

#### SLOcatoR: Label and coordinate transfer across modalities

To transfer cell types labels and UMAP coordinates across data modalities, we used the following approach:

1. Define the cell types annotation and UMAP coordinates using the transcriptomic data obtained from scRNA-seq and Multiome, as explained above.
2. For scATAC-seq data, we used the labeled data from Multiome; and for CITE-seq we used the labeled data from scRNA-seq. Before integration, we find assay-specific features and their integration as defined above to mitigate technology-specific biases between scRNA-seq and CITE-seq.
3. For both scRNA-seq/CITE-seq (gene expression) and Multiome/scATAC-seq (chromatin accessibility), we integrate them with Harmony, as explained above.
4. We define the success by assessing the degree of integration in the UMAP obtained from the harmony-corrected principal components.
5. We transfer the label from Multiome to scATAC-seq and from scRNA-seq to CITE-seq using a K-nearest neighbors (KNN) classifier, as implemented in the knn function of the class v7.3-19 package (default k = 5, optimized in the case of Multiome, see above).
6. We transfer the UMAP coordinates from Multiome to scATAC-seq and from scRNA-seq to CITE-seq using KNN regression, as implemented in the knnreg function of the caret v6.0-90 package (k = 5).

This workflow has been implemented in the SLOcatoR package (https://github.com/massonix/SLOcatoR) to connect data modalities and annotate unseen transcriptomes and chromatin accessibility profiles from SLO.

#### Data visualization

Different visualization strategies were applied throughout the study to ensure the correct capture and interpretation of the data. Because interleukin genes are expressed at low levels, we used the Nebulosa v1.5.0 R package (Alquicira-Hernandez and Powell, 2021) to recover signals. Similarly, we observed that certain ADT in the CITE-seq data were expressed at low levels. To visualize subtle signals in the UMAP plots, we set the order parameter to “TRUE” in the *FeaturePlot* function of Seurat, which plots cells in order of expression. This approach was applied for proteins CD103, CD54, CD161 and CD56 in Figure 3. Likewise, we applied the same parameter to show TF activity in UMAPs throughout the figures. On the other hand, we observed several ADT that had high levels of background noise. For these, we excluded the first and the last percentile when projecting their expression in the UMAPs (Seurat: FeaturePlot, min.cutoff = “q1”/"q5", max.cutoff = “q99”/"q95"). This was applied for all CITE-seq UMAPs in Figure 2C, and for scATAC-seq UMAPs. To represent heatmaps shown throughout the figures, we generated pseudo-bulk expression profiles for each cluster with the *AverageExpression* function (slot = data) from Seurat. Subsequently, each row is scaled from 0 to 1 and, finally, the resulting matrix is visualized with *pheatmap*2 function from the pheatmap2 R package.

#### HCATonsilData

HCATonsilData is a BioConductor data package developed following the vignette available here: <http://contributions.bioconductor.org/non-software.html>. Briefly, the deposited Seurat objects available in Zenodo were downloaded using zenodo_get. Subsequently, we saved the independent data slots as separate H5File:HDF5 or RDS files, which we later uploaded and stored in a provided Bioconductor Microsoft [Azure Data Lakes](https://azure.microsoft.com/en-us/services/data-lake-analytics). HCATonsilData uses ExperimentHub to query and download those data slots, which are then assembled and returned to the user as a SingleCellExperiment object. Finally, HCATonsilData implements a “updateAnnotation” function that allows users to propose new cell type/state annotations using GitHub issues.
